# Freshwater biodiversity is not adequately addressed by the current protected areas of the Caribbean biodiversity hotspot

**DOI:** 10.64898/2026.03.16.712036

**Authors:** Yusdiel Torres-Cambas, Yander L. Diez, Yoandri S. Megna, Juan Carlos Salazar-Salina, Sami Domisch

**Affiliations:** Leibniz Institute of Freshwater Ecology and Inland Fisheries, Department of Community and Ecosystem Ecology, Müggelseedamm 310, D-12489 Berlin, Germany; Smithsonian Marine Station at Fort Pierce, 701 Seaway Drive, Ft. Pierce, Florida 34949, United States; Universidad Nacional Mayor de San Marcos, Facultad de Ciencias Biológicas, Calle Germán Amézaga 375, Ciudad Universitaria, Apartado Postal 11-0058, Lima 11, Perú; Departamento de Biología y Geografía, Facultad de Ciencias Naturales y Exactas, Universidad de Oriente, Santiago de Cuba, 90500, Cuba

**Keywords:** freshwater biodiversity, Kunming-Montreal Global Biodiversity Framework, post-2020 global biodiversity framework, protected areas, spatial conservation prioritization

## Abstract

**Aim:** Freshwater species face significant challenges from direct and indirect anthropogenic impacts, leading to a global decline in freshwater biodiversity. Protected areas are a key tool for conservation, but their effectiveness in covering freshwater biodiversity remains uncertain. This study assesses the protection coverage of freshwater macroinvertebrates, vertebrates, and macrophytes in Cuba against the 30% conservation targets set by the Convention on Biological Diversity.

**Location:** Cuban biodiversity hotspot, including freshwater ecosystems across the Cuban archipelago.

**Methods:** We compiled occurrence data for 304 freshwater macroinvertebrates, 31 fish, and 24 macrophyte species across the Cuban archipelago and modelled species distributions at the sub-catchment scale using Random Forest and Maxent. Models were fitted for species with at least 10 occurrence records, and the resulting estimates were combined into conservative ensemble suitability estimates. We then performed spatial conservation prioritization using habitat suitability as feature amounts. For species for which model fitting was not feasible, feature amounts were derived by treating sub-catchments with occurrence records as suitable. Spatial conservation prioritization was conducted under a 30% conservation target for each species and two prioritization designs: (i) lock-in, in which planning units overlapping the Cuban National System of Protected Areas (SNAP) were forced into the solution, and (ii) free-choice, in which all planning units were available for selection. We compared these prioritization solutions with SNAP in terms of achieving 30% coverage of suitable habitat for each species and their contribution to hydrological connectivity.

**Results:** SNAP left major freshwater representation gaps across Cuba, with the strongest shortfalls in Decapoda, Hemiptera, Coleoptera, and Ephemeroptera, whereas Odonata and vertebrates were comparatively better represented. Endemic species were also poorly covered, and many were entirely absent from SNAP. Spatial prioritization solutions showed that the 30% target was achievable for all species, but only through substantial expansion or redesign of the current network. The lock-in solution required a larger and more costly network but achieved higher median species coverage, whereas the free-choice solution met the same target more efficiently with less area and lower cost. Both prioritization scenarios also improved hydrological network structure relative to SNAP, producing larger connected freshwater clusters and identifying connector units that linked previously disconnected protected areas. Patterns of planning-unit importance differed among taxonomic groups, indicating that priority areas for one group do not necessarily capture the conservation needs of others.

**Main conclusions:** Overall, our results show that the current protected areas of the Caribbean biodiversity hotspot do not adequately address freshwater biodiversity. SNAP leaves major representation gaps, with 217 of 359 species failing to meet the 30% target, particularly among invertebrates and endemic species, and provides neither an efficient nor well-connected basis for meeting ambitious freshwater conservation goals. The strong taxonomic variation in both protection gaps and planning-unit importance indicates that priority areas for one group do not necessarily represent freshwater biodiversity as a whole, underscoring the need for explicitly multi-taxon planning. Nevertheless, substantial improvement is possible: expanding around the current network would enhance representation and connectivity, although requiring an approximately 70% increase in area, whereas a less constrained redesign could meet the same targets more efficiently. The most effective long-term strategy may therefore be to retain and strengthen the parts of SNAP that already contribute to freshwater conservation, while strategically expanding and reconfiguring the network based on freshwater-specific priorities.

## 1. Introduction

The current detrimental anthropogenic impacts on nature, particularly freshwater biodiversity, are unprecedented in the Earth’s geological history (Flitcroft et al., 2019; Waters & Turner, 2022). Despite being recognized as highly biodiverse and essential providers of ecosystem services to human society, freshwater ecosystems are experiencing a rapid global decline that exceeds rates observed in marine and terrestrial ecosystems (Garcia-Moreno et al., 2014; Lynch et al., 2023; Sayer et al., 2025; WWF, 2020). As we progress through the 21st century and encounter new threats to freshwater ecosystems, it becomes increasingly urgent to take immediate action to halt and reverse this crisis (Reid et al., 2019; Tickner et al., 2020).

Protected areas play a critical role in broader efforts to address freshwater biodiversity loss and protect the essential services provided by freshwater ecosystems (Acreman et al., 2019; Donald et al., 2019; Watson et al., 2014). To maximize the potential of protected areas to maintain ecological processes, they require a catchment-wide perspective that takes into account the entire range of freshwater ecosystems within a given catchment, from headwaters to downstream reaches (Abell et al., 2016). Moreover, protected areas need to be strategically placed to maximize their conservation value and to address the complementarity and irreplaceability of species and their functions (Margules & Pressey, 2000; Rapacciuolo et al., 2018). Furthermore, spatial conservation prioritization must consider connections across the river network (longitudinal connectivity) and between river channels and adjacent habitats, such as lakes or wetlands (lateral connectivity) (Moilanen et al., 2009; Reis et al., 2019). Longitudinal connectivity is crucial for aquatic species’ dispersal and population dynamics, especially fish (O’Mara et al., 2021). Lateral connectivity, on the other hand, supports nutrient exchange, facilitates access to important breeding and feeding grounds for various species or contributes to maintaining macrophyte community diversity along rivers (Couto et al., 2018; Covino, 2017; Keruzoré et al., 2013). Addressing connectivity is particularly crucial in the face of climate change, as freshwater ecosystems experiment altered hydrological patterns, increased temperatures, and changes in precipitation patterns that lead to droughts and flooding events (Nickus et al., 2010; Verdonschot et al., 2010). Designing protected areas that account for connectivity ensures that species can move and adapt to changing environmental conditions, enhancing their chances of survival and resilience (Juffe-Bignoli et al., 2016).

The Kunming-Montreal Global Biodiversity Framework sets ambitious and transformative biodiversity conservation goals for the current decade (2021-2030) and beyond (Hughes, 2023). Among the proposed targets, this new framework urges to preserve by 2030 at least 30 % of terrestrial, marine and coastal areas, and inland waters (Hughes, 2023). Notably, this is the first time that inland waters are explicitly acknowledged as a distinct realm by the Convention on Biological Diversity (CBD), which has the high potential to benefit freshwater biodiversity (Cooke et al., 2023). However, given the legacy of the neglect of freshwater biodiversity (Abell et al., 2016), we have to acknowledge that existing protected areas may fall far short of adequately safeguarding it. This calls for testing the new post-2020 conservation strategy where at least 30% of freshwater species distribution is protected. In this regard, biodiversity hotspots are particularly importance for meeting the goals of the CBD.

One of those hotspots, the Caribbean islands, hosts an exceptionally high level of species diversity and it is also under significant threat of habitat loss and degradation (Mittermeier et al., 2011). The Cuban archipelago, including the main island of Cuba and its surrounding smaller islands, contributes significantly to the overall biodiversity and conservation value of the Caribbean hotspot. To date, approximately 800 freshwater species have been found across the Cuban archipelago, including 92 macrophytes, 46 fish, 35 mayflies (Ephemeroptera), 88 dragonflies and damselflies (Odonata), and 91 caddisflies (Trichoptera) among the insects (Aguilar, Betancourt, et al., 2009; Aguilar, Cabrera, et al., 2009; Mancina & Cruz Flores, 2017; Torres-Cambas et al., 2023; Urquiola, Vega, et al., 2000; Urquiola, Aguilar, et al., 2000; Urquiola, Vega, et al., 2009; Urquiola, Novo, et al., 2009; Urquiola & Betancourt, 2000; Urquiola & Novo, 2000; Urquiola & Pérez, 2009a, 2009b). Cuba’s high freshwater species richness is accompanied by a high degree of endemism, with at least 202 species found only in the Cuban archipelago (Torres-Cambas et al., 2023).

Cuba is a small (110,977 *km*^2^) yet highly populated (102.3 inhabitants/*km*^2^) tropical insular country, where freshwater is scarce, and consequently, freshwater ecosystems are under high anthropogenic pressure. To satisfy the growing demand for freshwater, the course and flow of major rivers in Cuba have been altered during the last sixty years given the construction of reservoirs or channels to transfer water from one river to another (Veloso & Pérez, 2013). Additional pressures expected in the near future include climate change, associated alterations in hydrological regimes, and the salinization of freshwater habitats resulting from sea-level rise, all of which are predicted to have profound impacts on Cuban freshwater ecosystems. These climate-change effects, together with the loss and fragmentation of freshwater habitats, are currently considered to be among the main threats to Cuban biodiversity (Mancina & Cruz Flores, 2017).

Cuban biodiversity is legally protected by the National System of Protected Areas (hereafter referred to as’SNAP’ - Sistema Nacional de Areas Protegidas). Terrestrial and inland water protected areas in the SNAP represent 16% of the national surface. However, evidence suggest that freshwater ecosystems are not effectively protected or represented in the SNAP, since Cuban rivers are among the ones receiving the least integral protection (i.e., local and upstream catchment protection) in the world (Abell et al., 2016; CNAP, 2013). Furthermore, the actual representativeness of freshwater biodiversity in the SNAP is poorly known. To date, gap analyses have been restricted to fish, diving beetles (Coleoptera: Dytiscidae) and threatened species of damselflies (Odonata: Zygoptera) (CNAP, 2013; Megna et al., 2021; Torres-Cambas et al., 2015). This narrow taxonomic spectrum therefore does not serve as an indicator for the representativeness of the freshwater biodiversity in the SNAP, or as a surrogate to derive the protection coverage of other taxonomic groups (Stewart et al., 2018).

Here we employ a spatial conservation prioritization approach (Moilanen et al., 2009) across the Cuban archipelago using freshwater species distributions as target features and ask (i) how well does the SNAP provide protection for various freshwater species groups, and (ii) where would possible conservation gaps occur? As the current focus of protected area networks is on charismatic species or landscapes, which bias conservation efforts, we tried to cover part of the “hidden biodiversity” (Delso et al., 2021) by including as many macroinvertebrates as possible in our analysis, in addition to more conspicuous groups such as vertebrates and macrophytes. We provide conservation planning solutions under the post-2020 global biodiversity framework targets (30%) for these species while addressing the connectivity, and discuss the results within the broader context of freshwater representativeness in protected areas in general.

## 2. Methods

### 2.1. Spatial modelling units

We used sub-catchments as spatial modelling units for species distribution modelling, as they capture both freshwater habitats and their surrounding terrestrial context, which can strongly influence freshwater biodiversity patterns (Hermoso et al., 2011). Spatial units were derived from the Hydrography90m river network (Amatulli et al., 2022), a high-resolution hydrographic dataset based on the MERIT Hydro digital elevation model (Yamazaki et al., 2014). These sub-catchments provide a spatially explicit representation of riverine drainage units suitable for modelling freshwater species distributions.

To restrict the modelling domain to areas relevant to freshwater habitats, we further subset the Hydrography90m sub-catchments using freshwater features from OpenStreetMap (OpenStreetMap contributors, 2025), including streams, rivers, reservoirs, and wetlands. Only sub-catchments overlapping with mapped freshwater areas across Cuba were retained for analysis. From an initial set of 584,637 Hydrography90m sub-catchments, this filtering resulted in 130,332 sub-catchments (median area = 0.14 km^2^, interquartile range = 0.08-0.25 km^2^) used as spatial modelling units across the Cuban archipelago.

### 2.2. Species data

For our analyses, we considered occurrences for 304 freshwater macroinvertebrates, 31 fish and 24 macrophyte species from 1980 to 2025 (see Table S1.1 Appendix S1 in Supporting Information and dataset at https://doi.org/10.5061/dryad.hqbzkh1zn). We obtained freshwater macroinvertebrate occurrences from a freshwater macroinvertebrates database specific to Cuba (Torres-Cambas et al., 2023) and macrophyte and fish data from GBIF (GBIF.org User, 2026a, 2026b).

For the GBIF data, we discarded occurrence records without coordinates or year information. We also removed records located within a 1 km buffer around country centroids, capital centroids, herbaria, and museum centroids (Zizka et al., 2019). To match the temporal coverage of the climate variables used, we retained only records from 1980 onwards. Occurrences were then snapped to the nearest sub-catchment overlapping a water body, and records with a snapping distance greater than 500 m were discarded. Finally, occurrence records were aggregated by sub-catchment. Species distribution models were fitted only for species with 10 or more occurrence records (n = 106, see Tables S1.2 and S1.3, Appendix S1 in Supporting Information). For species for which model fitting was not possible because low number of records (n = 253) or model predictions were discarded due to low model performance (n = 4), Hydrography90m sub-catchments with known occurrence records were treated as suitable.

### 2.3. Defining the accessible area

To avoid projecting species distributions into areas that are biogeographically unlikely to have been accessible to each species, we constrained the modelling extent by defining taxon-specific accessible areas based on known distribution ranges within Cuba. The accessible area for each species was defined using the biogeographic regions of Cuba, Western, Central, Central-eastern, and Eastern, following Núñez (1989). Each species was assigned to one or more regions based on its known distribution range, as documented in the taxonomic and distributional literature. For macroinvertebrates, regional assignments were based on Megna et al. (2021), Naranjo-López et al. (2019), Naranjo-López & González-Lazo (2005), Naranjo et al. (2010), Perera & Valderrama (2010) and Trapero-Quintana et al. (2018). For freshwater fishes, assignments followed Rodríguez-Machado & Ponce de León (2017). For macrophytes, regional assignments were based on distributional information reported in the relevant family-level fascicles of Flora de Cuba (Aguilar, Betancourt, et al., 2009; Aguilar, Cabrera, et al., 2009; Urquiola, Aguilar, et al., 2000; Urquiola, Novo, et al., 2009; Urquiola, Vega, et al., 2000, 2009; Urquiola & Betancourt, 2000; Urquiola & Novo, 2000; Urquiola & Pérez, 2009a, 2009b). These accessible areas were then used to restrict model calibration and prediction to regions considered reachable for each species.

### 2.4. Pseudoabsences

As our occurrence dataset contained only presence records, we generated pseudo-absences using a three-step procedure (Senay et al., 2013). First, we restricted the candidate pool to freshwater sub-catchments located within the regions considered accessible to each species and excluded sub-catchments with known presences. Second, we used a one-class support vector machine (Schölkopf et al., 2001) to characterize the environmental domain of the presence sub-catchments and removed candidate sub-catchments predicted to fall within this domain, retaining environmentally dissimilar candidates as potential pseudo-absences. Finally, pseudo-absences were selected from the remaining candidates using balanced k-means subsampling in environmental space, generating five independent replicates per species. For each replicate within a species, the number of pseudo-absences was set to twice the number of presences of the species.

### 2.5. Environmental data

We used environmental predictors from the freshwater-specific Environment90m dataset (Garcia Marquez et al., 2026), which provides environmental variables aggregated to the sub-catchments of the Hydrography90m river network (Amatulli et al., 2022). These predictors were selected to represent the main climatic, topographic, hydrological, and land-cover conditions of the study area. The variables included topographic and river-network descriptors, namely flow accumulation, downstream distance to the basin outlet, gradient, stream power index, and compound topographic index, as well as climatic predictors representing mean annual temperature, annual precipitation, and precipitation seasonality for the observed 1981–2010 climatological period.

Land-cover information was represented through a human-pressure index derived from three land-cover classes available annually from 1992 to 2020 and summarized at the sub-catchment level: rainfed cropland, mosaic cropland and natural vegetation, and urban areas. This index was calculated as the sum of the proportional cover of these classes within each sub-catchment, providing an aggregate measure of human-modified landscape conditions relevant to freshwater ecosystems.

Because land cover varied through time, we assigned the human-pressure index to occurrence records using a centred rolling mean based on the year of each occurrence. For occurrences dated between 1993 and 2019, values were calculated as the mean of the previous, current, and following year. For occurrences in the first and last years of the time series, where a full three-year window was not available, we used two-year averages: mean (1992, 1993) for 1992 and mean (2019, 2020) for 2020. Occurrences dated before 1992 were assigned the 1992 value, while occurrences dated after 2020 were assigned the 2020 value, corresponding to the temporal limits of the available land-cover time series.

To reduce multicollinearity among predictors, we assessed pairwise correlations and variance inflation factors (VIFs) before model fitting. Highly correlated variables were identified using a correlation threshold (|r| > 0.7). To refine our predictor set, we then used the function *vifstep()* from the *usdm* package (Naimi, 2023), which sequentially removes variables with high VIF values until all remaining predictors are below a given threshold (here we used 10 as the standard limit).

### 2.6. Modelling freshwater species distributions

We used species occurrence and pseudo-absence data, together with the environmental predictors, to build an ensemble species distribution model. We employed two modelling approaches, maximum entropy (Maxent, Phillips et al., 2006) and random forest (RF, Zhang et al., 2019). Maxent and RF were combined because they provide complementary spatial representations of habitat suitability. Random forest can capture complex nonlinear relationships and may produce more localized suitability signals, or suitability maps with higher discrimination and greater spatial heterogeneity (Zhao et al., 2022). Maxent uses regularized maximum-entropy estimation to produce constrained suitability surfaces based on environmental conditions at occurrence sites (Elith et al., 2011; Phillips et al., 2006; Phillips & Dudík, 2008). Combining both algorithms therefore allowed the ensemble to integrate localized and generalized suitability signals, with their disagreement contributing to the uncertainty estimate (Araújo & New, 2007; Thuiller et al., 2009).

For each species, presences were combined with species-specific pseudoabsences, and models were calibrated only within the species’ accessible area, defined from the biogeographic regions occupied by the species (Barve et al., 2012; Núñez, 1989). The number of cross-validation folds was adapted to the number of unique presence records: species with fewer than 15 presences were evaluated using two folds, species with 15–29 presences using three folds, and species with 30 or more presences using five folds; in all cases, cross-validation was repeated five times (Tables S1.2 and S1.3, Appendix S1 in Supporting Information). For RF, hyperparameters *mtry*, *min.node.size*, and *sample.fraction* were tuned for each species optimizing model performance using AUC (Zhao et al., 2022). Final RF models were then fitted for each repeat–fold combination using the selected values for hyperparameters. For Maxent, hyperparameters were tuned by testing combinations of feature classes (l, lq) and regularization multipliers, and final models were fitted using the selected feature class and regularization multiplier (Radosavljevic & Anderson, 2014). For both algorithms, model performance was evaluated on withheld test data using AUC, standardised True Skill Statistics (sTSS) (Barbosa, 2015), sensitivity, specificity, omission, and commission, and model-specific thresholds were estimated using the threshold that minimized the difference between sensitivity and specificity (Freeman & Moisen, 2008; Jiménez-Valverde & Lobo, 2007) (Tables S1.2 and S1.3, Appendix S1 in Supporting Information). Predictions were then projected to all sub-catchments within the species-specific accessible area, and predictions outside the accessible area were set to zero. Finally, predictions from all cross-validation models in both algorithms were retained only when model AUC ≥ 0.5 and sTSS ≥ 0.5 (Tables S1.2 and S1.3, Appendix S1 in Supporting Information).

To use model predictions as input for the spatial conservation prioritization analysis, we transformed continuous habitat suitability values by applying a threshold that minimized the difference between sensitivity and specificity (Freeman & Moisen, 2008; Jiménez-Valverde & Lobo, 2007). For each algorithm, predicted suitability values were first thresholded at the level of individual cross-validation models using the corresponding model-specific threshold. Values below the threshold were set to zero, whereas values above the threshold retained their continuous suitability values (Domisch et al., 2019). Thresholded predictions were then averaged across cross-validation runs to obtain algorithm-specific ensemble suitability values for RF and Maxent. Then, species-level weighted mean suitability values were produced by combining the RF and Maxent ensemble predictions using performance-based weights derived from mean sTSS values. Finally, we calculated a conservative suitability value for conservation prioritization that incorporated uncertainty by subtracting a combined uncertainty estimate from the weighted mean suitability. This uncertainty estimate included both within-algorithm uncertainty and disagreement between RF and Maxent predictions. When only one algorithm met the predefined performance criteria (AUC_mean ≥ 0.70 and sTSS_mean ≥ 0.70), the final species-level suitability value was based on that algorithm alone, and the conservative suitability value was calculated by subtracting the corresponding within-algorithm ensemble standard deviation from the algorithm-specific ensemble mean.

### 2.7. Sensitivity of species distribution models to occurrence age

To address potential temporal mismatch between occurrence records and environmental predictors, we evaluated the sensitivity of SDM results to record age using two RF model scenarios: (1) all records from 1980–2025 and (2) records restricted to 1992–2020 to match land-cover availability. We then compared model performance, variable importance, and spatial predictions between scenarios.

Differences in model performance between scenarios were assessed using paired comparisons across identical cross-validation folds. We used Wilcoxon signed-rank tests on fold-level evaluation metrics and additionally reported mean paired differences and associated variability.

To assess whether predictor importance changed between temporal scenarios, we extracted variable-importance values from all fitted models for each species and scenario. For each predictor, we calculated mean importance across model runs within each species and scenario. We then compared scenarios by examining differences in mean importance for each predictor and by calculating the Spearman rank correlation between the full variable-importance profiles of the two scenarios for each species. This allowed us to evaluate both changes in the absolute contribution of predictors and the consistency of predictor ranking between scenarios.

Spatial predictions were further used to compare spatial suitability patterns between temporal scenarios. Spatial agreement was quantified using Spearman correlation of suitability values across sub-catchments, and mean absolute differences in predicted suitability. In addition, to assess consistency in the identification of the most suitable spatial units, we calculated the overlap between scenarios in the top 10% and top 20% highest-ranked sub-catchments. Together, these metrics captured both overall agreement in spatial predictions and agreement in the sub-catchments most relevant for subsequent spatial prioritization.

### 2.8. Planning units

For the spatial conservation prioritization analysis, we used planning units that were larger than the original spatial modelling units used in the species distribution models. This decision was made to reduce computational complexity and to conduct prioritization at a spatial scale more suitable for conservation planning and management, while still preserving hydrological meaning.

To generate larger planning units, we followed the same workflow and used the same digital elevation model used to derive the Hydrography90m stream network and sub-catchments, but applied a larger upstream contributing area threshold to delineate larger sub-catchments (Amatulli et al., 2022). We first extracted a river network and the corresponding sub-catchments from the MERIT Hydro digital elevation model (DEM; 3-arc sec resolution, approximately 90 m at the Equator; Yamazaki et al., 2014) using GRASS GIS v.7.8.5. Stream channels were initiated using an upstream contributing area threshold of 100 DEM cells, corresponding to approximately 0.81 km². The resulting sub-catchments ranged from 0.008 to 33.55 km², with a median area of 1.28 km² and an interquartile range of 0.64–2.21 km². We then filtered these units by retaining only those sub-catchments that overlapped mapped freshwater features across Cuba as we did with spatial modelling units. From an initial set of 69,528 sub-catchments, this filtering resulted in 29,417 planning units across the Cuban archipelago.

To transfer SDMs predictions to the larger planning units, Hydrography90m sub-catchments were assigned to MERIT Hydro-derived planning units using spatial overlay. For each planning unit, species-specific feature amounts were then calculated as the sum of predicted suitability values across all Hydrography90m sub-catchments contained within it, weighted by sub-catchment area.

### 2.9. Spatial conservation prioritization

We formulated and solved the minimum-set coverage conservation prioritization problems with the R package *prioritizr* and the exact integer linear programming solver Gurobi to an optimality gap of 5 % (Beyer et al., 2016; Gurobi Optimization, LLC, 2026; Hanson et al., 2025; Moilanen et al., 2009). In this kind of prioritization problem, the objective is to find a solution that, at a minimum cost, achieves a particular conservation target. In our analysis, the objective was to identify a potential network of protected areas encompassing 30 % of the suitable habitat for each of the 359 analysed species. We implemented two prioritization designs. In the lock-in design, planning units overlapping the current Cuban protected areas (SNAP) were locked in, meaning they were pre-selected and the algorithm could only add additional planning units to meet species-specific targets. In the free-choice design all planning units, including those inside protected areas, were available for selection. The SNAP layer was sourced from the World Database on Protected Areas (UNEP-WCMC, 2025). A planning unit was considered lock-in only when at least 50 % of its area was covered by protected areas. We refer to the total lock-in area as the “inland water area of the SNAP”.

The spatial conservation planning requires costs as the counterpart of the conservation targets. In our case, we used the Human Footprint Index (HFI) for 2020 as an indicator of anthropogenic alteration to represent the cost of the solution (Gassert et al., 2023). Higher HFI values correspond to more human-altered planning units and contribute to higher solution costs. Hence, minimizing the cost of the solution would be equivalent to prioritizing those planning units that are less anthropogenically impacted, as opposed to planning units that show a high anthropogenic impact. The rationale behind considering highly human-altered areas as less suitable for conservation is that they would need additional conservation actions to ensure that biodiversity is adequately addressed (Hermoso et al., 2015). We aggregated the original HFI layer to the planning units.

To promote longitudinal connectivity in the solutions, we included a connectivity penalty term in our spatial conservation planning problems. This penalty is designed to account for the importance of promoting connectivity between planning units. The penalty’s magnitude is determined by connectivity scores that quantify the strength of connectivity between planning units. Higher scores indicate a stronger level of connectivity. We calculated these scores based on the inverse distance between planning units located within the same drainage basin (Hermoso et al., 2012). Higher penalty values yield solutions with higher connectivity, meaning that the selected planning units are spatially clumped to a greater extent. However, this also comes at a higher cost because planning units that show a higher HFI must be included in the solution to achieve higher connectivity. Therefore, to achieve a balance between cost and the overall connectivity of the solution, we generated a set of 29 candidate prioritizations at an optimality gap of 10 % that involved different penalty values ranging from 0.001 to 0.068. Then, we plotted the relationship between the overall cost and overall connectivity for these candidate solutions. Overall cost was calculated as the sum of planning-unit cost values weighted by their selection status, using the *eval_cost_summary()* function in the *prioritizr* package (Ball et al., 2009; Hanson et al., 2025). Overall connectivity was quantified as the sum of pairwise connectivity scores weighted by the value of the planning units in the solution (Ball et al., 2009; Hanson et al., 2023) using the *eval_connectivity_summary()* function (Ball et al., 2009; Hanson et al., 2025). Finally, from the cost-connectivity plot we identified the “elbow” of the curve in the R package *pathviewr* (Baliga et al., 2021) and chose the connectivity penalty at this point (0.031) to produce the final prioritizations at an optimality gap of 5 % following the two designs mentioned above (Hermoso et al., 2011; Stewart & Possingham, 2005).

After solving the conservation problems, we evaluated how well each species was represented by the SNAP and by the prioritization scenarios, and identified which planning units were most important for meeting the proposed targets. We calculated planning-unit importance scores following Ferrier et al. (2000), which quantify the contribution of each planning unit to achieving conservation targets. In this framework, highly important or irreplaceable planning units are those that make a large contribution to target achievement and for which there are relatively few spatial alternatives capable of providing the same contribution. This approach also allows us to examine feature-level importance within each planning unit, thereby revealing the species-specific contributions that underpin the overall importance of a given unit. We used these feature-level values to identify which taxonomic groups were responsible for high planning-unit importance, and to assess whether spatial patterns of irreplaceability were shared among groups or driven by particular components of freshwater biodiversity.

To evaluate spatial congruence in planning-unit importance among taxonomic groups, we calculated pairwise Spearman rank correlations between group-specific irreplaceability values. Correlations were calculated using planning units with positive irreplaceability for at least one of the two compared groups, thereby reducing the influence of shared zero values. Positive correlations were interpreted as evidence that planning units important for one taxonomic group also tended to be important for another group, whereas weak or negative correlations indicated spatial mismatch or complementarity. In addition, we assessed the overlap among the highest-importance planning units using Jaccard similarity (Chung et al., 2019). For each taxonomic group, we identified high-irreplaceability planning units as those within the upper 5% of positive group-specific irreplaceability values, and then calculated pairwise Jaccard similarity among these taxon-specific priority sets to quantify the degree to which the highest-importance planning units were shared among groups.

In addition to representation and importance, we assessed whether the resulting networks improved spatial connectivity along the stream system. We first quantified the connected-component structure of SNAP and prioritizations network. For each scenario, we built an undirected stream-network graph in which selected planning units were represented as nodes and direct stream-network links as edges. We then calculated the number of connected components, the size of the largest component, and the proportion of selected planning units contained within the largest component. These metrics describe whether each network was fragmented into many small clusters or organized into larger connected networks along the stream system. We also evaluated whether prioritization solutions connected existing protected areas. A pair of protected areas was considered connected when at least one planning unit from each protected area occurred within the same connected component of the stream-network graph.

## 3. Results

### 3.1. Species distribution models

Random Forest results were highly consistent between the full-occurrence scenario (1980–2025) and the predictor-overlap scenario (1992–2020). Across species, differences in mean AUC, sTSS, sensitivity, specificity, omission, and commission were minimal, with average paired differences close to zero for all metrics (dataset at https://doi.org/10.5061/dryad.hqbzkh1zn). More than half of the species showed no difference in mean fold-level performance between scenarios for a given metric, and no species-level differences remained significant after correction for multiple testing. Together, these results indicate that excluding pre-1992 records had negligible effects on overall predictive performance. Predictor importance was broadly stable between scenarios (dataset at https://doi.org/10.5061/dryad.hqbzkh1zn). Variable importance profiles in the full-occurrence scenario and the predictor-overlap scenario were strongly correlated (mean Spearman ρ = 0.962; median = 1.00), and the top-ranked predictor was the same in 94 species (91.2%). Mean differences in importance were small for all predictors, with only slight increases in the full-occurrence scenario for variables such as compound topographic index (cti), accumulation, annual precipitation (bio12), and precipitation seasonality (bio15). Overall, temporal restriction of records had negligible effects on the ranking and relative contribution of predictors.

Spatial predictions were broadly robust to temporal filtering of occurrences. Sub-catchment level suitability predictions from full-occurrence scenario and the predictor-overlap were strongly correlated (mean Spearman ρ = 0.984; median = 1.00) across species, with small absolute differences in suitability and high overlap of the top-ranked sub-catchments (mean overlap = 0.878 for the top 10% and 0.903 for the top 20%). Notably, 51 species showed identical spatial predictions under both scenarios. Given the negligible influence of excluding pre-1992 records, we used RF and Maxent models fitted to the complete occurrence dataset (1980–2025) to generate the ensemble suitability predictions for all subsequent conservation prioritization analyses.

A total of 102 species passed the performance filtering criteria for at least one algorithm (AUC_mean ≥ 0.70 and sTSS_mean ≥ 0.70), and only for these species were SDM-derived suitability values used as feature amounts in the spatial conservation prioritization. Across the 106 species modelled with each algorithm, Random Forest generally achieved higher predictive performance than Maxent, with higher average AUC (0.887 vs 0.834) and sTSS (0.817 vs 0.777), as well as slightly higher mean sensitivity (0.818 vs 0.784) and specificity (0.816 vs 0.770). On average, models were fitted with 23 occurrence records per species (median = 19, range = 10 - 74), and species were typically evaluated across 15 fitted cross-validation models, although this varied with sample size (Tables S1.2 and S1.3, Appendix S1 in Supporting Information).

### 3.2. Representation gaps in freshwater biodiversity under SNAP

Representation of freshwater biodiversity under SNAP was highly uneven across taxonomic groups (Figure 1). The largest representation gaps occurred in Decapoda, Hemiptera, Coleoptera, and Ephemeroptera, where at least three quarters of species remained below the 30% target. Decapoda showed the strongest shortfall, with 81.0% (n = 17) of species below target and a median SNAP coverage of only 4.9%, followed by Hemiptera, where 76.7% (n = 23) of species were below target and the median coverage was 0%. Coleoptera and Ephemeroptera also showed substantial under-representation, with 75.7% (n = 53) and 75.0% (n = 21) of species below target, respectively. Molluscs and macrophytes showed intermediate levels of under-representation, with 63.3% (n = 19) and 54.2% (n = 13) of species below target. Diptera and Trichoptera also had more than half or exactly half of species below target, with a subset of species relatively well represented while others remained poorly covered (Figure 1). Odonata and vertebrates were comparatively better represented, with median SNAP coverages above 30% and the lowest proportions of species below target, 44.3% (n = 31) and 38.7% (n = 12), respectively. Overall, these patterns indicate that SNAP provides relatively better coverage for odonates and vertebrates, but leaves major representation gaps for several freshwater invertebrate groups, particularly decapods, hemipterans, coleopterans, and ephemeropterans.

**Figure 1:**
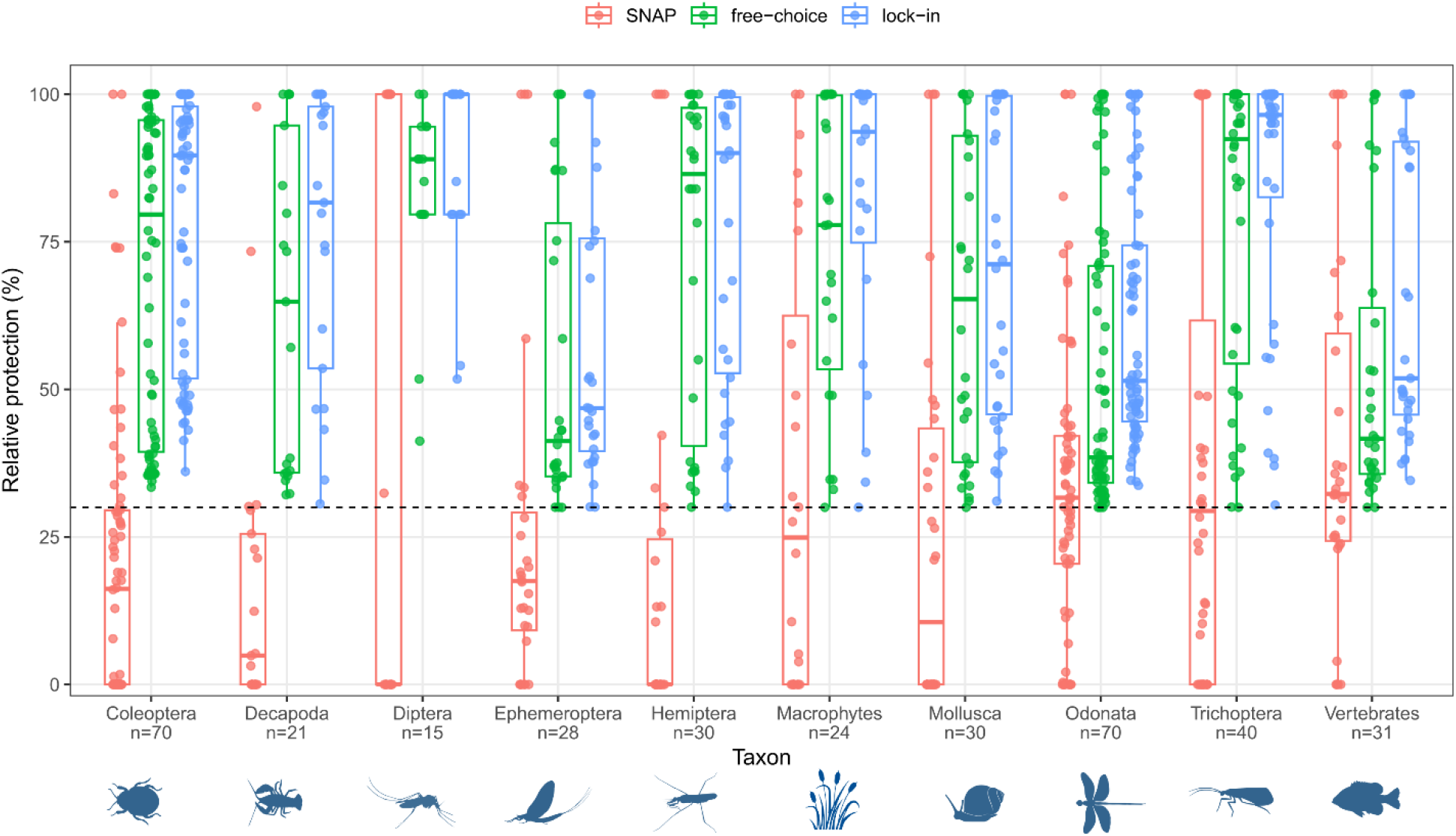
Relative protection of freshwater biodiversity features by taxonomic group under SNAP and the prioritization scenarios. Vascular aquatic plants are grouped as macrophytes. Boxplots show the distribution of species-level relative protection values within each group, with points representing individual species. Horizontal dashed line indicates conservation target threshold of 30%.

SNAP representation was generally low across species of conservation concern. Among Cuban endemic species that are not classified as threatened by the IUCN, 67 of 106 species were represented below the 30% target, equivalent to 63.2% of the group (Table S1.4 Appendix S1 in Supporting Information). In addition, 33 endemic species were completely absent from SNAP, representing 31.1% of endemic-only species (Table S1.4). The median SNAP coverage for this group was 21.3%, indicating that at least half of the endemic species had less than one quarter of their distribution covered by the current protected-area network. A similar pattern was observed for species that were both endemic and threatened. Of the five species in this category (*Hypolestes trinitatis*, *Lucifuga dentata*, *Nandopsis ramsdeni*, *Protoneura caligata*, *Quintana atrizona*), three species were below the 30% target (*H. trinitatis*, *N. ramsdeni*, *P. caligata*) (Table S1.4). However, none of these species was completely absent from SNAP.

Species classified as neither endemic nor threatened also showed substantial representation gaps. In this group, 147 of 247 species were below the 30% target (59.5%), and 85 species were absent from SNAP (34.4%) (Table S1.4). Median coverage was 24.1%, again below the target. This indicates that poor SNAP representation is not restricted to endemic or threatened species, but affects freshwater biodiversity more broadly.

### 3.3. Spatial conservation prioritization solutions

Both prioritization solutions achieved the 30% representation target for all species, but they differed strongly in area and cost (Figure 1, 2). The lock-in solution, which retained all SNAP planning units, required 10,739 planning units covering 18,575.9 km², equivalent to 16.9% of Cuba and 1.7 times the inland-water area currently covered by SNAP. This solution had the highest total cost (1738.6) but also the highest median species coverage (78.2%), indicating that incorporating the existing protected-area network produced a larger and more representative conservation network.

**Figure 2:**
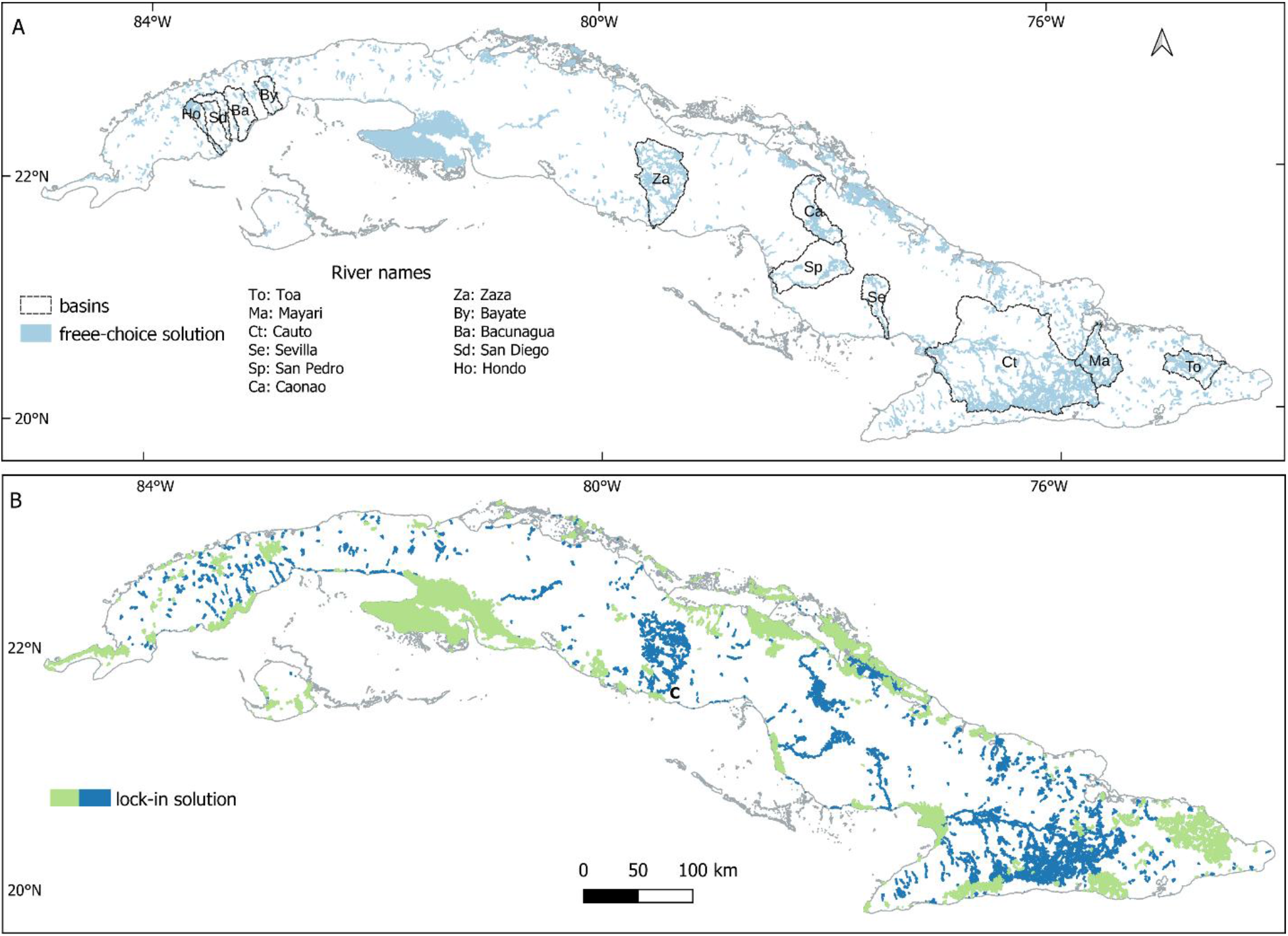
Results of a spatial conservation prioritization analysis in Cuba for the protection of freshwater biodiversity. We used the 30% conservation target of the post-2020 global biodiversity framework as a target for each species in the analysis. A: Free-choice solution. In this scenario, the Cuban System of Protected Areas (SNAP) is not locked in, and the algorithm selects among all planning units to meet species targets. Major river basins across Cuba covered by the solution are highlighted. B: Lock-in solution. In this scenario, the SNAP (green) is pre-selected, and additional planning units (blue) are identified to achieve the required conservation targets and hydrological connectivity.

The same 30% representation target could be achieved with a smaller and less costly network when SNAP was not forced into the solution. The free-choice solution selected 7,009 planning units covering 13,940.3 km² or 12.7% of Cuba, with a lower total cost (1061.8) and median species coverage (64.9%). Only 42.5% of SNAP overlapped with the free-choice solution, suggesting that much of the current protected-area network is not located in the most efficient set of planning units for representing the freshwater species analysed here. Mean cost per selected planning unit was highest in the lock-in scenario (0.162), intermediate in the free-choice scenario (0.151), and lowest in the existing SNAP network (0.141). This indicates that constraining solutions to retain current protected areas reduced cost efficiency relative to the free-choice solution, although the differences were modest.

By comparison, SNAP alone covered 6,840 planning units and 11,159.5 km² of inland-water planning units, representing 10.2% of Cuba, but provided substantially lower median species coverage (23.1%). Overall, these results show that the current protected-area system alone is insufficient to meet freshwater biodiversity targets, while the lock-in solution improves representation by expanding around SNAP, and the free-choice solution achieves targets more efficiently when not constrained by the existing protected-area network.The connected-component analysis revealed marked differences in the spatial structure of selected planning units among SNAP and the prioritization scenarios (Figure 3). In all scenarios, the largest connected component was associated with major freshwater systems, but its location differed among solutions. For both prioritization scenarios, the largest component included the stream network of the Cauto River basin, whereas in the SNAP scenario the largest component was located mainly in the western region of the Zapata Peninsula (Figure 3D-E). Both prioritization scenarios produced a higher number of connected components than SNAP, with the lock-in solution showing the highest number of components, indicating a more fragmented overall network. However, the largest component in both prioritization scenarios was approximately twice the size of the largest component in SNAP. This was also reflected in the largest-component fraction, which was highest in the free-choice solution, followed by the lock-in solution and SNAP. These results indicate that, although the prioritization scenarios selected more spatially dispersed sets of planning units, they also generated larger continuous stream-network clusters than the current protected-area system alone.

**Figure 3.**
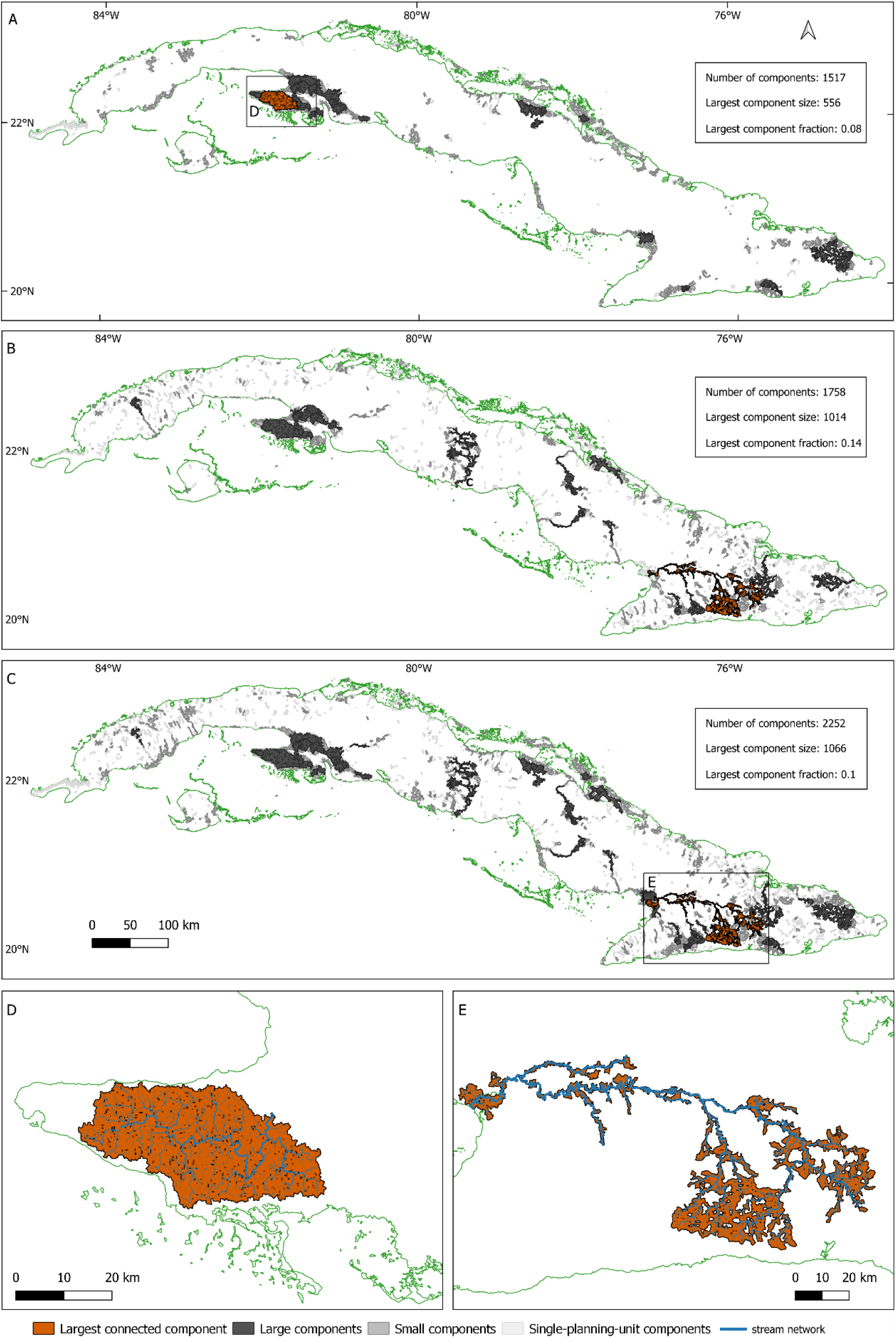
Connected-component structure of SNAP and prioritization networks along the stream system. Selected planning units were represented as nodes in an undirected stream-network graph, with direct upstream–downstream stream links used as edges. Maps show selected planning units dissolved and grouped by connected component for (A) the current Cuban protected-area system, (B) the free-choice prioritization solution, and (C) the lock-in prioritization solution. In each scenario, the largest connected component is highlighted. Single-planning-unit components were classified as isolated fragments, while the remaining multi-unit components were classified by size: large components correspond to those in the upper 95th percentile of component size, and small components include all other multi-unit components. Panels (D) and (E) show the stream network associated with the largest connected component in SNAP and in the prioritization solutions, respectively. For each scenario, the number of connected components, largest component size, and largest component fraction are reported. Larger largest-component sizes and fractions indicate that selected planning units are concentrated in a more continuous stream-network cluster, whereas a higher number of components indicates greater fragmentation.

Under the lock-in solution, several selected planning units outside SNAP contributed to hydrological connectivity among protected areas (Figure 4). These connector units were concentrated mainly along the stream networks of the Cauto and Mayarí river basins in eastern Cuba, where they linked 15 protected areas that were disconnected under SNAP alone (Figure 2, 4). In particular, the selected connectors linked upstream protected areas with protected areas located closer to river estuaries, suggesting that the lock-in solution can strengthen longitudinal connectivity along major freshwater systems while building on the existing protected-area network.

**Figure 4.**
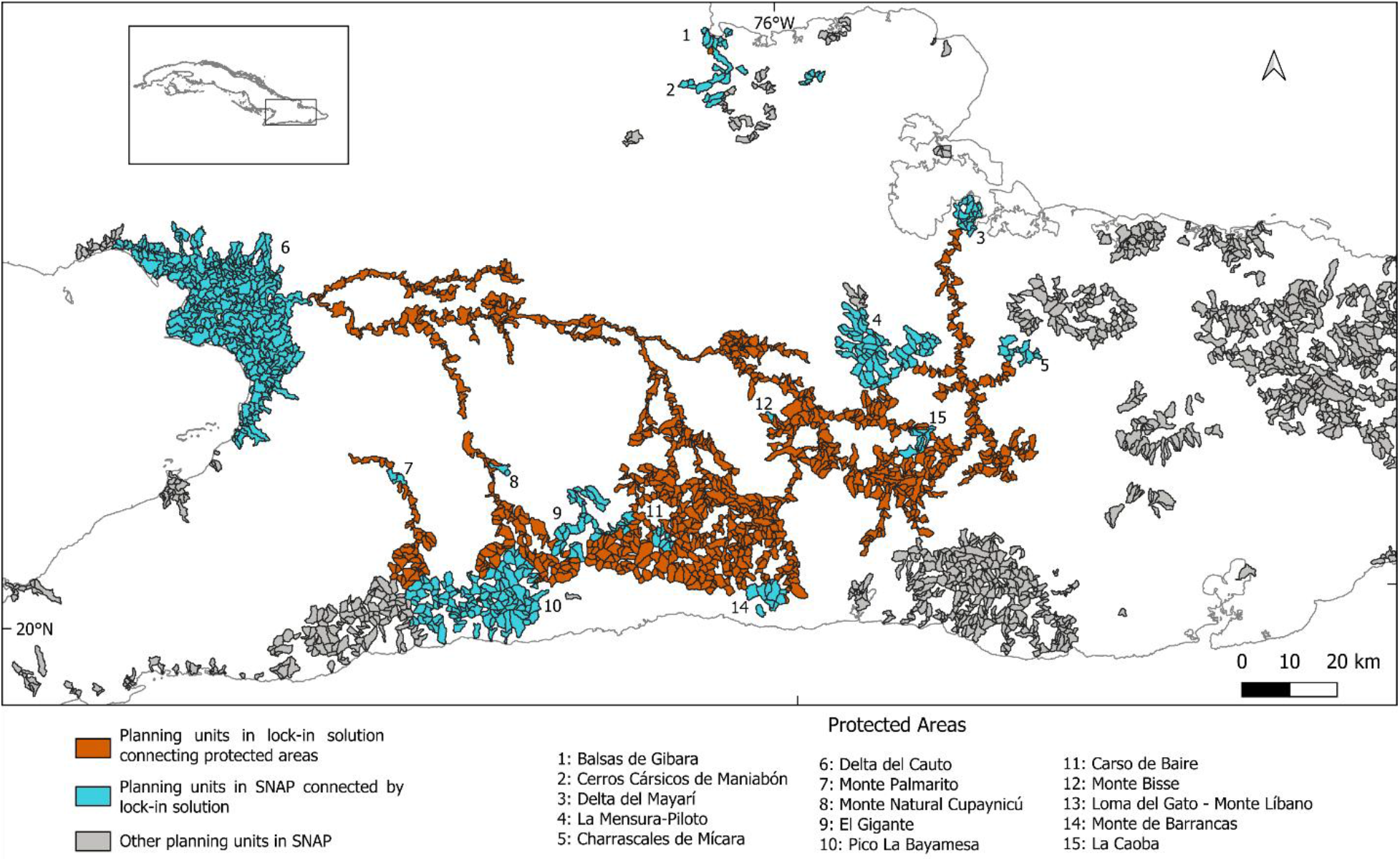
Planning units contributing to protected-area connectivity under the lock-in solution. Grey and blue planning units represent the current Cuban National System of Protected Areas (SNAP), with blue indicating SNAP planning units located within protected areas that became newly connected under the lock-in solution. Orange planning units represent areas selected by the lock-in solution outside SNAP that act as connectors between protected-area pairs that were disconnected under SNAP alone.

Broad spatial patterns of overall planning-unit importance were generally similar between prioritization scenarios, although differences in the location and extent of high-importance basins reflected the influence of retaining existing SNAP planning units in the lock-in solution (Figure 5). Basin-level summaries of irreplaceability revealed spatial variation in the concentration of conservation importance across Cuba’s freshwater planning units. Most basins in the upper 5% of accumulated irreplaceability were located in eastern Cuba.

**Figure 5.**
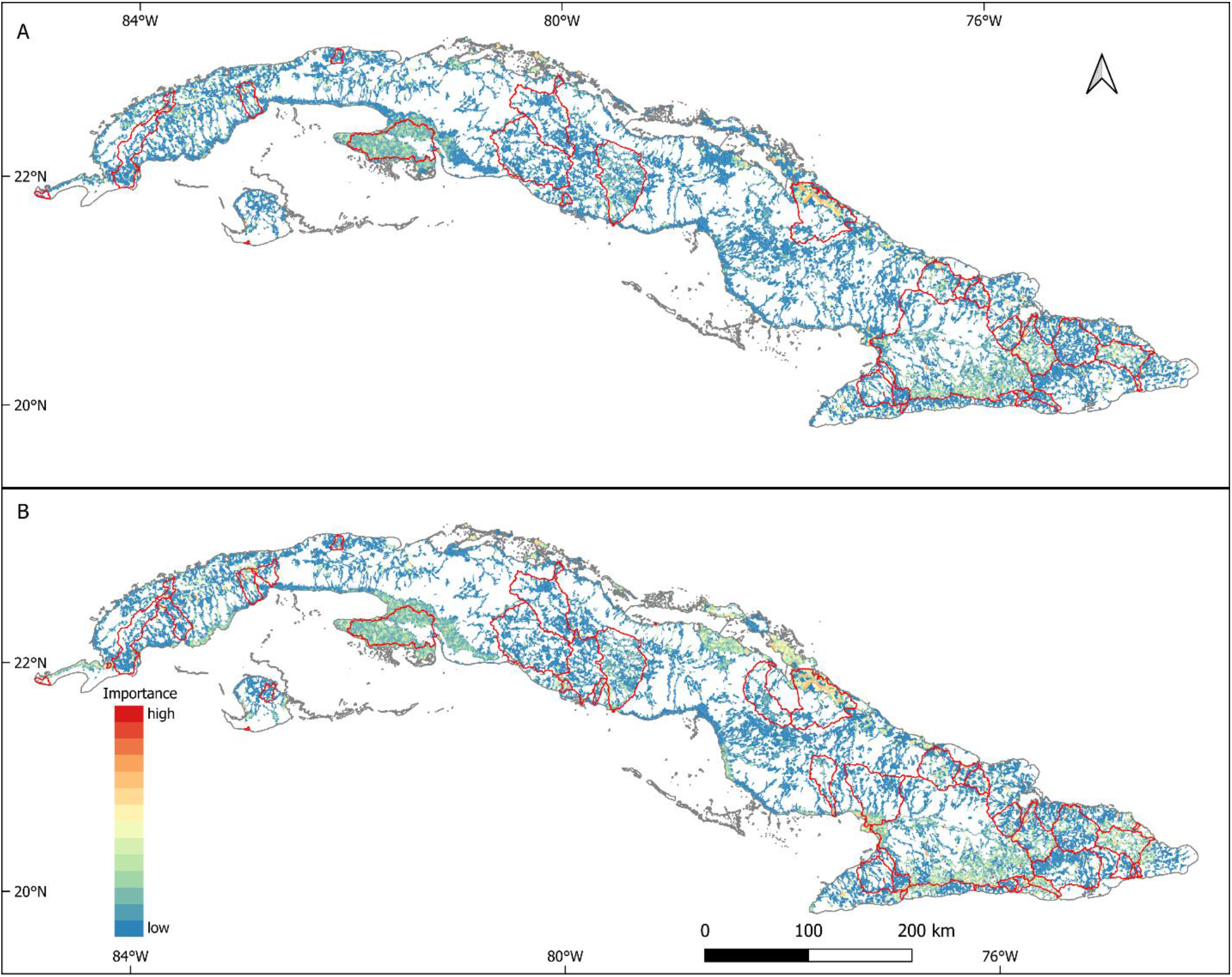
Planning unit importance for meeting the conservation target of 30% coverage for 359 freshwater macroinvertebrate, vertebrate, and macrophyte species across Cuba. Panel A shows the free-choice conservation prioritization scenario, and panel B shows the lock-in scenario. Basins highlighted in red correspond to the top 5% of basins ranked by accumulated irreplaceability. These basins therefore represent the areas with the highest cumulative contribution to achieving freshwater biodiversity representation targets.

Pairwise Spearman correlations calculated across planning units value revealed heterogeneous patterns of spatial congruence among taxonomic groups. Several group pairs showed strong positive congruence in both prioritization scenarios, particularly Coleoptera–Hemiptera, Mollusca–Odonata, and Ephemeroptera–Trichoptera (Figure 6, Table S1.5 Appendix S1 in Supporting Information). However, congruence was not universal. Several pairwise relationships were weak, near zero, or negative, especially those involving Diptera, macrophytes, Mollusca, and Trichoptera (Figure 6, Table S1.5). The strongest negative association was observed between Diptera and macrophytes in both scenarios, suggesting spatial mismatch between areas important for these groups (Table S1.5). Some negative associations became stronger in the lock-in scenario, particularly for Diptera–macrophytes and Diptera–Mollusca (Table S1.5). High-irreplaceability planning units overlap among taxonomic groups was generally limited, indicating that broad spatial congruence in continuous irreplaceability values did not always translate into direct overlap among the highest-priority planning units. Jaccard similarity revealed that only a few taxonomic pairs shared moderate to high proportions of high-irreplaceability planning units (Figure 6, Table S1.6 Appendix S1 in Supporting Information). In both prioritization scenarios, the strongest overlaps involved Decapoda, Hemiptera, vertebrates, Coleoptera, and the Ephemeroptera–Trichoptera pair (Figure 6, Table S1.6). For example, Decapoda–Hemiptera and Decapoda–vertebrates showed the highest overlap in the free-choice scenario, while both pair of taxa were also among the strongest overlaps in the lock-in scenario. In contrast, many pairwise comparisons showed very low Jaccard similarity, particularly those involving Diptera, Mollusca, Trichoptera, and macrophytes, suggesting that the highest-priority planning units for these groups were often spatially distinct.

**Figure 6.**
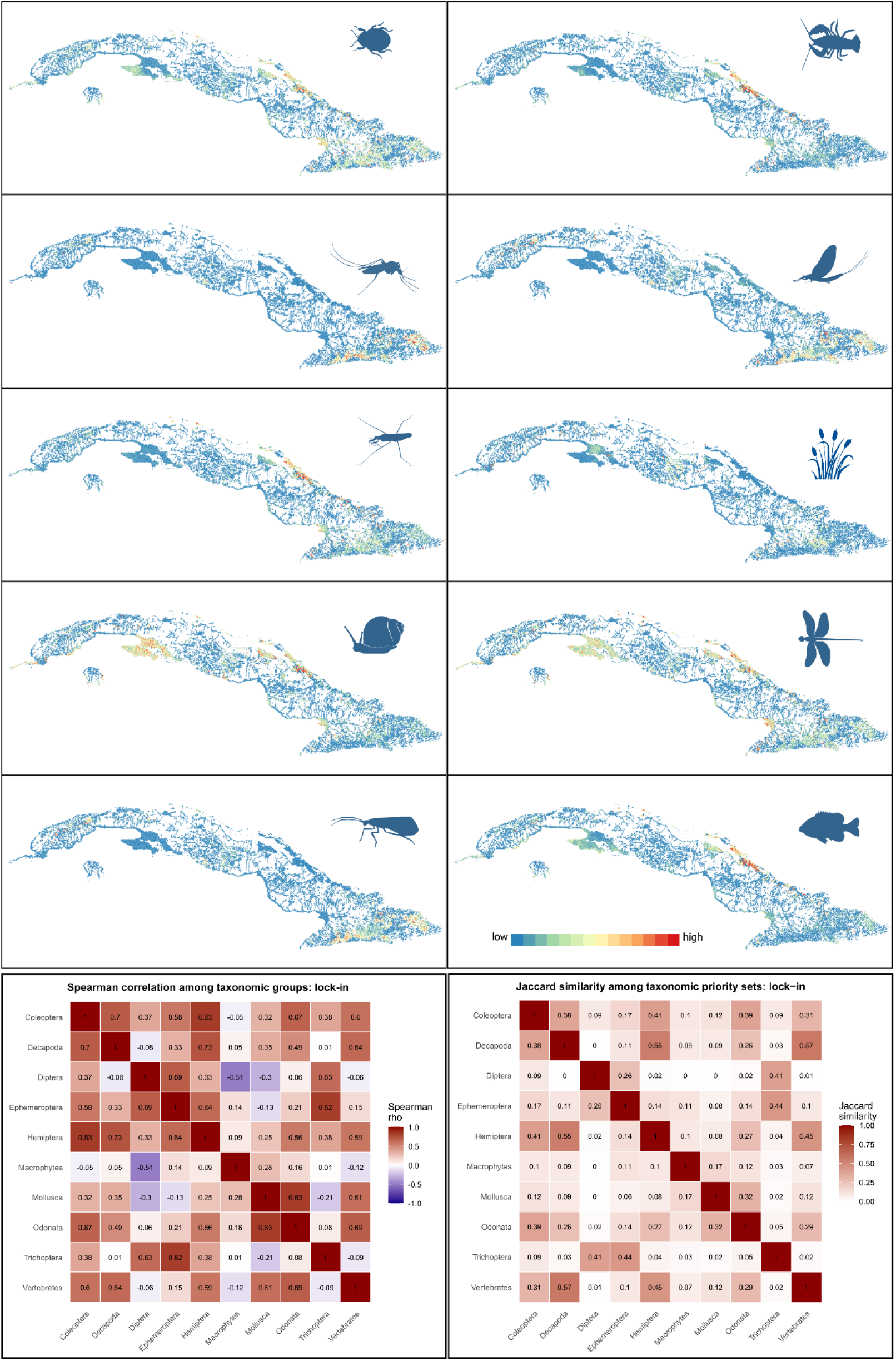
Planning-unit importance for meeting the conservation target of 30% coverage for freshwater macroinvertebrate, vertebrate, and macrophyte species groups across Cuba under the lock-in scenario. The lock-in scenario retained planning units currently included in the Cuban National System of Protected Areas (SNAP) while identifying additional planning units needed to achieve the conservation targets. Irreplaceability values were aggregated by taxonomic group within each planning unit, showing the relative spatial distribution of planning-unit importance for each group. Heatmaps summarize pairwise spatial congruence among taxonomic groups using Spearman correlations of group-specific planning-unit importance values and Jaccard similarity among taxon-specific sets of high-irreplaceability planning units. Spearman correlations indicate similarity in continuous importance patterns, whereas Jaccard similarity indicates the degree of overlap among the highest-importance planning units.

## 4. Discussion

Our analyses show that the current SNAP does not adequately cover freshwater biodiversity, despite Cuba being part of the Caribbean biodiversity hotspot (Mittermeier et al., 2011; Myers et al., 2000). Representation gaps were widespread and strongly uneven across taxonomic groups. Overall, the results reinforce the view that protected-area systems designed primarily from terrestrial perspectives may be poorly aligned with freshwater conservation needs (Tomiczek et al., 2026).

The weak representation of endemic species further underscores the conservation limitations of SNAP. More than 60 % of the assessed Cuban endemic were below the 30% target, and nearly one third were completely absent from the current protected-area system. The lack of protection for endemic species is especially concerning in Cuba, where endemism is a central component of the biodiversity value (Mancina et al., 2017). Endemic species, which typically have restricted geographic distributions, are particularly vulnerable to habitat loss, climate change, hydrological alteration, and local disturbance (Clifford et al., 2025; Fernández-Palacios et al., 2021; Manes et al., 2021). Therefore, an effective system of protected areas must ensure adequately coverage of endemic species’ distribution ranges.

A similar concern applies to species of conservation interest. Although none of the endemic and threatened species examined here was completely absent from SNAP, most still fell below the 30% target. Thus, the current network may provide some degree of incidental protection, but not at levels likely to ensure adequate representation under more ambitious conservation goals. At the same time, the relatively small number of officially recognized threatened freshwater insects should be interpreted cautiously. This likely reflects incomplete Red List assessment, rather than a genuinely low level of threat, since most of the world’s described species have not yet been evaluated and published on the Red List (Bachman et al., 2019; Eisenhauer et al., 2019). Freshwater insect groups such as Ephemeroptera, Trichoptera, and Coleoptera remain poorly assessed in Cuba, and many species with restricted distributions, particularly those confined to mountain ranges or isolated basins, are likely to face elevated extinction risk (Megna et al., 2021; Tagliacollo et al., 2021). In this sense, the present analysis may underestimate the true conservation gap affecting Cuba’s freshwater biodiversity.

Spatial prioritization showed that the 30% representation target is achievable for all freshwater species, but not under the current SNAP configuration. Both the lock-in and free-choice solutions substantially outperformed the existing protected-area system, demonstrating that the present network is insufficient for freshwater conservation but also that this shortfall is tractable. The contrast between scenarios revealed a clear trade-off between institutional continuity and spatial efficiency: retaining SNAP produced a larger, costlier network with higher median species coverage, whereas the free-choice solution achieved the same target with fewer planning units, less area, and lower cost. The limited overlap between SNAP and the free-choice solution further indicates that much of the current network is not positioned in the most efficient locations for representing freshwater biodiversity (Tomiczek et al., 2026).

The value of freshwater-focused prioritization extended beyond species representation alone. In both scenarios, the selected planning units generated substantially larger connected stream-network clusters than SNAP, despite being more spatially dispersed overall. This finding highlights that conservation gains in freshwater systems depend not only on the extent of protected areas but also on how protection is configured along hydrological networks (Hermoso et al., 2016). This was especially evident in the lock-in solution, where strategically selected connector units linked previously isolated protected areas in major basins such as Cauto and Mayarí, showing that large improvements in longitudinal connectivity could be achieved by complementing, rather than replacing, the current network. Taken together, these results suggest that effective freshwater conservation in Cuba will require moving beyond terrestrial reserve logic toward hydrologically structured planning (Hermoso et al., 2015), while recognizing that the most feasible path forward may be to strengthen SNAP through targeted additions in river corridors and connectivity-critical sub-catchments.

The basin-level irreplaceability analyses reveal a strong concentration of freshwater conservation priority areas in eastern Cuba. This regional pattern is consistent with previous recognition of eastern Cuba as a hotspot of freshwater biodiversity and endemism within Cuba (Naranjo-López et al., 2019; Naranjo et al., 2010; Trapero-Quintana & Naranjo-López, 2003). The prominence of eastern basins likely reflects a concentration of species with restricted distributions. This result has direct implications for spatial planning. Although priority basins also occur in central and western Cuba, the concentration of high-irreplaceability basins in the east suggests that national freshwater conservation policy should give particular emphasis to eastern river systems. At the same time, the presence of important basins in other parts of the country indicates that a single-region strategy would be insufficient. The challenge is therefore to combine strong regional focus in the east with a broader network that captures complementary freshwater priorities across the rest of the Cuban archipelago.

The conservation significance of the basins identified here is heightened by the fact that many of Cuba’s river systems have already been extensively modified by reservoirs and associated hydromorphological alterations. Since the 1960s, reservoirs have been widely constructed for flood control, water supply, and agriculture, affecting most major rivers in the country, yet their consequences for freshwater biodiversity remain strikingly understudied (Veloso & Pérez, 2013). Existing Cuban studies have focused mainly on downstream effects on marine fisheries and crustaceans, whereas the implications for freshwater taxa in priority basins have received far less attention (Alvarez-Lajonchère et al., 2018; Baisre & Arboleya, 2006). This gap is especially concerning because dams and flow regulation can disrupt natural flow regimes, sediment and nutrient transport, oxygen dynamics, and longitudinal habitat connectivity, with cascading effects on freshwater communities (Bredenhand & Samways, 2009; Li et al., 2020; Rosero-López et al., 2020; J. Wang et al., 2020). Such alterations may drive upstream extirpation of migratory fish and shrimp by blocking movement, reduce the abundance and richness of benthic invertebrates downstream, and alter food-web structure above barriers through the loss of native consumers (Greathouse et al., 2006; Holmquist et al., 1998). Despite the global evidence for these ecological impacts, reservoir development in Cuba has largely been framed in terms of its social and economic benefits, while its biodiversity costs have remained comparatively overlooked (Veloso & Pérez, 2013). In this context, the high conservation value of many prioritized basins should be interpreted not only as a matter of representation, but also as an urgent call to address the hydrological and ecological degradation already affecting the most critical freshwater systems for biodiversity conservation.

The pronounced taxonomic imbalance in SNAP coverage, combined with the weak and heterogeneous spatial congruence among taxonomic groups, strongly argues against the using surrogate-based approaches for freshwater conservation planning in Cuba. Our results show that assessments of protected area effectiveness are highly contingent on the taxa considered: a network that appears moderately effective for vertebrates or relatively well-studied groups such as odonates can still fail to capture large portions of freshwater biodiversity, particularly invertebrates. This limitation became even more evident in the prioritization analyses, where both pairwise correlations and Jaccard overlap revealed that areas of high conservation importance were often not shared among groups. Even when continuous irreplaceability gradients were broadly similar, overlap among the highest-priority planning units was frequently low, indicating that apparent spatial congruence did not translate into common conservation hotspots. Taken together, these findings show that no single taxonomic group provides a reliable surrogate for freshwater biodiversity as a whole in contraposition with other cases and areas (Slimani et al., 2019), and that effective conservation planning in Cuba must move beyond one-size-fits-all strategies toward explicitly multi-taxon frameworks that reflect the distinct and often non-overlapping spatial priorities of freshwater biota (Stewart et al., 2018; Wang et al., 2018).

At the same time, some limitations of the present study should be acknowledged. Species distribution models remain uncertain for species with limited occurrence records (Araújo et al., 2019), and several freshwater taxa could not be modelled because of insufficient data. In addition, the conservation status of many freshwater insects remains poorly resolved, suggesting that current estimates of underrepresentation for threatened species are likely conservative. The prioritization analyses focused on representation and cost, but did not explicitly account for all socio-political constraints affecting implementation; accordingly, our findings should be viewed as decision-support outputs rather than prescriptive solutions (Araújo, 2025). Even so, these limitations do not alter the main conclusion; if anything, they suggest that the present estimates may underestimate the true extent of freshwater conservation gaps in Cuba and that future analyses should more fully integrate socio-political dimensions.

## 5. Policy recommendations and concluding remarks

Our results show that freshwater biodiversity is not adequately addressed by the current Cuban protected-area system, but they also identify clear pathways for improvement. A first priority should be the strategic expansion of SNAP to close the largest representation gaps, particularly for underrepresented freshwater invertebrates, endemic species, and highly irreplaceable planning units and river corridors specially in eastern Cuba basins. In practice, the lock-in scenario offers a realistic pathway, as it shows that substantial gains can be achieved by building around the existing network rather than replacing it entirely.

At the same time, freshwater biodiversity needs to be incorporated explicitly into conservation policy and management. Protected-area planning in Cuba should move beyond predominantly terrestrial approaches (Rodríguez-Machado & Ponce de León, 2017) and adopt basin-scale, connectivity-based strategies that reflect the ecological structure of freshwater systems (e.g. Hermoso et al., 2016). This includes prioritizing sub-catchments and connector units, maintaining longitudinal connectivity, and protecting headwaters, tributaries, and downstream reaches. Such hydrologically connected conservation units are likely to provide greater ecological resilience under future climate and land-use changes. Because many of the priority basins identified here are already affected by reservoirs (Veloso & Pérez, 2013), conservation policy must also be better integrated with water management, including barrier assessment, environmental flows, and restoration where feasible (Tickner et al., 2020).

Effective implementation will also require stronger research, monitoring, and institutional coordination. Understudied freshwater taxa, especially insects, should be prioritized for additional sampling and Red List assessment, and long-term biodiversity monitoring should be established in priority basins. Freshwater conservation should also be given greater visibility in national biodiversity policy and reporting, in line with international commitments such as the Kunming–Montreal Global Biodiversity Framework (Hughes, 2023). Translating prioritization into effective conservation action will require collaboration among protected-area authorities, water managers, scientists, and local communities, together with adequate resource allocation (e.g. Lindenmayer & Elgar, 2024; Zabala et al., 2024). Together, these measures would help shift conservation in Cuba from a system in which freshwater biodiversity is only incidentally protected to one in which it is explicitly recognized as a central conservation priority.

Our study demonstrates that achieving global freshwater biodiversity targets requires more than expanding protected-area coverage; it requires redesigning conservation networks around freshwater ecological processes. In Cuba, and potentially across other island biodiversity hotspots, effective freshwater conservation will depend on integrating species distributions, hydrological connectivity, and multi-taxon priorities into protected-area planning.

## 6. Data availability statement

The data that support the findings of this study are openly available in Dryad at https://doi.org/10.5061/dryad.hqbzkh1zn.

## Acknowledgment

YT-C and YLD were founded by a Georg Foster Postdoctoral Fellowship of the Alexander von Humboldt Foundation (Ref 3.2 - CUB - 1212347 - GF-P; Ref 3.2 - CUB - 1226121 - GF-P, respectively). YSM was supported by the Alexander von Humboldt Foundation through a Digital Cooperation Fellowship (Ref. 3.4 - CUB / 1161268) and the Universidad Nacional Mayor de San Marcos—RR N° 03202-R-18, project number B18100681 and CONCYTEC through the PROCIENCIA program within the framework of the call “Interinstitutional Alliances for Doctorate Programs”, within the framework of the E033-2023-01-BM in according to contract PE501084299-2023- PROCIENCIA-BM. SD was funded by the Leibniz Competition (J45/2018). Currently, YLD is supported by the Smithsonian Institution, USA, through the Smithsonian Marine Station Postdoctoral Fellowship Program. The authors acknowledge funding by the German Federal Ministry of Education and Research (BMBF grant agreement number no. 033W034A).

