## Appendix S1 in Supporting Information for "Freshwater biodiversity is not adequately addressed by the current protected areas of the Caribbean biodiversity hotspot"

Table S1.1: Table listing the 359 freshwater macroinvertebrates, vertebrates, and macrophyte species analyzed to conduct spatial conservation prioritization across Cuba. N: number of presences.

| | Species | Order | Phylum | N | Cuba endemic |
|---|---|---|---|---|---|
| 1 | *Agonostomus monticola* | Mugiliformes | Chordata | 24 | no |
| 2 | *Alepidomus evermanni* | Atheriniformes | Chordata | 12 | yes |
| 3 | *Alisotrichia* alayoana | Trichoptera | Arthropoda | 4 | yes |
| 4 | *Alisotrichia chiquitica* | Trichoptera | Arthropoda | 4 | yes |
| 5 | *Alisotrichia flintiana* | Trichoptera | Arthropoda | 1 | yes |
| 6 | *Americabaetis naranjoi* | Ephemeroptera | Arthropoda | 6 | yes |
| 7 | *Anax amazili* | Odonata | Arthropoda | 3 | no |
| 8 | *Anax junius* | Odonata | Arthropoda | 7 | no |
| 9 | *Anguilla rostrata* | Anguilliformes | Chordata | 17 | no |
| 10 | *Anodocheilus exiguus* | Coleoptera | Arthropoda | 8 | no |
| 11 | *Antillopsyche wrighti* | Trichoptera | Arthropoda | 2 | yes |
| 12 | *Aphylla caraiba* | Odonata | Arthropoda | 4 | no |
| 13 | *Atopsyche cubana* | Trichoptera | Arthropoda | 2 | yes |
| 14 | *Atopsyche vinai* | Trichoptera | Arthropoda | 7 | yes |
| 15 | *Atya innocous* | Decapoda | Arthropoda | 2 | no |
| 16 | *Atya lanipes* | Decapoda | Arthropoda | 4 | no |
| 17 | *Austrotinodes cubanus* | Trichoptera | Arthropoda | 7 | yes |
| 18 | *Awaous banana* | Perciformes | Chordata | 12 | no |
| 19 | *Azolla caroliniana* | Salviniales | Tracheophyta | 2 | no |
| 20 | *Belostoma minor* | Hemiptera | Arthropoda | 5 | no |
| 21 | *Berosus chevrolati* | Coleoptera | Arthropoda | 5 | yes |
| 22 | *Berosus exiguus* | Coleoptera | Arthropoda | 1 | no |
| 23 | *Berosus infuscatus* | Coleoptera | Arthropoda | 3 | no |
| 24 | *Berosus interstitialis* | Coleoptera | Arthropoda | 10 | no |
| 25 | *Berosus quadridens* | Coleoptera | Arthropoda | 2 | no |
| 26 | *Berosus trilobus* | Coleoptera | Arthropoda | 31 | no |
| 27 | *Berosus undatus* | Coleoptera | Arthropoda | 5 | no |
| 28 | *Bidessonotus browneanus* | Coleoptera | Arthropoda | 24 | no |

| | Species | Order | Phylum | N | Cuba endemic |
|---|---|---|---|---|---|
| 29 | *Biomphalaria havanensis* | Basommatophora | Mollusca | 1 | no |
| 30 | *Biomphalaria helophila* | Basommatophora | Mollusca | 8 | no |
| 31 | *Biomphalaria pallida* | Basommatophora | Mollusca | 1 | no |
| 32 | *Borinquena sexta* | Ephemeroptera | Arthropoda | 1 | yes |
| 33 | *Brachymesia furcata* | Odonata | Arthropoda | 3 | no |
| 34 | *Brachymesia herbida* | Odonata | Arthropoda | 15 | no |
| 35 | *Buenoa antigone* | Hemiptera | Arthropoda | 1 | no |
| 36 | *Buenoa gracilis* | Hemiptera | Arthropoda | 1 | no |
| 37 | *Buenoa platycnemis* | Hemiptera | Arthropoda | 1 | no |
| 38 | *Caenis cubensis* | Ephemeroptera | Arthropoda | 53 | yes |
| 39 | *Callibaetis floridanus* | Ephemeroptera | Arthropoda | 23 | no |
| 40 | *Campsiophora mulata* | Trichoptera | Arthropoda | 3 | yes |
| 41 | *Cannaphila insularis* | Odonata | Arthropoda | 11 | no |
| 42 | *Careospina baconaoi* | Ephemeroptera | Arthropoda | 20 | yes |
| 43 | *Careospina hespera* | Ephemeroptera | Arthropoda | 49 | yes |
| 44 | *Careospina sierramaestrae* | Ephemeroptera | Arthropoda | 1 | yes |
| 45 | *Caribaetis alcarrazae* | Ephemeroptera | Arthropoda | 2 | yes |
| 46 | *Caribaetis planifrons* | Ephemeroptera | Arthropoda | 74 | no |
| 47 | *Cariboptila poquita* | Trichoptera | Arthropoda | 2 | yes |
| 48 | *Cariboptila soltera* | Trichoptera | Arthropoda | 1 | yes |
| 49 | *Celina imitatrix* | Coleoptera | Arthropoda | 3 | no |
| 50 | *Celina palustris* | Coleoptera | Arthropoda | 1 | no |
| 51 | *Celina slossoni* | Coleoptera | Arthropoda | 8 | no |
| 52 | *Cercyon insularis* | Coleoptera | Arthropoda | 7 | no |
| 53 | *Chimarra guapa* | Trichoptera | Arthropoda | 10 | yes |
| 54 | *Chimarra moesta* | Trichoptera | Arthropoda | 2 | yes |
| 55 | *Chimarra pulchra* | Trichoptera | Arthropoda | 14 | yes |
| 56 | *Chimarra quina* | Trichoptera | Arthropoda | 1 | yes |
| 57 | *Cladopelma forcipis* | Diptera | Arthropoda | 1 | no |
| 58 | *Cloeodes inferior* | Ephemeroptera | Arthropoda | 18 | no |
| 59 | *Cloeodes superior* | Ephemeroptera | Arthropoda | 17 | no |

| | Species | Order | Phylum | N | Cuba endemic |
|---|---|---|---|---|---|
| 60 | *Coelotanypus cletis* | Diptera | Arthropoda | 1 | no |
| 61 | *Coelotanypus scapularis* | Diptera | Arthropoda | 1 | no |
| 62 | *Copelatus caelatipennis* | Coleoptera | Arthropoda | 8 | no |
| 63 | *Copelatus cordovai* | Coleoptera | Arthropoda | 3 | yes |
| 64 | *Copelatus cubaensis* | Coleoptera | Arthropoda | 2 | no |
| 65 | *Copelatus insolitus* | Coleoptera | Arthropoda | 9 | no |
| 66 | *Copelatus posticatus* | Coleoptera | Arthropoda | 25 | no |
| 67 | *Corbicula fluminea* | Venerida | Mollusca | 8 | no |
| 68 | *Corisella edulis* | Hemiptera | Arthropoda | 1 | no |
| 69 | *Coryphaeschna adnexa* | Odonata | Arthropoda | 9 | no |
| 70 | *Coryphaeschna viriditas* | Odonata | Arthropoda | 3 | no |
| 71 | *Crocothemis servilia* | Odonata | Arthropoda | 39 | no |
| 72 | *Cubanichthys cubensis* | Cyprinodontiformes | Chordata | 11 | yes |
| 73 | *Cubanoptila botosaneanui* | Trichoptera | Arthropoda | 1 | yes |
| 74 | *Cubanoptila cubana* | Trichoptera | Arthropoda | 1 | yes |
| 75 | *Cubanoptila madremia* | Trichoptera | Arthropoda | 1 | yes |
| 76 | *Cubanoptila muybonita* | Trichoptera | Arthropoda | 3 | yes |
| 77 | *Cubanoptila purpurea* | Trichoptera | Arthropoda | 19 | yes |
| 78 | *Cyprinodon variegatus* | Cyprinodontiformes | Chordata | 9 | no |
| 79 | *Cyrenoida americana* | Venerida | Mollusca | 1 | no |
| 80 | *Dajaus monticola* | Mugiliformes | Chordata | 2 | no |
| 81 | *Desmopachria andreae* | Coleoptera | Arthropoda | 5 | yes |
| 82 | *Desmopachria tarda* | Coleoptera | Arthropoda | 5 | yes |
| 83 | *Dicrotendipes simpsoni* | Diptera | Arthropoda | 1 | no |
| 84 | *Dineutus americanus* | Coleoptera | Arthropoda | 1 | no |
| 85 | *Dineutus longimanus* | Coleoptera | Arthropoda | 24 | yes |
| 86 | *Dormitator maculatus* | Perciformes | Chordata | 8 | no |
| 87 | *Drepanotrema aeruginosum* | Basommatophora | Mollusca | 1 | no |
| 88 | *Drepanotrema anatinum* | Basommatophora | Mollusca | 8 | no |
| 89 | *Drepanotrema cimex* | Basommatophora | Mollusca | 3 | no |
| 90 | *Drepanotrema lucidum* | Basommatophora | Mollusca | 9 | no |

| | Species | Order | Phylum | N | Cuba endemic |
|---|---|---|---|---|---|
| 91 | *Drosera capillaris* | Caryophyllales | Tracheophyta | 9 | no |
| 92 | *Dythemis rufinervis* | Odonata | Arthropoda | 22 | no |
| 93 | *Dythemis sterilis* | Odonata | Arthropoda | 23 | no |
| 94 | *Eleotris pisonis* | Perciformes | Chordata | 12 | no |
| 95 | *Elodea densa* | Alismatales | Tracheophyta | 5 | no |
| 96 | *Elodes angustata* | Coleoptera | Arthropoda | 1 | yes |
| 97 | *Enacantha caribbea* | Odonata | Arthropoda | 1 | no |
| 98 | *Enallagma civile* | Odonata | Arthropoda | 19 | no |
| 99 | *Enallagma coecum* | Odonata | Arthropoda | 49 | no |
| 100 | *Enochrus hamiltoni* | Coleoptera | Arthropoda | 1 | no |
| 101 | *Enochrus pygmaeus* | Coleoptera | Arthropoda | 2 | no |
| 102 | *Epilobocera capolongoi* | Decapoda | Arthropoda | 5 | yes |
| 103 | *Epilobocera cubensis* | Decapoda | Arthropoda | 26 | yes |
| 104 | *Epilobocera diazbeltrani* | Decapoda | Arthropoda | 1 | yes |
| 105 | *Epilobocera gilmanii* | Decapoda | Arthropoda | 2 | yes |
| 106 | *Epilobocera placensis* | Decapoda | Arthropoda | 1 | yes |
| 107 | *Epilobocera synoecia* | Decapoda | Arthropoda | 7 | yes |
| 108 | *Erythemis attala* | Odonata | Arthropoda | 2 | no |
| 109 | *Erythemis plebeja* | Odonata | Arthropoda | 19 | no |
| 110 | *Erythemis simplicicollis* | Odonata | Arthropoda | 10 | no |
| 111 | *Erythemis vesiculosa* | Odonata | Arthropoda | 26 | no |
| 112 | *Erythrodiplax berenice* | Odonata | Arthropoda | 21 | no |
| 113 | *Erythrodiplax bromeliicola* | Odonata | Arthropoda | 4 | no |
| 114 | *Erythrodiplax fervida* | Odonata | Arthropoda | 21 | no |
| 115 | *Erythrodiplax justiniana* | Odonata | Arthropoda | 27 | no |
| 116 | *Erythrodiplax umbrata* | Odonata | Arthropoda | 53 | no |
| 117 | *Eupera cubensis* | Venerida | Mollusca | 5 | yes |
| 118 | *Eurygerris cariniventris* | Hemiptera | Arthropoda | 1 | no |
| 119 | *Eustoma exaltatum* | Gentianales | Tracheophyta | 7 | no |
| 120 | *Fallceon longifolius* | Ephemeroptera | Arthropoda | 7 | no |
| 121 | *Fallceon poeyi* | Ephemeroptera | Arthropoda | 28 | no |

| | Species | Order | Phylum | N | Cuba endemic |
|---|---|---|---|---|---|
| 122 | *Fallceon sextus* | Ephemeroptera | Arthropoda | 1 | yes |
| 123 | *Fallceon testudineus* | Ephemeroptera | Arthropoda | 1 | yes |
| 124 | *Farrodes bimaculatus* | Ephemeroptera | Arthropoda | 42 | yes |
| 125 | *Galba cubensis* | Basommatophora | Mollusca | 17 | no |
| 126 | *Gambusia punctata* | Cyprinodontiformes | Chordata | 49 | yes |
| 127 | *Gambusia puncticulata* | Cyprinodontiformes | Chordata | 49 | no |
| 128 | *Gambusia rhizophorae* | Cyprinodontiformes | Chordata | 2 | no |
| 129 | *Girardinus creolus* | Cyprinodontiformes | Chordata | 4 | yes |
| 130 | *Girardinus denticulatus* | Cyprinodontiformes | Chordata | 16 | yes |
| 131 | *Girardinus falcatus* | Cyprinodontiformes | Chordata | 7 | yes |
| 132 | *Girardinus metallicus* | Cyprinodontiformes | Chordata | 31 | yes |
| 133 | *Girardinus microdactylus* | Cyprinodontiformes | Chordata | 8 | yes |
| 134 | *Girardinus uninonatus* | Cyprinodontiformes | Chordata | 5 | yes |
| 135 | *Girardinus uninotatus* | Cyprinodontiformes | Chordata | 1 | no |
| 136 | *Gobiesox nudus* | Perciformes | Chordata | 1 | no |
| 137 | *Goeldichironomus amazonicus* | Diptera | Arthropoda | 1 | no |
| 138 | *Goeldichironomus devineyae* | Diptera | Arthropoda | 2 | no |
| 139 | *Goeldichironomus holoprasinus* | Diptera | Arthropoda | 1 | no |
| 140 | *Goeldichironomus natans* | Diptera | Arthropoda | 1 | no |
| 141 | *Gundlachia radiata* | Basommatophora | Mollusca | 12 | no |
| 142 | *Gymnochthebius fossatus* | Coleoptera | Arthropoda | 9 | no |
| 143 | *Gynacantha ereagris* | Odonata | Arthropoda | 2 | no |
| 144 | *Gynacantha nervosa* | Odonata | Arthropoda | 9 | no |
| 145 | *Gyraulus parvus* | Basommatophora | Mollusca | 1 | no |
| 146 | *Hagenulus caligatus* | Ephemeroptera | Arthropoda | 9 | yes |
| 147 | *Hagenulus morrisonae* | Ephemeroptera | Arthropoda | 44 | yes |
| 148 | *Haliplus havaniensis* | Coleoptera | Arthropoda | 2 | no |
| 149 | *Haliplus tumidus* | Coleoptera | Arthropoda | 1 | no |
| 150 | *Helicopsyche comosa* | Trichoptera | Arthropoda | 17 | yes |
| 151 | *Helicopsyche cubana* | Trichoptera | Arthropoda | 7 | yes |
| 152 | *Helicopsyche falcigona* | Trichoptera | Arthropoda | 1 | yes |

|  | Species | Order | Phylum | N | Cuba endemic |
|---|---|---|---|---|---|
| 153 | *Helicopsyche hageni* | Trichoptera | Arthropoda | 13 | no |
| 154 | *Helicopsyche occidentalis* | Trichoptera | Arthropoda | 1 | yes |
| 155 | *Hexacylloepus filiformis* | Coleoptera | Arthropoda | 1 | yes |
| 156 | *Hydaticus bimarginatus* | Coleoptera | Arthropoda | 3 | no |
| 157 | *Hydaticus rimosus* | Coleoptera | Arthropoda | 1 | no |
| 158 | *Hydraena decui* | Coleoptera | Arthropoda | 5 | yes |
| 159 | *Hydraena guadelupensis* | Coleoptera | Arthropoda | 4 | no |
| 160 | *Hydraena matthiasi* | Coleoptera | Arthropoda | 1 | yes |
| 161 | *Hydraena perkinsi* | Coleoptera | Arthropoda | 9 | yes |
| 162 | *Hydrocanthus oblongus* | Coleoptera | Arthropoda | 4 | no |
| 163 | *Hydrochus rugosus* | Coleoptera | Arthropoda | 2 | no |
| 164 | *Hydrometra australis* | Hemiptera | Arthropoda | 2 | no |
| 165 | *Hydrometra caraiba* | Hemiptera | Arthropoda | 3 | no |
| 166 | *Hydrometra gibara* | Hemiptera | Arthropoda | 1 | yes |
| 167 | *Hydropsyche cubana* | Trichoptera | Arthropoda | 23 | yes |
| 168 | *Hydropsyche dearmasi* | Trichoptera | Arthropoda | 1 | yes |
| 169 | *Hydrovatus caraibus* | Coleoptera | Arthropoda | 18 | no |
| 170 | *Hydrovatus hornii* | Coleoptera | Arthropoda | 1 | no |
| 171 | *Hymenachne amplexicaulis* | Poales | Tracheophyta | 1 | no |
| 172 | *Hypolestes trinitatis* | Odonata | Arthropoda | 34 | yes |
| 173 | *Idiataphe cubensis* | Odonata | Arthropoda | 2 | no |
| 174 | *Ischnura capreolus* | Odonata | Arthropoda | 13 | no |
| 175 | *Ischnura hastata* | Odonata | Arthropoda | 9 | no |
| 176 | *Ischnura ramburii* | Odonata | Arthropoda | 40 | no |
| 177 | *Jonga serrei* | Decapoda | Arthropoda | 1 | no |
| 178 | *Joturus pichardi* | Mugiliformes | Chordata | 14 | no |
| 179 | *Kryptolebias marmoratus* | Cyprinodontiformes | Chordata | 2 | no |
| 180 | *Labrundinia neopilosella* | Diptera | Arthropoda | 1 | no |
| 181 | *Laccobius antillensis* | Coleoptera | Arthropoda | 1 | no |
| 182 | *Laccomimus pumilio* | Coleoptera | Arthropoda | 3 | no |
| 183 | *Laccophilus bifasciatus* | Coleoptera | Arthropoda | 12 | no |

| | Species | Order | Phylum | N | Cuba endemic |
|---|---|---|---|---|---|
| 184 | *Laccophilus gentilis* | Coleoptera | Arthropoda | 11 | no |
| 185 | *Laccophilus proximus* | Coleoptera | Arthropoda | 30 | no |
| 186 | *Laccophilus vacaensis* | Coleoptera | Arthropoda | 9 | no |
| 187 | *Laccophilus venustus* | Coleoptera | Arthropoda | 22 | no |
| 188 | *Leptobasis vacillans* | Odonata | Arthropoda | 9 | no |
| 189 | *Leptonema poeyi* | Trichoptera | Arthropoda | 1 | no |
| 190 | *Lestes forficula* | Odonata | Arthropoda | 6 | no |
| 191 | *Lestes spumarius* | Odonata | Arthropoda | 1 | no |
| 192 | *Lestes tenuatus* | Odonata | Arthropoda | 6 | no |
| 193 | *Lethocerus colossicus* | Hemiptera | Arthropoda | 1 | no |
| 194 | *Libellula needhami* | Odonata | Arthropoda | 5 | no |
| 195 | *Limia vittata* | Cyprinodontiformes | Chordata | 43 | yes |
| 196 | *Limnogonus franciscanus* | Hemiptera | Arthropoda | 22 | no |
| 197 | *Liodessus noviaffinis* | Coleoptera | Arthropoda | 1 | no |
| 198 | *Littoridinops monroensis* | Littorinomorpha | Mollusca | 1 | no |
| 199 | *Lucifuga dentata* | Ophidiiformes | Chordata | 1 | yes |
| 200 | *Ludwigia erecta* | Myrtales | Tracheophyta | 11 | no |
| 201 | *Ludwigia grandiflora* | Myrtales | Tracheophyta | 2 | no |
| 202 | *Ludwigia leptocarpa* | Myrtales | Tracheophyta | 3 | no |
| 203 | *Ludwigia peduncularis* | Myrtales | Tracheophyta | 2 | no |
| 204 | *Ludwigia peploides* | Myrtales | Tracheophyta | 10 | no |
| 205 | *Ludwigia peruviana* | Myrtales | Tracheophyta | 4 | no |
| 206 | *Ludwigia stricta* | Myrtales | Tracheophyta | 1 | no |
| 207 | *Lutrochus geniculatus* | Coleoptera | Arthropoda | 2 | no |
| 208 | *Macrobrachium acanthurus* | Decapoda | Arthropoda | 12 | no |
| 209 | *Macrobrachium carcinus* | Decapoda | Arthropoda | 2 | no |
| 210 | *Macrobrachium faustinum* | Decapoda | Arthropoda | 11 | no |
| 211 | *Macrobrachium heterochirus* | Decapoda | Arthropoda | 2 | no |
| 212 | *Macrodiplax balteata* | Odonata | Arthropoda | 6 | no |
| 213 | *Macronema tremenda* | Trichoptera | Arthropoda | 2 | yes |
| 214 | *Macrothemis celeno* | Odonata | Arthropoda | 51 | no |

| | Species | Order | Phylum | N | Cuba endemic |
|---|---|---|---|---|---|
| 215 | *Marilia scudderi* | Trichoptera | Arthropoda | 7 | yes |
| 216 | *Marilia wrighti* | Trichoptera | Arthropoda | 1 | yes |
| 217 | *Marisa cornuarietis* | Caenogastropoda | Mollusca | 3 | no |
| 218 | *Megadytes fraternus* | Coleoptera | Arthropoda | 1 | no |
| 219 | *Melanoides tuberculata* | Sorbeoconcha | Mollusca | 9 | no |
| 220 | *Meridiorhantus calidus* | Coleoptera | Arthropoda | 10 | no |
| 221 | *Merragata hebroides* | Hemiptera | Arthropoda | 2 | no |
| 222 | *Mesonoterus addendus* | Coleoptera | Arthropoda | 1 | no |
| 223 | *Mesoplocia inaccessibile* | Ephemeroptera | Arthropoda | 2 | no |
| 224 | *Mesovelia mulsanti* | Hemiptera | Arthropoda | 10 | no |
| 225 | *Metrichia munieca* | Trichoptera | Arthropoda | 1 | yes |
| 226 | *Metrobates tumidus* | Hemiptera | Arthropoda | 6 | no |
| 227 | *Miathyria marcella* | Odonata | Arthropoda | 14 | no |
| 228 | *Miathyria simplex* | Odonata | Arthropoda | 4 | no |
| 229 | *Micrathyria aequalis* | Odonata | Arthropoda | 10 | no |
| 230 | *Micrathyria didyma* | Odonata | Arthropoda | 6 | no |
| 231 | *Micrathyria dissocians* | Odonata | Arthropoda | 6 | no |
| 232 | *Micrathyria hagenii* | Odonata | Arthropoda | 10 | no |
| 233 | *Micratya poeyi* | Decapoda | Arthropoda | 2 | no |
| 234 | *Microvelia albonotata* | Hemiptera | Arthropoda | 1 | no |
| 235 | *Microvelia cubana* | Hemiptera | Arthropoda | 10 | no |
| 236 | *Microvelia paludicola* | Hemiptera | Arthropoda | 1 | no |
| 237 | *Mugil liza* | Mugiliformes | Chordata | 3 | no |
| 238 | *Nandopsis ramsdeni* | Perciformes | Chordata | 11 | yes |
| 239 | *Nandopsis tetracanthus* | Perciformes | Chordata | 37 | yes |
| 240 | *Nectopsyche cubana* | Trichoptera | Arthropoda | 11 | no |
| 241 | *Nehalennia minuta* | Odonata | Arthropoda | 2 | no |
| 242 | *Nelumbo lutea* | Proteales | Tracheophyta | 1 | no |
| 243 | *Neoepilobocera gertraudae* | Decapoda | Arthropoda | 1 | yes |
| 244 | *Neoerythromma cultellatum* | Odonata | Arthropoda | 1 | no |
| 245 | *Neoneura carnatica* | Odonata | Arthropoda | 5 | yes |

| | Species | Order | Phylum | N | Cuba endemic |
|---|---|---|---|---|---|
| 246 | *Neoneura maria* | Odonata | Arthropoda | 50 | yes |
| 247 | *Nereina punctulata* | Neritimorpha | Mollusca | 3 | no |
| 248 | *Nilothauma babiyi* | Diptera | Arthropoda | 1 | no |
| 249 | *Noelmis minima* | Coleoptera | Arthropoda | 1 | yes |
| 250 | *Notomicrus sharpi* | Coleoptera | Arthropoda | 3 | no |
| 251 | *Notonecta indica* | Hemiptera | Arthropoda | 10 | no |
| 252 | *Nymphaea ampla* | Nymphaeales | Tracheophyta | 3 | no |
| 253 | *Nymphaea odorata* | Nymphaeales | Tracheophyta | 7 | no |
| 254 | *Ochrotrichia caramba* | Trichoptera | Arthropoda | 1 | yes |
| 255 | *Ochthebius attritus* | Coleoptera | Arthropoda | 3 | no |
| 256 | *Ophisternon aenigmaticus* | Synbranchiformes | Chordata | 8 | no |
| 257 | *Orthemis discolor* | Odonata | Arthropoda | 7 | no |
| 258 | *Orthemis ferruginea* | Odonata | Arthropoda | 37 | no |
| 259 | *Pachychilus nigratus* | Sorbeoconcha | Mollusca | 2 | yes |
| 260 | *Pachychilus violaceus* | Sorbeoconcha | Mollusca | 5 | yes |
| 261 | *Pachydiplax longipennis* | Odonata | Arthropoda | 5 | no |
| 262 | *Pachydrus obniger* | Coleoptera | Arthropoda | 11 | no |
| 263 | *Palaemon pandaliformis* | Decapoda | Arthropoda | 6 | no |
| 264 | *Paltostoma palominoi* | Diptera | Arthropoda | 1 | no |
| 265 | *Pantala flavescens* | Odonata | Arthropoda | 24 | no |
| 266 | *Pantala hymenaea* | Odonata | Arthropoda | 3 | no |
| 267 | *Paracymus londingi* | Coleoptera | Arthropoda | 1 | no |
| 268 | *Parakiefferiella coronata* | Diptera | Arthropoda | 1 | no |
| 269 | *Paraplea puella* | Hemiptera | Arthropoda | 7 | no |
| 270 | *Pelocoris poeyi* | Hemiptera | Arthropoda | 11 | no |
| 271 | *Pelonomus obscurus* | Coleoptera | Arthropoda | 4 | no |
| 272 | *Perithemis domitia* | Odonata | Arthropoda | 41 | no |
| 273 | *Phaenonotum exstriatum* | Coleoptera | Arthropoda | 1 | no |
| 274 | *Phylloicus chalybeus* | Trichoptera | Arthropoda | 17 | yes |
| 275 | *Phylloicus cubanus* | Trichoptera | Arthropoda | 3 | yes |
| 276 | *Physella acuta* | Basommatophora | Mollusca | 26 | no |

| | Species | Order | Phylum | N | Cuba endemic |
|---|---|---|---|---|---|
| 277 | *Pisidium consanguineum* | Venerida | Mollusca | 1 | yes |
| 278 | *Planorbella duryi* | Basommatophora | Mollusca | 10 | no |
| 279 | *Planorbella trivolvis* | Basommatophora | Mollusca | 1 | no |
| 280 | *Platyvelia brachialis* | Hemiptera | Arthropoda | 1 | no |
| 281 | *Poecilophlebia pacoi* | Ephemeroptera | Arthropoda | 3 | yes |
| 282 | *Polycentropus mathisi* | Trichoptera | Arthropoda | 2 | yes |
| 283 | *Polycentropus turquino* | Trichoptera | Arthropoda | 1 | yes |
| 284 | *Pomacea paludosa* | Caenogastropoda | Mollusca | 18 | no |
| 285 | *Pomacea poeyana* | Caenogastropoda | Mollusca | 8 | yes |
| 286 | *Potimirin americana* | Decapoda | Arthropoda | 2 | no |
| 287 | *Procambarus cubensis* | Decapoda | Arthropoda | 3 | yes |
| 288 | *Procambarus niveus* | Decapoda | Arthropoda | 1 | yes |
| 289 | *Progomphus integer* | Odonata | Arthropoda | 20 | no |
| 290 | *Protoneura caligata* | Odonata | Arthropoda | 8 | yes |
| 291 | *Protoneura capillaris* | Odonata | Arthropoda | 27 | yes |
| 292 | *Protoneura viridis* | Odonata | Arthropoda | 2 | no |
| 293 | *Pseudosuccinea columella* | Basommatophora | Mollusca | 11 | no |
| 294 | *Psilopelmia hematopotum* | Diptera | Arthropoda | 11 | no |
| 295 | *Psilopelmia ochraceum* | Diptera | Arthropoda | 1 | no |
| 296 | *Pyrgophorus parvulus* | Littorinomorpha | Mollusca | 8 | no |
| 297 | *Quintana atrizona* | Cyprinodontiformes | Chordata | 1 | yes |
| 298 | *Ramphocorixa rotundocephala* | Hemiptera | Arthropoda | 1 | no |
| 299 | *Ranatra fabricii* | Hemiptera | Arthropoda | 6 | yes |
| 300 | *Ranatra sagrai* | Hemiptera | Arthropoda | 1 | yes |
| 301 | *Remartinia secreta* | Odonata | Arthropoda | 1 | no |
| 302 | *Rhagovelia collaris* | Hemiptera | Arthropoda | 38 | no |
| 303 | *Rhagovelia mira* | Hemiptera | Arthropoda | 2 | yes |
| 304 | *Rheumatobates clanis* | Hemiptera | Arthropoda | 2 | no |
| 305 | *Rheumatobates meinerti* | Hemiptera | Arthropoda | 2 | no |
| 306 | *Rhionaeschna psilus* | Odonata | Arthropoda | 12 | no |
| 307 | *Rivulus cylindraceus* | Cyprinodontiformes | Chordata | 5 | yes |

| | Species | Order | Phylum | N | Cuba endemic |
|---|---|---|---|---|---|
| 308 | *Sacciolepis striata* | Poales | Tracheophyta | 1 | no |
| 309 | *Sagittaria intermedia* | Alismatales | Tracheophyta | 1 | no |
| 310 | *Sagittaria lancifolia* | Alismatales | Tracheophyta | 7 | no |
| 311 | *Salvinia auriculata* | Salviniales | Tracheophyta | 2 | no |
| 312 | *Scapanea frontalis* | Odonata | Arthropoda | 28 | no |
| 313 | *Setaria geminata* | Poales | Tracheophyta | 1 | no |
| 314 | *Sicydium plumieri* | Perciformes | Chordata | 17 | no |
| 315 | *Smicridea comma* | Trichoptera | Arthropoda | 59 | no |
| 316 | *Smicridea obesa* | Trichoptera | Arthropoda | 1 | yes |
| 317 | *Steinovelia stagnalis* | Hemiptera | Arthropoda | 2 | no |
| 318 | *Suphis inflatus* | Coleoptera | Arthropoda | 1 | no |
| 319 | *Suphisellus insularis* | Coleoptera | Arthropoda | 4 | no |
| 320 | *Suphisellus nigrinus* | Coleoptera | Arthropoda | 3 | no |
| 321 | *Suphisellus tenuicornis* | Coleoptera | Arthropoda | 1 | yes |
| 322 | *Sympetrum illotum* | Odonata | Arthropoda | 3 | no |
| 323 | *Tanytarsus limneticus* | Diptera | Arthropoda | 1 | no |
| 324 | *Tarebia granifera* | Sorbeoconcha | Mollusca | 45 | no |
| 325 | *Tauriphila australis* | Odonata | Arthropoda | 4 | no |
| 326 | *Telebasis dominicana* | Odonata | Arthropoda | 23 | no |
| 327 | *Thermonectus basillaris* | Coleoptera | Arthropoda | 11 | no |
| 328 | *Thermonectus circumscriptus* | Coleoptera | Arthropoda | 10 | no |
| 329 | *Thermonectus margineguttatus* | Coleoptera | Arthropoda | 2 | no |
| 330 | *Thermonectus succinctus* | Coleoptera | Arthropoda | 10 | no |
| 331 | *Tholymis citrina* | Odonata | Arthropoda | 9 | no |
| 332 | *Tramea abdominalis* | Odonata | Arthropoda | 25 | no |
| 333 | *Tramea calverti* | Odonata | Arthropoda | 3 | no |
| 334 | *Tramea insularis* | Odonata | Arthropoda | 3 | no |
| 335 | *Tramea onusta* | Odonata | Arthropoda | 12 | no |
| 336 | *Traverina cubensis* | Ephemeroptera | Arthropoda | 2 | yes |
| 337 | *Traverina oriente* | Ephemeroptera | Arthropoda | 19 | yes |
| 338 | *Trepobates taylori* | Hemiptera | Arthropoda | 6 | no |

| | Species | Order | Phylum | N | Cuba endemic |
|---|---|---|---|---|---|
| 339 | *Triacanthagyna septima* | Odonata | Arthropoda | 5 | no |
| 340 | *Triacanthagyna trifida* | Odonata | Arthropoda | 2 | no |
| 341 | *Tricorythodes cubensis* | Ephemeroptera | Arthropoda | 17 | yes |
| 342 | *Tricorythodes grallator* | Ephemeroptera | Arthropoda | 30 | yes |
| 343 | *Tricorythodes montanus* | Ephemeroptera | Arthropoda | 14 | yes |
| 344 | *Tricorythodes sacculobranchis* | Ephemeroptera | Arthropoda | 44 | yes |
| 345 | *Tricorythodes sierramaestrae* | Ephemeroptera | Arthropoda | 2 | yes |
| 346 | *Tropisternus chalybeus* | Coleoptera | Arthropoda | 1 | no |
| 347 | *Tropisternus collaris* | Coleoptera | Arthropoda | 1 | yes |
| 348 | *Tropisternus lateralis* | Coleoptera | Arthropoda | 4 | yes |
| 349 | *Tropisternus mergus* | Coleoptera | Arthropoda | 1 | yes |
| 350 | *Turquinophlebia grandis* | Ephemeroptera | Arthropoda | 2 | yes |
| 351 | *Vitta usnea* | Neritimorpha | Mollusca | 1 | no |
| 352 | *Vitta virginea* | Neritimorpha | Mollusca | 4 | no |
| 353 | *Xiphocaris elongata* | Decapoda | Arthropoda | 10 | no |
| 354 | *Xiphocaris gomezi* | Decapoda | Arthropoda | 4 | yes |
| 355 | *Xiphocentron cubanum* | Trichoptera | Arthropoda | 6 | yes |
| 356 | *Xyris ambigua* | Poales | Tracheophyta | 3 | no |
| 357 | *Xyris grandiceps* | Poales | Tracheophyta | 1 | no |
| 358 | *Xyris jupicai* | Poales | Tracheophyta | 1 | no |
| 359 | *Xyris navicularis* | Poales | Tracheophyta | 2 | no |

Table S1.2: Mean (m) and standard deviation (sd) of performance metrics for ensemble Random Forest species distribution models fitted for freshwater species of Cuba. sTSS: standardized True Skill Statistic; AUC: Area Under the Receiver Operating Characteristic Curve; Sen: sensitivity; Spec: specificity; Om: omission rate; Com: commission rate; n_p: number of presences used to fit the models; n_f_m: number of models fitted across species-specific cross-validation runs.

| | Species | n_p | AUC_m | AUC_sd | sTSS_m | sTSS_sd | Sen_m | Sen_sd | Spec_m | Spec_sd | Om_m | Om_sd | Com_m | Com_sd | n_f_m |
|---|---|---|---|---|---|---|---|---|---|---|---|---|---|---|---|
| 1 | *Agonostomus monticola* | 24 | 0.9 | 0.091 | 0.85 | 0.122 | 0.85 | 0.127 | 0.84 | 0.12 | 0.15 | 0.127 | 0.16 | 0.12 | 15 |
| 2 | *Alepidomus evermanni* | 12 | 0.94 | 0.04 | 0.87 | 0.072 | 0.87 | 0.07 | 0.88 | 0.081 | 0.13 | 0.07 | 0.12 | 0.081 | 10 |
| 3 | *Anguilla rostrata* | 17 | 0.97 | 0.025 | 0.9 | 0.058 | 0.9 | 0.085 | 0.9 | 0.056 | 0.1 | 0.085 | 0.1 | 0.056 | 15 |
| 4 | *Awaous banana* | 12 | 0.94 | 0.065 | 0.85 | 0.088 | 0.85 | 0.095 | 0.86 | 0.088 | 0.15 | 0.095 | 0.14 | 0.088 | 10 |
| 5 | *Berosus interstitialis* | 10 | 0.76 | 0.089 | 0.68 | 0.132 | 0.68 | 0.14 | 0.67 | 0.134 | 0.32 | 0.14 | 0.33 | 0.134 | 10 |
| 6 | *Berosus trilobus* | 31 | 0.98 | 0.027 | 0.91 | 0.074 | 0.91 | 0.082 | 0.92 | 0.07 | 0.09 | 0.082 | 0.08 | 0.07 | 25 |
| 7 | *Bidessonotus browneanus* | 24 | 0.83 | 0.133 | 0.78 | 0.152 | 0.78 | 0.172 | 0.78 | 0.139 | 0.22 | 0.172 | 0.22 | 0.139 | 15 |
| 8 | *Brachymesia herbida* | 15 | 0.85 | 0.077 | 0.77 | 0.115 | 0.76 | 0.135 | 0.78 | 0.101 | 0.24 | 0.135 | 0.22 | 0.101 | 15 |
| 9 | *Caenis cubensis* | 53 | 0.95 | 0.039 | 0.88 | 0.065 | 0.89 | 0.067 | 0.88 | 0.066 | 0.11 | 0.067 | 0.12 | 0.066 | 25 |
| 10 | *Callibaetis floridanus* | 23 | 0.87 | 0.087 | 0.79 | 0.094 | 0.79 | 0.104 | 0.79 | 0.088 | 0.21 | 0.104 | 0.21 | 0.088 | 15 |
| 11 | *Cannaphila insularis* | 11 | 0.85 | 0.135 | 0.78 | 0.156 | 0.79 | 0.168 | 0.78 | 0.15 | 0.21 | 0.168 | 0.22 | 0.15 | 10 |
| 12 | *Careospina baconaoi* | 20 | 0.9 | 0.086 | 0.82 | 0.096 | 0.82 | 0.089 | 0.83 | 0.108 | 0.18 | 0.089 | 0.17 | 0.108 | 15 |
| 13 | *Careospina hespera* | 49 | 0.92 | 0.063 | 0.85 | 0.085 | 0.85 | 0.089 | 0.85 | 0.084 | 0.15 | 0.089 | 0.15 | 0.084 | 25 |
| 14 | *Caribaetis planifrons* | 74 | 0.97 | 0.026 | 0.91 | 0.057 | 0.91 | 0.06 | 0.9 | 0.056 | 0.09 | 0.06 | 0.1 | 0.056 | 25 |
| 15 | *Chimarra guapa* | 10 | 0.8 | 0.128 | 0.69 | 0.12 | 0.68 | 0.14 | 0.7 | 0.105 | 0.32 | 0.14 | 0.3 | 0.105 | 10 |
| 16 | *Chimarra pulchra* | 14 | 0.75 | 0.106 | 0.69 | 0.118 | 0.71 | 0.117 | 0.68 | 0.123 | 0.29 | 0.117 | 0.32 | 0.123 | 7 |
| 17 | *Cloeodes inferior* | 18 | 0.9 | 0.065 | 0.84 | 0.071 | 0.82 | 0.076 | 0.85 | 0.078 | 0.18 | 0.076 | 0.15 | 0.078 | 15 |
| 18 | *Cloeodes superior* | 17 | 0.97 | 0.028 | 0.91 | 0.076 | 0.91 | 0.091 | 0.92 | 0.07 | 0.09 | 0.091 | 0.08 | 0.07 | 15 |
| 19 | *Copelatus posticatus* | 25 | 0.9 | 0.063 | 0.81 | 0.097 | 0.81 | 0.111 | 0.8 | 0.089 | 0.19 | 0.111 | 0.2 | 0.089 | 15 |
| 20 | *Crocothemis servilia* | 39 | 0.92 | 0.055 | 0.85 | 0.067 | 0.86 | 0.068 | 0.84 | 0.071 | 0.14 | 0.068 | 0.16 | 0.071 | 25 |
| 21 | *Cubanichthys cubensis* | 11 | 0.8 | 0.063 | 0.7 | 0.08 | 0.69 | 0.089 | 0.7 | 0.086 | 0.31 | 0.089 | 0.3 | 0.086 | 10 |
| 22 | *Cubanoptila purpurea* | 19 | 0.95 | 0.036 | 0.87 | 0.064 | 0.88 | 0.073 | 0.86 | 0.069 | 0.12 | 0.073 | 0.14 | 0.069 | 15 |
| 23 | *Dineutus longimanus* | 24 | 0.99 | 0.016 | 0.95 | 0.059 | 0.96 | 0.061 | 0.95 | 0.059 | 0.04 | 0.061 | 0.05 | 0.059 | 15 |
| 24 | *Dythemis rufinervis* | 22 | 0.88 | 0.063 | 0.81 | 0.075 | 0.8 | 0.088 | 0.81 | 0.068 | 0.2 | 0.088 | 0.19 | 0.068 | 15 |
| 25 | *Dythemis sterilis* | 23 | 0.83 | 0.09 | 0.74 | 0.087 | 0.73 | 0.088 | 0.75 | 0.095 | 0.27 | 0.088 | 0.25 | 0.095 | 14 |
| 26 | *Eleotris pisonis* | 12 | 0.89 | 0.063 | 0.82 | 0.033 | 0.83 | 0 | 0.82 | 0.066 | 0.17 | 0 | 0.18 | 0.066 | 10 |

| | Species | n_p | AUC_m | AUC_sd | sTSS_m | sTSS_sd | Sen_m | Sen_sd | Spec_m | Spec_sd | Om_m | Om_sd | Com_m | Com_sd | n_f_m |
|---|---|---|---|---|---|---|---|---|---|---|---|---|---|---|---|
| 27 | *Enallagma civile* | 19 | 0.8 | 0.097 | 0.76 | 0.107 | 0.76 | 0.125 | 0.77 | 0.096 | 0.24 | 0.125 | 0.23 | 0.096 | 15 |
| 28 | *Enallagma coecum* | 49 | 0.95 | 0.036 | 0.88 | 0.062 | 0.88 | 0.064 | 0.87 | 0.065 | 0.12 | 0.064 | 0.13 | 0.065 | 25 |
| 29 | *Epilobocera cubensis* | 26 | 0.9 | 0.059 | 0.85 | 0.062 | 0.85 | 0.071 | 0.85 | 0.055 | 0.15 | 0.071 | 0.15 | 0.055 | 15 |
| 30 | *Erythemis plebeja* | 19 | 0.83 | 0.097 | 0.74 | 0.107 | 0.74 | 0.127 | 0.74 | 0.09 | 0.26 | 0.127 | 0.26 | 0.09 | 15 |
| 31 | *Erythemis simplicicollis* | 10 | 0.78 | 0.123 | 0.73 | 0.125 | 0.73 | 0.141 | 0.72 | 0.12 | 0.27 | 0.141 | 0.28 | 0.12 | 9 |
| 32 | *Erythemis vesiculosa* | 26 | 0.89 | 0.055 | 0.8 | 0.085 | 0.81 | 0.087 | 0.8 | 0.089 | 0.19 | 0.087 | 0.2 | 0.089 | 14 |
| 33 | *Erythrodiplax berenice* | 21 | 0.85 | 0.074 | 0.76 | 0.11 | 0.76 | 0.117 | 0.76 | 0.11 | 0.24 | 0.117 | 0.24 | 0.11 | 15 |
| 34 | *Erythrodiplax fervida* | 21 | 0.82 | 0.07 | 0.74 | 0.092 | 0.76 | 0.104 | 0.73 | 0.085 | 0.24 | 0.104 | 0.27 | 0.085 | 14 |
| 35 | *Erythrodiplax justiniana* | 27 | 0.85 | 0.062 | 0.77 | 0.068 | 0.78 | 0.073 | 0.77 | 0.067 | 0.22 | 0.073 | 0.23 | 0.067 | 15 |
| 36 | *Erythrodiplax umbrata* | 53 | 0.93 | 0.047 | 0.85 | 0.076 | 0.85 | 0.077 | 0.85 | 0.075 | 0.15 | 0.077 | 0.15 | 0.075 | 25 |
| 37 | *Fallceon poeyi* | 28 | 0.9 | 0.079 | 0.82 | 0.095 | 0.82 | 0.1 | 0.81 | 0.094 | 0.18 | 0.1 | 0.19 | 0.094 | 15 |
| 38 | *Farrodes bimaculatus* | 42 | 0.91 | 0.062 | 0.84 | 0.072 | 0.84 | 0.074 | 0.83 | 0.074 | 0.16 | 0.074 | 0.17 | 0.074 | 25 |
| 39 | *Galba cubensis* | 17 | 0.93 | 0.067 | 0.86 | 0.092 | 0.86 | 0.097 | 0.87 | 0.091 | 0.14 | 0.097 | 0.13 | 0.091 | 15 |
| 40 | *Gambusia punctata* | 49 | 0.9 | 0.054 | 0.82 | 0.078 | 0.83 | 0.079 | 0.82 | 0.082 | 0.17 | 0.079 | 0.18 | 0.082 | 25 |
| 41 | *Gambusia puncticulata* | 49 | 0.94 | 0.044 | 0.88 | 0.072 | 0.87 | 0.072 | 0.88 | 0.075 | 0.13 | 0.072 | 0.12 | 0.075 | 25 |
| 42 | *Girardinus denticulatus* | 16 | 0.88 | 0.096 | 0.81 | 0.117 | 0.81 | 0.125 | 0.81 | 0.113 | 0.19 | 0.125 | 0.19 | 0.113 | 15 |
| 43 | *Girardinus metallicus* | 31 | 0.9 | 0.066 | 0.83 | 0.091 | 0.83 | 0.093 | 0.83 | 0.094 | 0.17 | 0.093 | 0.17 | 0.094 | 25 |
| 44 | *Gundlachia radiata* | 12 | 0.78 | 0.086 | 0.71 | 0.105 | 0.72 | 0.118 | 0.7 | 0.103 | 0.28 | 0.118 | 0.3 | 0.103 | 9 |
| 45 | *Hagenulus morrisonae* | 44 | 0.91 | 0.039 | 0.83 | 0.071 | 0.83 | 0.075 | 0.83 | 0.073 | 0.17 | 0.075 | 0.17 | 0.073 | 25 |
| 46 | *Helicopsyche comosa* | 17 | 0.94 | 0.034 | 0.87 | 0.064 | 0.86 | 0.074 | 0.88 | 0.064 | 0.14 | 0.074 | 0.12 | 0.064 | 15 |
| 47 | *Helicopsyche hageni* | 13 | 0.86 | 0.068 | 0.8 | 0.068 | 0.8 | 0.072 | 0.81 | 0.065 | 0.2 | 0.072 | 0.19 | 0.065 | 10 |
| 48 | *Hydropsyche cubana* | 23 | 0.96 | 0.03 | 0.91 | 0.069 | 0.91 | 0.08 | 0.9 | 0.061 | 0.09 | 0.08 | 0.1 | 0.061 | 15 |
| 49 | *Hydrovatus caraibus* | 18 | 0.85 | 0.1 | 0.79 | 0.108 | 0.81 | 0.11 | 0.77 | 0.11 | 0.19 | 0.11 | 0.23 | 0.11 | 14 |
| 50 | *Hypolestes trinitatis* | 34 | 0.93 | 0.08 | 0.89 | 0.087 | 0.91 | 0.093 | 0.88 | 0.086 | 0.09 | 0.093 | 0.12 | 0.086 | 25 |
| 51 | *Ischnura capreolus* | 13 | 0.75 | 0.103 | 0.68 | 0.102 | 0.7 | 0.099 | 0.66 | 0.11 | 0.3 | 0.099 | 0.34 | 0.11 | 9 |
| 52 | *Ischnura ramburii* | 40 | 0.93 | 0.055 | 0.87 | 0.07 | 0.86 | 0.071 | 0.87 | 0.073 | 0.14 | 0.071 | 0.13 | 0.073 | 25 |
| 53 | *Joturus pichardi* | 14 | 0.8 | 0.079 | 0.73 | 0.083 | 0.74 | 0.09 | 0.72 | 0.079 | 0.26 | 0.09 | 0.28 | 0.079 | 10 |
| 54 | *Laccophilus bifasciatus* | 12 | 0.96 | 0.038 | 0.91 | 0.081 | 0.9 | 0.086 | 0.92 | 0.079 | 0.1 | 0.086 | 0.08 | 0.079 | 10 |
| 55 | *Laccophilus gentilis* | 11 | 0.82 | 0.095 | 0.77 | 0.115 | 0.76 | 0.127 | 0.77 | 0.107 | 0.24 | 0.127 | 0.23 | 0.107 | 10 |
| 56 | *Laccophilus proximus* | 30 | 0.88 | 0.078 | 0.84 | 0.099 | 0.84 | 0.113 | 0.84 | 0.088 | 0.16 | 0.113 | 0.16 | 0.088 | 25 |
| 57 | *Laccophilus venustus* | 22 | 0.95 | 0.042 | 0.89 | 0.08 | 0.89 | 0.079 | 0.89 | 0.082 | 0.11 | 0.079 | 0.11 | 0.082 | 15 |

| | Species | n_p | AUC_m | AUC_sd | sTSS_m | sTSS_sd | Sen_m | Sen_sd | Spec_m | Spec_sd | Om_m | Om_sd | Com_m | Com_sd | n_f_m |
|---|---|---|---|---|---|---|---|---|---|---|---|---|---|---|---|
| 58 | *Limia vittata* | 43 | 0.93 | 0.053 | 0.87 | 0.063 | 0.87 | 0.068 | 0.87 | 0.062 | 0.13 | 0.068 | 0.13 | 0.062 | 25 |
| 59 | *Limnogonus franciscanus* | 22 | 0.95 | 0.034 | 0.87 | 0.065 | 0.87 | 0.085 | 0.88 | 0.049 | 0.13 | 0.085 | 0.12 | 0.049 | 15 |
| 60 | *Ludwigia erecta* | 11 | 0.76 | 0.079 | 0.66 | 0.078 | 0.66 | 0.066 | 0.66 | 0.094 | 0.34 | 0.066 | 0.34 | 0.094 | 8 |
| 61 | *Ludwigia peploides* | 10 | 0.79 | 0.063 | 0.7 | 0.1 | 0.68 | 0.103 | 0.72 | 0.103 | 0.32 | 0.103 | 0.28 | 0.103 | 10 |
| 62 | *Macrobrachium acanthurus* | 12 | 0.88 | 0.092 | 0.81 | 0.086 | 0.81 | 0.1 | 0.81 | 0.083 | 0.19 | 0.1 | 0.19 | 0.083 | 9 |
| 63 | *Macrobrachium faustinum* | 11 | 0.89 | 0.06 | 0.82 | 0.078 | 0.82 | 0.081 | 0.82 | 0.086 | 0.18 | 0.081 | 0.18 | 0.086 | 10 |
| 64 | *Macrothemis celeno* | 51 | 0.88 | 0.069 | 0.8 | 0.084 | 0.81 | 0.091 | 0.8 | 0.079 | 0.19 | 0.091 | 0.2 | 0.079 | 25 |
| 65 | *Meridiorhantus calidus* | 10 | 0.97 | 0.041 | 0.94 | 0.094 | 0.94 | 0.097 | 0.93 | 0.095 | 0.06 | 0.097 | 0.07 | 0.095 | 10 |
| 66 | *Mesovelia mulsanti* | 10 | 0.95 | 0.088 | 0.89 | 0.139 | 0.9 | 0.17 | 0.88 | 0.114 | 0.1 | 0.17 | 0.12 | 0.114 | 10 |
| 67 | *Miathyria marcella* | 14 | 0.76 | 0.093 | 0.69 | 0.079 | 0.7 | 0.081 | 0.69 | 0.09 | 0.3 | 0.081 | 0.31 | 0.09 | 10 |
| 68 | *Micrathyria aequalis* | 10 | 0.76 | 0.141 | 0.71 | 0.15 | 0.71 | 0.176 | 0.71 | 0.127 | 0.29 | 0.176 | 0.29 | 0.127 | 9 |
| 69 | *Micrathyria hagenii* | 10 | 0.7 | 0.125 | 0.69 | 0.131 | 0.7 | 0.115 | 0.68 | 0.15 | 0.3 | 0.115 | 0.32 | 0.15 | 4 |
| 70 | *Microvelia cubana* | 10 | 0.93 | 0.069 | 0.88 | 0.1 | 0.88 | 0.103 | 0.89 | 0.099 | 0.12 | 0.103 | 0.11 | 0.099 | 10 |
| 71 | *Nandopsis ramsdeni* | 11 | 0.88 | 0.068 | 0.79 | 0.12 | 0.8 | 0.133 | 0.78 | 0.115 | 0.2 | 0.133 | 0.22 | 0.115 | 10 |
| 72 | *Nandopsis tetracanthus* | 37 | 0.94 | 0.049 | 0.88 | 0.074 | 0.88 | 0.087 | 0.87 | 0.063 | 0.12 | 0.087 | 0.13 | 0.063 | 25 |
| 73 | *Nectopsyche cubana* | 11 | 0.8 | 0.115 | 0.73 | 0.115 | 0.74 | 0.134 | 0.73 | 0.105 | 0.26 | 0.134 | 0.27 | 0.105 | 10 |
| 74 | *Neoneura maria* | 50 | 0.91 | 0.056 | 0.82 | 0.072 | 0.82 | 0.076 | 0.83 | 0.074 | 0.18 | 0.076 | 0.17 | 0.074 | 25 |
| 75 | *Notonecta indica* | 10 | 0.76 | 0.15 | 0.71 | 0.137 | 0.7 | 0.141 | 0.72 | 0.14 | 0.3 | 0.141 | 0.28 | 0.14 | 10 |
| 76 | *Orthemis ferruginea* | 37 | 0.92 | 0.062 | 0.84 | 0.09 | 0.84 | 0.094 | 0.83 | 0.09 | 0.16 | 0.094 | 0.17 | 0.09 | 25 |
| 77 | *Pachydrus obniger* | 11 | 0.86 | 0.099 | 0.75 | 0.13 | 0.73 | 0.149 | 0.76 | 0.115 | 0.27 | 0.149 | 0.24 | 0.115 | 10 |
| 78 | *Pantala flavescens* | 24 | 0.91 | 0.06 | 0.84 | 0.093 | 0.85 | 0.108 | 0.84 | 0.079 | 0.15 | 0.108 | 0.16 | 0.079 | 15 |
| 79 | *Pelocoris poeyi* | 11 | 0.89 | 0.072 | 0.8 | 0.081 | 0.8 | 0.094 | 0.8 | 0.072 | 0.2 | 0.094 | 0.2 | 0.072 | 10 |
| 80 | *Perithemis domitia* | 41 | 0.93 | 0.06 | 0.85 | 0.087 | 0.85 | 0.089 | 0.84 | 0.089 | 0.15 | 0.089 | 0.16 | 0.089 | 25 |
| 81 | *Phylloicus chalybeus* | 17 | 0.87 | 0.091 | 0.77 | 0.091 | 0.76 | 0.094 | 0.78 | 0.1 | 0.24 | 0.094 | 0.22 | 0.1 | 14 |
| 82 | *Physella acuta* | 26 | 0.86 | 0.074 | 0.78 | 0.083 | 0.78 | 0.08 | 0.78 | 0.091 | 0.22 | 0.08 | 0.22 | 0.091 | 15 |
| 83 | *Planorbella duryi* | 10 | 0.9 | 0.066 | 0.82 | 0.108 | 0.82 | 0.114 | 0.81 | 0.11 | 0.18 | 0.114 | 0.19 | 0.11 | 10 |
| 84 | *Pomacea paludosa* | 18 | 0.91 | 0.063 | 0.84 | 0.068 | 0.84 | 0.076 | 0.83 | 0.07 | 0.16 | 0.076 | 0.17 | 0.07 | 15 |
| 85 | *Progomphus integer* | 20 | 0.94 | 0.04 | 0.85 | 0.041 | 0.86 | 0.04 | 0.85 | 0.058 | 0.14 | 0.04 | 0.15 | 0.058 | 15 |
| 86 | *Protoneura capillaris* | 27 | 0.98 | 0.025 | 0.96 | 0.05 | 0.98 | 0.046 | 0.95 | 0.057 | 0.02 | 0.046 | 0.05 | 0.057 | 15 |
| 87 | *Pseudosuccinea columella* | 11 | 0.91 | 0.065 | 0.87 | 0.068 | 0.88 | 0.086 | 0.85 | 0.064 | 0.12 | 0.086 | 0.15 | 0.064 | 10 |
| 88 | *Psilopelmia hematopotum* | 11 | 0.97 | 0.026 | 0.9 | 0.085 | 0.89 | 0.096 | 0.9 | 0.08 | 0.11 | 0.096 | 0.1 | 0.08 | 10 |

| | Species | n_p | AUC_m | AUC_sd | sTSS_m | sTSS_sd | Sen_m | Sen_sd | Spec_m | Spec_sd | Om_m | Om_sd | Com_m | Com_sd | n_f_m |
|---|---|---|---|---|---|---|---|---|---|---|---|---|---|---|---|
| 89 | *Rhagovelia collaris* | 38 | 0.93 | 0.05 | 0.84 | 0.071 | 0.84 | 0.086 | 0.84 | 0.06 | 0.16 | 0.086 | 0.16 | 0.06 | 25 |
| 90 | *Rhionaeschna psilus* | 12 | 0.96 | 0.03 | 0.9 | 0.068 | 0.88 | 0.081 | 0.92 | 0.061 | 0.12 | 0.081 | 0.07 | 0.061 | 10 |
| 91 | *Scapanea frontalis* | 28 | 0.9 | 0.049 | 0.81 | 0.082 | 0.81 | 0.087 | 0.8 | 0.082 | 0.19 | 0.087 | 0.2 | 0.082 | 15 |
| 92 | *Sicydium plumieri* | 17 | 0.93 | 0.059 | 0.88 | 0.101 | 0.88 | 0.106 | 0.88 | 0.102 | 0.12 | 0.106 | 0.12 | 0.102 | 15 |
| 93 | *Smicridea comma* | 59 | 0.94 | 0.037 | 0.86 | 0.061 | 0.86 | 0.068 | 0.86 | 0.057 | 0.14 | 0.068 | 0.14 | 0.057 | 25 |
| 94 | *Tarebia granifera* | 45 | 0.93 | 0.044 | 0.87 | 0.07 | 0.88 | 0.071 | 0.86 | 0.072 | 0.12 | 0.071 | 0.14 | 0.072 | 25 |
| 95 | *Telebasis dominicana* | 23 | 0.82 | 0.096 | 0.77 | 0.094 | 0.77 | 0.102 | 0.76 | 0.093 | 0.23 | 0.102 | 0.24 | 0.093 | 15 |
| 96 | *Thermonectus basillaris* | 11 | 0.85 | 0.077 | 0.73 | 0.106 | 0.73 | 0.115 | 0.73 | 0.102 | 0.27 | 0.115 | 0.27 | 0.102 | 10 |
| 97 | *Thermonectus circumscriptus* | 10 | 0.84 | 0.051 | 0.77 | 0.064 | 0.8 | 0.094 | 0.75 | 0.063 | 0.2 | 0.094 | 0.25 | 0.063 | 10 |
| 98 | *Thermonectus succinctus* | 10 | 0.9 | 0.058 | 0.81 | 0.11 | 0.8 | 0.133 | 0.82 | 0.092 | 0.2 | 0.133 | 0.18 | 0.092 | 10 |
| 99 | *Tramea abdominalis* | 25 | 0.91 | 0.061 | 0.84 | 0.084 | 0.85 | 0.086 | 0.84 | 0.084 | 0.15 | 0.086 | 0.16 | 0.084 | 15 |
| 100 | *Tramea onusta* | 12 | 0.88 | 0.095 | 0.81 | 0.125 | 0.8 | 0.131 | 0.82 | 0.121 | 0.2 | 0.131 | 0.17 | 0.121 | 10 |
| 101 | *Traverina oriente* | 19 | 0.9 | 0.07 | 0.81 | 0.103 | 0.81 | 0.112 | 0.8 | 0.105 | 0.19 | 0.112 | 0.2 | 0.105 | 15 |
| 102 | *Tricorythodes cubensis* | 17 | 0.8 | 0.109 | 0.75 | 0.089 | 0.76 | 0.087 | 0.74 | 0.101 | 0.24 | 0.087 | 0.26 | 0.101 | 14 |
| 103 | *Tricorythodes grallator* | 30 | 0.94 | 0.07 | 0.89 | 0.086 | 0.87 | 0.087 | 0.9 | 0.096 | 0.13 | 0.087 | 0.1 | 0.096 | 25 |
| 104 | *Tricorythodes montanus* | 14 | 0.96 | 0.041 | 0.92 | 0.072 | 0.93 | 0.075 | 0.92 | 0.071 | 0.07 | 0.075 | 0.08 | 0.071 | 10 |
| 105 | *Tricorythodes sacculobranchis* | 44 | 0.93 | 0.036 | 0.86 | 0.066 | 0.86 | 0.073 | 0.85 | 0.065 | 0.14 | 0.073 | 0.15 | 0.065 | 25 |
| 106 | *Xiphocaris elongata* | 10 | 0.9 | 0.082 | 0.79 | 0.126 | 0.8 | 0.133 | 0.78 | 0.123 | 0.2 | 0.133 | 0.22 | 0.123 | 10 |

Table S1.3: Mean (m) and standard deviation (sd) of performance metrics for ensemble Maxent species distribution models fitted for freshwater species of Cuba. sTSS: standardized True Skill Statistic; AUC: Area Under the Receiver Operating Characteristic Curve; Sen: sensitivity; Spec: specificity; Om: omission rate; Com: commission rate; n_p: number of presences used to fit the models; n_f_m: number of models fitted across species-specific cross-validation runs.

| | Species | n_p | AUC_m | AUC_sd | sTSS_m | sTSS_sd | Sen_m | Sen_sd | Spec_m | Spec_sd | Om_m | Om_sd | Com_m | Com_sd | n_f_m |
|---|---|---|---|---|---|---|---|---|---|---|---|---|---|---|---|
| 1 | *Agonostomus monticola* | 24 | 0.84 | 0.076 | 0.78 | 0.097 | 0.79 | 0.102 | 0.77 | 0.096 | 0.21 | 0.102 | 0.23 | 0.096 | 15 |
| 2 | *Alepidomus evermanni* | 12 | 0.93 | 0.062 | 0.89 | 0.063 | 0.88 | 0.081 | 0.9 | 0.066 | 0.12 | 0.081 | 0.1 | 0.066 | 10 |
| 3 | *Anguilla rostrata* | 17 | 0.94 | 0.077 | 0.89 | 0.091 | 0.89 | 0.103 | 0.9 | 0.088 | 0.11 | 0.103 | 0.1 | 0.088 | 15 |
| 4 | *Awaous banana* | 12 | 0.87 | 0.095 | 0.83 | 0.087 | 0.83 | 0.083 | 0.82 | 0.097 | 0.17 | 0.083 | 0.18 | 0.097 | 9 |
| 5 | *Berosus interstitialis* | 10 | 0.8 | 0.073 | 0.75 | 0.1 | 0.77 | 0.151 | 0.73 | 0.082 | 0.23 | 0.151 | 0.27 | 0.082 | 6 |
| 6 | *Berosus trilobus* | 31 | 0.98 | 0.028 | 0.93 | 0.075 | 0.92 | 0.082 | 0.93 | 0.071 | 0.08 | 0.082 | 0.07 | 0.071 | 25 |
| 7 | *Bidessonotus browneanus* | 24 | 0.83 | 0.091 | 0.77 | 0.091 | 0.77 | 0.093 | 0.77 | 0.091 | 0.23 | 0.093 | 0.23 | 0.091 | 15 |
| 8 | *Brachymesia herbida* | 15 | 0.87 | 0.087 | 0.79 | 0.127 | 0.8 | 0.151 | 0.78 | 0.115 | 0.2 | 0.151 | 0.22 | 0.115 | 15 |
| 9 | *Caenis cubensis* | 53 | 0.81 | 0.07 | 0.75 | 0.068 | 0.75 | 0.069 | 0.75 | 0.072 | 0.25 | 0.069 | 0.25 | 0.072 | 25 |
| 10 | *Callibaetis floridanus* | 23 | 0.72 | 0.067 | 0.64 | 0.082 | 0.63 | 0.085 | 0.64 | 0.098 | 0.37 | 0.085 | 0.36 | 0.098 | 15 |
| 11 | *Cannaphila insularis* | 11 | 0.86 | 0.101 | 0.79 | 0.135 | 0.81 | 0.155 | 0.77 | 0.123 | 0.19 | 0.155 | 0.23 | 0.123 | 10 |
| 12 | *Careospina baconaoi* | 20 | 0.74 | 0.12 | 0.72 | 0.128 | 0.73 | 0.136 | 0.72 | 0.124 | 0.27 | 0.136 | 0.28 | 0.124 | 15 |
| 13 | *Careospina hespera* | 49 | 0.81 | 0.071 | 0.73 | 0.079 | 0.73 | 0.081 | 0.73 | 0.08 | 0.27 | 0.081 | 0.27 | 0.08 | 25 |
| 14 | *Caribaetis planifrons* | 74 | 0.89 | 0.06 | 0.83 | 0.079 | 0.84 | 0.079 | 0.83 | 0.083 | 0.16 | 0.079 | 0.17 | 0.083 | 25 |
| 15 | *Chimarra guapa* | 10 | 0.72 | 0.103 | 0.67 | 0.087 | 0.67 | 0.1 | 0.68 | 0.083 | 0.33 | 0.1 | 0.32 | 0.083 | 9 |
| 16 | *Chimarra pulchra* | 14 | 0.69 | 0.117 | 0.69 | 0.118 | 0.69 | 0.129 | 0.68 | 0.116 | 0.31 | 0.129 | 0.32 | 0.116 | 7 |
| 17 | *Cloeodes inferior* | 18 | 0.9 | 0.071 | 0.82 | 0.095 | 0.81 | 0.107 | 0.83 | 0.097 | 0.19 | 0.107 | 0.17 | 0.097 | 15 |
| 18 | *Cloeodes superior* | 17 | 0.96 | 0.06 | 0.92 | 0.104 | 0.92 | 0.107 | 0.92 | 0.106 | 0.08 | 0.107 | 0.08 | 0.106 | 15 |
| 19 | *Copelatus posticatus* | 25 | 0.85 | 0.099 | 0.75 | 0.125 | 0.76 | 0.141 | 0.75 | 0.114 | 0.24 | 0.141 | 0.25 | 0.114 | 15 |
| 20 | *Crocothemis servilia* | 39 | 0.93 | 0.047 | 0.84 | 0.076 | 0.85 | 0.082 | 0.84 | 0.074 | 0.15 | 0.082 | 0.16 | 0.074 | 25 |
| 21 | *Cubanichthys cubensis* | 11 | 0.85 | 0.072 | 0.79 | 0.085 | 0.78 | 0.114 | 0.79 | 0.075 | 0.22 | 0.114 | 0.21 | 0.075 | 10 |
| 22 | *Cubanoptila purpurea* | 19 | 0.86 | 0.075 | 0.82 | 0.069 | 0.82 | 0.077 | 0.82 | 0.069 | 0.18 | 0.077 | 0.18 | 0.069 | 15 |
| 23 | *Dineutus longimanus* | 24 | 0.94 | 0.063 | 0.87 | 0.068 | 0.88 | 0.074 | 0.86 | 0.069 | 0.12 | 0.074 | 0.14 | 0.069 | 15 |
| 24 | *Dythemis rufinervis* | 22 | 0.83 | 0.06 | 0.78 | 0.069 | 0.79 | 0.074 | 0.77 | 0.076 | 0.21 | 0.074 | 0.23 | 0.076 | 15 |
| 25 | *Dythemis sterilis* | 23 | 0.83 | 0.078 | 0.75 | 0.092 | 0.75 | 0.097 | 0.75 | 0.091 | 0.25 | 0.097 | 0.25 | 0.091 | 15 |
| 26 | *Eleotris pisonis* | 12 | 0.82 | 0.043 | 0.78 | 0.065 | 0.8 | 0.07 | 0.77 | 0.066 | 0.2 | 0.07 | 0.23 | 0.066 | 10 |
| 27 | *Enallagma civile* | 19 | 0.84 | 0.127 | 0.8 | 0.112 | 0.8 | 0.114 | 0.79 | 0.113 | 0.2 | 0.114 | 0.21 | 0.113 | 13 |

| | Species | n_p | AUC_m | AUC_sd | sTSS_m | sTSS_sd | Sen_m | Sen_sd | Spec_m | Spec_sd | Om_m | Om_sd | Com_m | Com_sd | n_f_m |
|---|---|---|---|---|---|---|---|---|---|---|---|---|---|---|---|
| 28 | *Enallagma coecum* | 49 | 0.9 | 0.073 | 0.82 | 0.072 | 0.82 | 0.073 | 0.82 | 0.074 | 0.18 | 0.073 | 0.18 | 0.074 | 25 |
| 29 | *Epilobocera cubensis* | 26 | 0.77 | 0.11 | 0.71 | 0.122 | 0.71 | 0.134 | 0.72 | 0.113 | 0.29 | 0.134 | 0.28 | 0.113 | 15 |
| 30 | *Erythemis plebeja* | 19 | 0.84 | 0.082 | 0.76 | 0.108 | 0.76 | 0.127 | 0.75 | 0.096 | 0.24 | 0.127 | 0.25 | 0.096 | 15 |
| 31 | *Erythemis simplicicollis* | 10 | 0.76 | 0.088 | 0.68 | 0.083 | 0.67 | 0.1 | 0.69 | 0.078 | 0.33 | 0.1 | 0.31 | 0.078 | 9 |
| 32 | *Erythemis vesiculosa* | 26 | 0.88 | 0.05 | 0.83 | 0.079 | 0.84 | 0.095 | 0.83 | 0.067 | 0.16 | 0.095 | 0.17 | 0.067 | 15 |
| 33 | *Erythrodiplax berenice* | 21 | 0.86 | 0.085 | 0.76 | 0.083 | 0.76 | 0.088 | 0.77 | 0.081 | 0.24 | 0.088 | 0.23 | 0.081 | 15 |
| 34 | *Erythrodiplax fervida* | 21 | 0.82 | 0.1 | 0.78 | 0.094 | 0.8 | 0.108 | 0.76 | 0.087 | 0.2 | 0.108 | 0.24 | 0.087 | 14 |
| 35 | *Erythrodiplax justiniana* | 27 | 0.92 | 0.086 | 0.84 | 0.106 | 0.84 | 0.117 | 0.84 | 0.1 | 0.16 | 0.117 | 0.16 | 0.1 | 15 |
| 36 | *Erythrodiplax umbrata* | 53 | 0.91 | 0.045 | 0.84 | 0.073 | 0.85 | 0.07 | 0.84 | 0.077 | 0.15 | 0.07 | 0.16 | 0.077 | 25 |
| 37 | *Fallceon poeyi* | 28 | 0.85 | 0.062 | 0.76 | 0.078 | 0.76 | 0.085 | 0.75 | 0.075 | 0.24 | 0.085 | 0.25 | 0.075 | 15 |
| 38 | *Farrodes bimaculatus* | 42 | 0.81 | 0.083 | 0.73 | 0.091 | 0.73 | 0.097 | 0.72 | 0.089 | 0.27 | 0.097 | 0.28 | 0.089 | 24 |
| 39 | *Galba cubensis* | 17 | 0.84 | 0.111 | 0.78 | 0.12 | 0.78 | 0.151 | 0.79 | 0.1 | 0.22 | 0.151 | 0.21 | 0.1 | 15 |
| 40 | *Gambusia punctata* | 49 | 0.77 | 0.085 | 0.72 | 0.088 | 0.74 | 0.088 | 0.71 | 0.092 | 0.26 | 0.088 | 0.29 | 0.092 | 25 |
| 41 | *Gambusia puncticulata* | 49 | 0.95 | 0.037 | 0.89 | 0.061 | 0.89 | 0.061 | 0.89 | 0.063 | 0.11 | 0.061 | 0.11 | 0.063 | 25 |
| 42 | *Girardinus denticulatus* | 16 | 0.67 | 0.093 | 0.65 | 0.071 | 0.68 | 0.119 | 0.62 | 0.125 | 0.32 | 0.119 | 0.38 | 0.125 | 13 |
| 43 | *Girardinus metallicus* | 31 | 0.88 | 0.073 | 0.81 | 0.077 | 0.81 | 0.08 | 0.82 | 0.079 | 0.19 | 0.08 | 0.18 | 0.079 | 25 |
| 44 | *Gundlachia radiata* | 12 | 0.85 | 0.08 | 0.83 | 0.099 | 0.87 | 0.105 | 0.79 | 0.113 | 0.13 | 0.105 | 0.21 | 0.113 | 10 |
| 45 | *Hagenulus morrisonae* | 44 | 0.78 | 0.078 | 0.71 | 0.084 | 0.71 | 0.08 | 0.7 | 0.093 | 0.29 | 0.08 | 0.3 | 0.093 | 25 |
| 46 | *Helicopsyche comosa* | 17 | 0.85 | 0.072 | 0.83 | 0.078 | 0.84 | 0.1 | 0.82 | 0.064 | 0.16 | 0.1 | 0.18 | 0.064 | 15 |
| 47 | *Helicopsyche hageni* | 13 | 0.76 | 0.124 | 0.75 | 0.115 | 0.76 | 0.106 | 0.74 | 0.128 | 0.24 | 0.106 | 0.26 | 0.128 | 9 |
| 48 | *Hydropsyche cubana* | 23 | 0.84 | 0.09 | 0.79 | 0.099 | 0.8 | 0.1 | 0.77 | 0.103 | 0.2 | 0.1 | 0.23 | 0.103 | 15 |
| 49 | *Hydrovatus caraibus* | 18 | 0.83 | 0.076 | 0.77 | 0.086 | 0.78 | 0.081 | 0.77 | 0.096 | 0.22 | 0.081 | 0.23 | 0.096 | 15 |
| 50 | *Hypolestes trinitatis* | 34 | 0.9 | 0.062 | 0.82 | 0.069 | 0.82 | 0.074 | 0.82 | 0.071 | 0.18 | 0.074 | 0.18 | 0.071 | 25 |
| 51 | *Ischnura capreolus* | 13 | 0.83 | 0.043 | 0.72 | 0.049 | 0.71 | 0.051 | 0.74 | 0.056 | 0.29 | 0.051 | 0.26 | 0.056 | 9 |
| 52 | *Ischnura ramburii* | 40 | 0.96 | 0.034 | 0.9 | 0.06 | 0.91 | 0.068 | 0.9 | 0.058 | 0.09 | 0.068 | 0.1 | 0.058 | 25 |
| 53 | *Joturus pichardi* | 14 | 0.74 | 0.07 | 0.75 | 0.068 | 0.78 | 0.104 | 0.71 | 0.051 | 0.22 | 0.104 | 0.29 | 0.051 | 9 |
| 54 | *Laccophilus bifasciatus* | 12 | 0.87 | 0.079 | 0.85 | 0.093 | 0.88 | 0.112 | 0.82 | 0.107 | 0.12 | 0.112 | 0.17 | 0.107 | 10 |
| 55 | *Laccophilus gentilis* | 11 | 0.76 | 0.074 | 0.7 | 0.101 | 0.7 | 0.102 | 0.71 | 0.103 | 0.3 | 0.102 | 0.29 | 0.103 | 10 |
| 56 | *Laccophilus proximus* | 30 | 0.84 | 0.085 | 0.76 | 0.111 | 0.75 | 0.128 | 0.76 | 0.098 | 0.25 | 0.128 | 0.24 | 0.098 | 25 |
| 57 | *Laccophilus venustus* | 22 | 0.91 | 0.053 | 0.84 | 0.075 | 0.85 | 0.082 | 0.84 | 0.073 | 0.15 | 0.082 | 0.16 | 0.073 | 15 |
| 58 | *Limia vittata* | 43 | 0.92 | 0.054 | 0.85 | 0.071 | 0.86 | 0.077 | 0.85 | 0.066 | 0.14 | 0.077 | 0.15 | 0.066 | 25 |

| | Species | n_p | AUC_m | AUC_sd | sTSS_m | sTSS_sd | Sen_m | Sen_sd | Spec_m | Spec_sd | Om_m | Om_sd | Com_m | Com_sd | n_f_m |
|---|---|---|---|---|---|---|---|---|---|---|---|---|---|---|---|
| 59 | *Limnogonus franciscanus* | 22 | 0.9 | 0.061 | 0.81 | 0.104 | 0.83 | 0.111 | 0.8 | 0.101 | 0.17 | 0.111 | 0.2 | 0.101 | 15 |
| 60 | *Ludwigia erecta* | 11 | 0.73 | 0.084 | 0.69 | 0.098 | 0.71 | 0.133 | 0.68 | 0.08 | 0.29 | 0.133 | 0.32 | 0.08 | 9 |
| 61 | *Ludwigia peploides* | 10 | 0.83 | 0.136 | 0.75 | 0.19 | 0.78 | 0.186 | 0.72 | 0.199 | 0.22 | 0.186 | 0.28 | 0.199 | 9 |
| 62 | *Macrobrachium acanthurus* | 12 | 0.83 | 0.13 | 0.78 | 0.09 | 0.78 | 0.081 | 0.77 | 0.11 | 0.22 | 0.081 | 0.23 | 0.11 | 10 |
| 63 | *Macrobrachium faustinum* | 11 | 0.91 | 0.069 | 0.85 | 0.072 | 0.85 | 0.079 | 0.85 | 0.077 | 0.15 | 0.079 | 0.15 | 0.077 | 10 |
| 64 | *Macrothemis celeno* | 51 | 0.82 | 0.079 | 0.75 | 0.064 | 0.76 | 0.065 | 0.75 | 0.068 | 0.24 | 0.065 | 0.25 | 0.068 | 25 |
| 65 | *Meridiorhantus calidus* | 10 | 0.88 | 0.105 | 0.82 | 0.118 | 0.82 | 0.148 | 0.82 | 0.092 | 0.18 | 0.148 | 0.18 | 0.092 | 10 |
| 66 | *Mesovelia mulsanti* | 10 | 0.87 | 0.083 | 0.78 | 0.085 | 0.78 | 0.114 | 0.79 | 0.057 | 0.22 | 0.114 | 0.21 | 0.057 | 10 |
| 67 | *Miathyria marcella* | 14 | 0.78 | 0.115 | 0.71 | 0.115 | 0.73 | 0.105 | 0.69 | 0.127 | 0.27 | 0.105 | 0.31 | 0.127 | 10 |
| 68 | *Micrathyria aequalis* | 10 | 0.83 | 0.114 | 0.77 | 0.125 | 0.8 | 0.141 | 0.74 | 0.113 | 0.2 | 0.141 | 0.26 | 0.113 | 9 |
| 69 | *Micrathyria hagenii* | 10 | 0.73 | 0.138 | 0.68 | 0.112 | 0.71 | 0.105 | 0.64 | 0.142 | 0.29 | 0.105 | 0.36 | 0.142 | 9 |
| 70 | *Microvelia cubana* | 10 | 0.89 | 0.087 | 0.79 | 0.115 | 0.78 | 0.148 | 0.8 | 0.094 | 0.22 | 0.148 | 0.2 | 0.094 | 10 |
| 71 | *Nandopsis ramsdeni* | 11 | 0.85 | 0.103 | 0.82 | 0.064 | 0.82 | 0.096 | 0.83 | 0.052 | 0.18 | 0.096 | 0.17 | 0.052 | 10 |
| 72 | *Nandopsis tetracanthus* | 37 | 0.75 | 0.11 | 0.69 | 0.115 | 0.69 | 0.121 | 0.69 | 0.116 | 0.31 | 0.121 | 0.31 | 0.116 | 25 |
| 73 | *Nectopsyche cubana* | 11 | 0.65 | 0.118 | 0.64 | 0.101 | 0.66 | 0.109 | 0.62 | 0.121 | 0.34 | 0.109 | 0.38 | 0.121 | 6 |
| 74 | *Neoneura maria* | 50 | 0.76 | 0.081 | 0.7 | 0.091 | 0.7 | 0.096 | 0.69 | 0.092 | 0.3 | 0.096 | 0.31 | 0.092 | 25 |
| 75 | *Notonecta indica* | 10 | 0.7 | 0.107 | 0.66 | 0.086 | 0.68 | 0.104 | 0.64 | 0.119 | 0.32 | 0.104 | 0.36 | 0.119 | 8 |
| 76 | *Orthemis ferruginea* | 37 | 0.93 | 0.055 | 0.85 | 0.099 | 0.86 | 0.104 | 0.85 | 0.097 | 0.14 | 0.104 | 0.15 | 0.097 | 25 |
| 77 | *Pachydrus obniger* | 11 | 0.93 | 0.09 | 0.88 | 0.116 | 0.9 | 0.119 | 0.87 | 0.115 | 0.1 | 0.119 | 0.13 | 0.115 | 10 |
| 78 | *Pantala flavescens* | 24 | 0.87 | 0.082 | 0.78 | 0.091 | 0.79 | 0.09 | 0.77 | 0.096 | 0.21 | 0.09 | 0.23 | 0.096 | 15 |
| 79 | *Pelocoris poeyi* | 11 | 0.87 | 0.081 | 0.83 | 0.084 | 0.82 | 0.112 | 0.83 | 0.09 | 0.18 | 0.112 | 0.17 | 0.09 | 10 |
| 80 | *Perithemis domitia* | 41 | 0.88 | 0.073 | 0.83 | 0.072 | 0.84 | 0.075 | 0.82 | 0.075 | 0.16 | 0.075 | 0.18 | 0.075 | 25 |
| 81 | *Phylloicus chalybeus* | 17 | 0.68 | 0.091 | 0.65 | 0.054 | 0.69 | 0.115 | 0.61 | 0.126 | 0.31 | 0.115 | 0.39 | 0.126 | 12 |
| 82 | *Physella acuta* | 26 | 0.78 | 0.09 | 0.73 | 0.069 | 0.74 | 0.059 | 0.73 | 0.086 | 0.26 | 0.059 | 0.27 | 0.086 | 15 |
| 83 | *Planorbella duryi* | 10 | 0.9 | 0.08 | 0.86 | 0.134 | 0.86 | 0.165 | 0.85 | 0.108 | 0.14 | 0.165 | 0.15 | 0.108 | 10 |
| 84 | *Pomacea paludosa* | 18 | 0.86 | 0.057 | 0.82 | 0.098 | 0.82 | 0.133 | 0.82 | 0.085 | 0.18 | 0.133 | 0.18 | 0.085 | 15 |
| 85 | *Progomphus integer* | 20 | 0.77 | 0.065 | 0.73 | 0.094 | 0.75 | 0.116 | 0.71 | 0.085 | 0.25 | 0.116 | 0.29 | 0.085 | 15 |
| 86 | *Protoneura capillaris* | 27 | 0.94 | 0.039 | 0.88 | 0.088 | 0.88 | 0.098 | 0.87 | 0.08 | 0.12 | 0.098 | 0.13 | 0.08 | 15 |
| 87 | *Pseudosuccinea columella* | 11 | 0.93 | 0.064 | 0.88 | 0.108 | 0.87 | 0.118 | 0.88 | 0.105 | 0.13 | 0.118 | 0.12 | 0.105 | 10 |
| 88 | *Psilopelmia hematopotum* | 11 | 0.84 | 0.109 | 0.78 | 0.1 | 0.8 | 0.107 | 0.76 | 0.123 | 0.2 | 0.107 | 0.24 | 0.123 | 10 |
| 89 | *Rhagovelia collaris* | 38 | 0.81 | 0.068 | 0.75 | 0.077 | 0.75 | 0.081 | 0.75 | 0.078 | 0.25 | 0.081 | 0.25 | 0.078 | 25 |

| | Species | n_p | AUC_m | AUC_sd | sTSS_m | sTSS_sd | Sen_m | Sen_sd | Spec_m | Spec_sd | Om_m | Om_sd | Com_m | Com_sd | n_f_m |
|---|---|---|---|---|---|---|---|---|---|---|---|---|---|---|---|
| 90 | *Rhionaeschna psilus* | 12 | 0.78 | 0.059 | 0.72 | 0.088 | 0.7 | 0.139 | 0.73 | 0.056 | 0.3 | 0.139 | 0.27 | 0.056 | 9 |
| 91 | *Scapanea frontalis* | 28 | 0.9 | 0.048 | 0.8 | 0.082 | 0.81 | 0.096 | 0.8 | 0.071 | 0.19 | 0.096 | 0.2 | 0.071 | 15 |
| 92 | *Sicydium plumieri* | 17 | 0.86 | 0.065 | 0.81 | 0.079 | 0.81 | 0.087 | 0.81 | 0.084 | 0.19 | 0.087 | 0.19 | 0.084 | 15 |
| 93 | *Smicridea comma* | 59 | 0.8 | 0.067 | 0.73 | 0.072 | 0.73 | 0.083 | 0.72 | 0.072 | 0.27 | 0.083 | 0.28 | 0.072 | 25 |
| 94 | *Tarebia granifera* | 45 | 0.84 | 0.074 | 0.78 | 0.084 | 0.8 | 0.089 | 0.77 | 0.083 | 0.2 | 0.089 | 0.23 | 0.083 | 25 |
| 95 | *Telebasis dominicana* | 23 | 0.85 | 0.066 | 0.78 | 0.088 | 0.78 | 0.103 | 0.77 | 0.081 | 0.22 | 0.103 | 0.23 | 0.081 | 15 |
| 96 | *Thermonectus basillaris* | 11 | 0.86 | 0.095 | 0.78 | 0.093 | 0.78 | 0.159 | 0.77 | 0.11 | 0.22 | 0.159 | 0.23 | 0.11 | 10 |
| 97 | *Thermonectus circumscriptus* | 10 | 0.82 | 0.144 | 0.78 | 0.162 | 0.8 | 0.2 | 0.76 | 0.136 | 0.2 | 0.2 | 0.24 | 0.136 | 9 |
| 98 | *Thermonectus succinctus* | 10 | 0.7 | 0.107 | 0.64 | 0.088 | 0.67 | 0.141 | 0.62 | 0.097 | 0.33 | 0.141 | 0.38 | 0.097 | 9 |
| 99 | *Tramea abdominalis* | 25 | 0.84 | 0.059 | 0.77 | 0.07 | 0.78 | 0.079 | 0.76 | 0.067 | 0.22 | 0.079 | 0.24 | 0.067 | 15 |
| 100 | *Tramea onusta* | 12 | 0.81 | 0.126 | 0.77 | 0.141 | 0.78 | 0.137 | 0.76 | 0.149 | 0.22 | 0.137 | 0.24 | 0.149 | 10 |
| 101 | *Traverina oriente* | 19 | 0.72 | 0.078 | 0.69 | 0.082 | 0.7 | 0.107 | 0.68 | 0.069 | 0.3 | 0.107 | 0.32 | 0.069 | 14 |
| 102 | *Tricorythodes cubensis* | 17 | 0.6 | 0.061 | 0.61 | 0.055 | 0.66 | 0.104 | 0.56 | 0.076 | 0.34 | 0.104 | 0.44 | 0.076 | 10 |
| 103 | *Tricorythodes grallator* | 30 | 0.87 | 0.116 | 0.81 | 0.101 | 0.81 | 0.092 | 0.81 | 0.117 | 0.19 | 0.092 | 0.19 | 0.117 | 25 |
| 104 | *Tricorythodes montanus* | 14 | 0.88 | 0.078 | 0.82 | 0.106 | 0.83 | 0.113 | 0.81 | 0.102 | 0.17 | 0.113 | 0.19 | 0.102 | 10 |
| 105 | *Tricorythodes sacculobranchis* | 44 | 0.78 | 0.068 | 0.72 | 0.08 | 0.73 | 0.087 | 0.72 | 0.077 | 0.27 | 0.087 | 0.28 | 0.077 | 25 |
| 106 | *Xiphocaris elongata* | 10 | 0.85 | 0.092 | 0.76 | 0.109 | 0.76 | 0.126 | 0.75 | 0.097 | 0.24 | 0.126 | 0.25 | 0.097 | 10 |

Table S1.4. Species-level representation under the current protected-area system and prioritization scenarios. For each species, the table reports the percentage of its conservation target represented by the current Cuban protected-area system (SNAP), the free-choice prioritization solution, and the lock-in prioritization solution. The columns Met target SNAP and Met target solution indicate whether the conservation target was achieved under SNAP and under the prioritization solution, respectively.

| species | Relative protection SNAP | Met target SNAP | Relative protection free-choice | Relative protection lock-in | Met target solution | Endemic status | IUCN category |
|---|---|---|---|---|---|---|---|
| *Agonostomus monticola* | 34 | TRUE | 37 | 48 | TRUE | Non-endemic | NE |
| *Alepidomus evermanni* | 57 | TRUE | 45 | 66 | TRUE | Endemic | LC |
| *Alisotrichia alayoana* | 23 | FALSE | 78 | 96 | TRUE | Endemic | NE |
| *Alisotrichia chiquitica* | 0 | FALSE | 56 | 96 | TRUE | Endemic | NE |
| *Alisotrichia flintiana* | 0 | FALSE | 100 | 100 | TRUE | Endemic | NE |
| *Americabaetis naranjoi* | 17 | FALSE | 43 | 52 | TRUE | Endemic | NE |
| *Anax amazili* | 0 | FALSE | 48 | 48 | TRUE | Non-endemic | LC |
| *Anax junius* | 47 | TRUE | 77 | 80 | TRUE | Non-endemic | LC |
| *Anguilla rostrata* | 33 | TRUE | 38 | 49 | TRUE | Non-endemic | EN |
| *Anodocheilus exiguus* | 47 | TRUE | 44 | 77 | TRUE | Non-endemic | NE |
| *Antillopsyche wrighti* | 0 | FALSE | 100 | 100 | TRUE | Endemic | NE |
| *Aphylla caraiba* | 0 | FALSE | 69 | 69 | TRUE | Non-endemic | LC |
| *Atopsyche cubana* | 24 | FALSE | 100 | 100 | TRUE | Endemic | NE |
| *Atopsyche vinai* | 35 | TRUE | 98 | 93 | TRUE | Endemic | NE |
| *Atya innocous* | 30 | TRUE | 36 | 97 | TRUE | Non-endemic | LC |
| *Atya lanipes* | 0 | FALSE | 35 | 54 | TRUE | Non-endemic | LC |
| *Austrotinodes cubanus* | 49 | TRUE | 98 | 98 | TRUE | Endemic | NE |
| *Awaous banana* | 32 | TRUE | 42 | 50 | TRUE | Non-endemic | LC |
| *Azolla caroliniana* | 82 | TRUE | 78 | 82 | TRUE | Non-endemic | NE |
| *Belostoma minor* | 0 | FALSE | 78 | 78 | TRUE | Non-endemic | NE |
| *Berosus chevrolati* | 30 | TRUE | 41 | 72 | TRUE | Endemic | NE |
| *Berosus exiguus* | 74 | TRUE | 40 | 74 | TRUE | Non-endemic | NE |
| *Berosus infuscatus* | 16 | FALSE | 77 | 93 | TRUE | Non-endemic | NE |
| *Berosus interstitialis* | 44 | TRUE | 37 | 56 | TRUE | Non-endemic | NE |
| *Berosus quadridens* | 18 | FALSE | 82 | 100 | TRUE | Non-endemic | NE |
| *Berosus trilobus* | 29 | FALSE | 36 | 46 | TRUE | Non-endemic | NE |
| *Berosus undatus* | 8 | FALSE | 97 | 97 | TRUE | Non-endemic | NE |
| *Bidessonotus browneanus* | 26 | FALSE | 33 | 44 | TRUE | Non-endemic | NE |

| species | Relative protection SNAP | Met target SNAP | Relative protection free-choice | Relative protection lock-in | Met target solution | Endemic status | IUCN category |
|---|---|---|---|---|---|---|---|
| *Biomphalaria havanensis* | 0 | FALSE | 100 | 100 | TRUE | Non-endemic | LC |
| *Biomphalaria helophila* | 36 | TRUE | 60 | 61 | TRUE | Non-endemic | LC |
| *Biomphalaria pallida* | 0 | FALSE | 100 | 100 | TRUE | Non-endemic | NE |
| *Borinquena sexta* | 100 | TRUE | 87 | 100 | TRUE | Endemic | NE |
| *Brachymesia furcata* | 83 | TRUE | 99 | 99 | TRUE | Non-endemic | LC |
| *Brachymesia herbida* | 37 | TRUE | 35 | 51 | TRUE | Non-endemic | LC |
| *Buenoa antigone* | 100 | TRUE | 84 | 100 | TRUE | Non-endemic | NE |
| *Buenoa gracilis* | 0 | FALSE | 100 | 100 | TRUE | Non-endemic | NE |
| *Buenoa platycnemis* | 100 | TRUE | 84 | 100 | TRUE | Non-endemic | NE |
| *Caenis cubensis* | 10 | FALSE | 37 | 38 | TRUE | Endemic | NE |
| *Callibaetis floridanus* | 7 | FALSE | 36 | 38 | TRUE | Non-endemic | NE |
| *Campsiophora mulata* | 38 | TRUE | 60 | 98 | TRUE | Endemic | NE |
| *Cannaphila insularis* | 29 | FALSE | 32 | 43 | TRUE | Non-endemic | LC |
| *Careospina baconaoi* | 28 | FALSE | 35 | 42 | TRUE | Endemic | NE |
| *Careospina hespera* | 13 | FALSE | 30 | 30 | TRUE | Endemic | NE |
| *Careospina sierramaestrae* | 0 | FALSE | 92 | 92 | TRUE | Endemic | NE |
| *Caribaetis alcarrazae* | 0 | FALSE | 45 | 45 | TRUE | Endemic | NE |
| *Caribaetis planifrons* | 15 | FALSE | 30 | 34 | TRUE | Non-endemic | NE |
| *Cariboptila poquita* | 100 | TRUE | 36 | 100 | TRUE | Endemic | NE |
| *Cariboptila soltera* | 100 | TRUE | 100 | 100 | TRUE | Endemic | NE |
| *Celina imitatrix* | 0 | FALSE | 53 | 53 | TRUE | Non-endemic | NE |
| *Celina palustris* | 0 | FALSE | 100 | 100 | TRUE | Non-endemic | NE |
| *Celina slossoni* | 0 | FALSE | 49 | 49 | TRUE | Non-endemic | NE |
| *Cercyon insularis* | 18 | FALSE | 82 | 100 | TRUE | Non-endemic | NE |
| *Chimarra guapa* | 26 | FALSE | 50 | 55 | TRUE | Endemic | NE |
| *Chimarra moesta* | 100 | TRUE | 60 | 100 | TRUE | Endemic | NE |
| *Chimarra pulchra* | 0 | FALSE | 86 | 78 | TRUE | Endemic | NE |
| *Chimarra quina* | 100 | TRUE | 100 | 100 | TRUE | Endemic | NE |
| *Cladopelma forcipis* | 0 | FALSE | 80 | 80 | TRUE | Non-endemic | NE |
| *Cloeodes inferior* | 34 | TRUE | 37 | 51 | TRUE | Non-endemic | NE |
| *Cloeodes superior* | 33 | TRUE | 34 | 47 | TRUE | Non-endemic | NE |
| *Coelotanypus cletis* | 0 | FALSE | 100 | 100 | TRUE | Non-endemic | NE |
| *Coelotanypus scapularis* | 100 | TRUE | 95 | 100 | TRUE | Non-endemic | NE |

| species | Relative protection SNAP | Met target SNAP | Relative protection free-choice | Relative protection lock-in | Met target solution | Endemic status | IUCN category |
|---|---|---|---|---|---|---|---|
| *Copelatus caelatipennis* | 0 | FALSE | 52 | 52 | TRUE | Non-endemic | NE |
| *Copelatus cordovai* | 0 | FALSE | 93 | 93 | TRUE | Endemic | NE |
| *Copelatus cubaensis* | 0 | FALSE | 100 | 100 | TRUE | Non-endemic | NE |
| *Copelatus insolitus* | 53 | TRUE | 64 | 94 | TRUE | Non-endemic | NE |
| *Copelatus posticatus* | 23 | FALSE | 36 | 43 | TRUE | Non-endemic | NE |
| *Corbicula fluminea* | 0 | FALSE | 49 | 47 | TRUE | Non-endemic | LC |
| *Corisella edulis* | 0 | FALSE | 98 | 98 | TRUE | Non-endemic | NE |
| *Coryphaeschna adnexa* | 0 | FALSE | 36 | 43 | TRUE | Non-endemic | LC |
| *Coryphaeschna viriditas* | 58 | TRUE | 100 | 100 | TRUE | Non-endemic | LC |
| *Crocothemis servilia* | 37 | TRUE | 38 | 51 | TRUE | Non-endemic | LC |
| *Cubanichthys cubensis* | 46 | TRUE | 35 | 55 | TRUE | Endemic | LC |
| *Cubanoptila botosaneanui* | 100 | TRUE | 100 | 100 | TRUE | Endemic | NE |
| *Cubanoptila cubana* | 100 | TRUE | 100 | 100 | TRUE | Endemic | NE |
| *Cubanoptila madremia* | 0 | FALSE | 95 | 95 | TRUE | Endemic | NE |
| *Cubanoptila muybonita* | 40 | TRUE | 91 | 97 | TRUE | Endemic | NE |
| *Cubanoptila purpurea* | 28 | FALSE | 30 | 38 | TRUE | Endemic | NE |
| *Cyprinodon variegatus* | 25 | FALSE | 50 | 52 | TRUE | Non-endemic | LC |
| *Cyrenoida americana* | 0 | FALSE | 92 | 92 | TRUE | Non-endemic | NE |
| *Dajaus monticola* | 0 | FALSE | 100 | 100 | TRUE | Non-endemic | LC |
| *Desmopachria andreae* | 34 | TRUE | 73 | 100 | TRUE | Endemic | NE |
| *Desmopachria tarda* | 83 | TRUE | 75 | 100 | TRUE | Endemic | NE |
| *Dicrotendipes simpsoni* | 100 | TRUE | 89 | 100 | TRUE | Non-endemic | NE |
| *Dineutus americanus* | 0 | FALSE | 96 | 96 | TRUE | Non-endemic | NE |
| *Dineutus longimanus* | 19 | FALSE | 36 | 41 | TRUE | Endemic | NE |
| *Dormitator maculatus* | 25 | FALSE | 33 | 42 | TRUE | Non-endemic | LC |
| *Drepanotrema aeruginosum* | 0 | FALSE | 100 | 100 | TRUE | Non-endemic | NE |
| *Drepanotrema anatinum* | 0 | FALSE | 31 | 31 | TRUE | Non-endemic | NE |
| *Drepanotrema cimex* | 0 | FALSE | 99 | 99 | TRUE | Non-endemic | LC |
| *Drepanotrema lucidum* | 38 | TRUE | 74 | 75 | TRUE | Non-endemic | NE |
| *Drosera capillaris* | 30 | TRUE | 33 | 34 | TRUE | Non-endemic | NE |
| *Dythemis rufinervis* | 33 | TRUE | 32 | 44 | TRUE | Non-endemic | LC |
| *Dythemis sterilis* | 32 | TRUE | 31 | 42 | TRUE | Non-endemic | LC |

| species | Relative protection SNAP | Met target SNAP | Relative protection free-choice | Relative protection lock-in | Met target solution | Endemic status | IUCN category |
|---|---|---|---|---|---|---|---|
| *Eleotris pisonis* | 37 | TRUE | 37 | 50 | TRUE | Non-endemic | LC |
| *Elodea densa* | 11 | FALSE | 62 | 39 | TRUE | Non-endemic | NE |
| *Elodes angustata* | 0 | FALSE | 100 | 100 | TRUE | Endemic | NE |
| *Enacantha caribbea* | 100 | TRUE | 100 | 100 | TRUE | Non-endemic | LC |
| *Enallagma civile* | 28 | FALSE | 30 | 42 | TRUE | Non-endemic | LC |
| *Enallagma coecum* | 11 | FALSE | 33 | 34 | TRUE | Non-endemic | LC |
| *Enochrus hamiltoni* | 74 | TRUE | 40 | 74 | TRUE | Non-endemic | NE |
| *Enochrus pygmaeus* | 0 | FALSE | 90 | 90 | TRUE | Non-endemic | NE |
| *Epilobocera capolongoi* | 98 | TRUE | 38 | 98 | TRUE | Endemic | DD |
| *Epilobocera cubensis* | 12 | FALSE | 32 | 35 | TRUE | Endemic | LC |
| *Epilobocera diazbeltrani* | 0 | FALSE | 100 | 100 | TRUE | Endemic | NE |
| *Epilobocera gilmanii* | 3 | FALSE | 57 | 60 | TRUE | Endemic | NE |
| *Epilobocera placensis* | 0 | FALSE | 80 | 80 | TRUE | Endemic | NE |
| *Epilobocera synoecia* | 5 | FALSE | 65 | 31 | TRUE | Endemic | NE |
| *Erythemis attala* | 0 | FALSE | 50 | 50 | TRUE | Non-endemic | LC |
| *Erythemis plebeja* | 42 | TRUE | 35 | 52 | TRUE | Non-endemic | LC |
| *Erythemis simplicicollis* | 44 | TRUE | 39 | 54 | TRUE | Non-endemic | LC |
| *Erythemis vesiculosa* | 42 | TRUE | 33 | 50 | TRUE | Non-endemic | LC |
| *Erythrodiplax berenice* | 59 | TRUE | 39 | 64 | TRUE | Non-endemic | LC |
| *Erythrodiplax bromeliicola* | 40 | TRUE | 38 | 40 | TRUE | Non-endemic | DD |
| *Erythrodiplax fervida* | 28 | FALSE | 35 | 44 | TRUE | Non-endemic | LC |
| *Erythrodiplax justiniana* | 32 | TRUE | 36 | 47 | TRUE | Non-endemic | LC |
| *Erythrodiplax umbrata* | 44 | TRUE | 35 | 53 | TRUE | Non-endemic | LC |
| *Eupera cubensis* | 54 | TRUE | 83 | 79 | TRUE | Endemic | LC |
| *Eurygerris cariniventris* | 100 | TRUE | 84 | 100 | TRUE | Non-endemic | NE |
| *Eustoma exaltatum* | 100 | TRUE | 65 | 100 | TRUE | Non-endemic | NE |
| *Fallceon longifolius* | 59 | TRUE | 59 | 69 | TRUE | Non-endemic | NE |
| *Fallceon poeyi* | 10 | FALSE | 30 | 30 | TRUE | Non-endemic | NE |
| *Fallceon sextus* | 100 | TRUE | 100 | 100 | TRUE | Endemic | NE |
| *Fallceon testudineus* | 100 | TRUE | 87 | 100 | TRUE | Endemic | NE |
| *Farrodes bimaculatus* | 13 | FALSE | 38 | 40 | TRUE | Endemic | NE |
| *Galba cubensis* | 21 | FALSE | 33 | 39 | TRUE | Non-endemic | LC |
| *Gambusia punctata* | 23 | FALSE | 30 | 38 | TRUE | Endemic | LC |

| species | Relative protection SNAP | Met target SNAP | Relative protection free-choice | Relative protection lock-in | Met target solution | Endemic status | IUCN category |
|---|---|---|---|---|---|---|---|
| *Gambusia puncticulata* | 37 | TRUE | 34 | 48 | TRUE | Non-endemic | LC |
| *Gambusia rhizophorae* | 0 | FALSE | 90 | 90 | TRUE | Non-endemic | LC |
| *Girardinus creolus* | 36 | TRUE | 66 | 66 | TRUE | Endemic | LC |
| *Girardinus denticulatus* | 4 | FALSE | 42 | 37 | TRUE | Endemic | LC |
| *Girardinus falcatus* | 72 | TRUE | 36 | 93 | TRUE | Endemic | LC |
| *Girardinus metallicus* | 28 | FALSE | 30 | 38 | TRUE | Endemic | LC |
| *Girardinus microdactylus* | 70 | TRUE | 47 | 100 | TRUE | Endemic | LC |
| *Girardinus uninonatus* | 62 | TRUE | 41 | 88 | TRUE | Endemic | NE |
| *Girardinus uninotatus* | 100 | TRUE | 99 | 100 | TRUE | Non-endemic | LC |
| *Gobiesox nudus* | 100 | TRUE | 100 | 100 | TRUE | Non-endemic | NE |
| *Goeldichironomus amazonicus* | 0 | FALSE | 80 | 80 | TRUE | Non-endemic | NE |
| *Goeldichironomus devineyae* | 0 | FALSE | 52 | 52 | TRUE | Non-endemic | NE |
| *Goeldichironomus holoprasinus* | 100 | TRUE | 95 | 100 | TRUE | Non-endemic | NE |
| *Goeldichironomus natans* | 100 | TRUE | 95 | 100 | TRUE | Non-endemic | NE |
| *Gundlachia radiata* | 45 | TRUE | 34 | 53 | TRUE | Non-endemic | NE |
| *Gymnochthebius fossatus* | 40 | TRUE | 69 | 87 | TRUE | Non-endemic | NE |
| *Gynacantha ereagris* | 0 | FALSE | 61 | 66 | TRUE | Non-endemic | LC |
| *Gynacantha nervosa* | 31 | TRUE | 72 | 84 | TRUE | Non-endemic | LC |
| *Gyraulus parvus* | 0 | FALSE | 100 | 100 | TRUE | Non-endemic | LC |
| *Hagenulus caligatus* | 21 | FALSE | 87 | 77 | TRUE | Endemic | NE |
| *Hagenulus morrisonae* | 20 | FALSE | 33 | 37 | TRUE | Endemic | NE |
| *Haliplus havaniensis* | 16 | FALSE | 75 | 91 | TRUE | Non-endemic | NE |
| *Haliplus tumidus* | 0 | FALSE | 84 | 84 | TRUE | Non-endemic | NE |
| *Helicopsyche comosa* | 14 | FALSE | 30 | 30 | TRUE | Endemic | NE |
| *Helicopsyche cubana* | 49 | TRUE | 91 | 100 | TRUE | Endemic | NE |
| *Helicopsyche falcigona* | 100 | TRUE | 100 | 100 | TRUE | Endemic | NE |
| *Helicopsyche hageni* | 40 | TRUE | 40 | 61 | TRUE | Non-endemic | NE |
| *Helicopsyche occidentalis* | 100 | TRUE | 100 | 100 | TRUE | Endemic | NE |
| *Hexacylloepus filiformis* | 0 | FALSE | 95 | 95 | TRUE | Endemic | NE |
| *Hydaticus bimarginatus* | 23 | FALSE | 98 | 100 | TRUE | Non-endemic | NE |
| *Hydaticus rimosus* | 0 | FALSE | 96 | 96 | TRUE | Non-endemic | NE |
| *Hydraena decui* | 74 | TRUE | 42 | 75 | TRUE | Endemic | NE |
| *Hydraena guadelupensis* | 47 | TRUE | 94 | 94 | TRUE | Non-endemic | NE |
| *Hydraena matthiasi* | 100 | TRUE | 93 | 100 | TRUE | Endemic | NE |

| species | Relative protection SNAP | Met target SNAP | Relative protection free-choice | Relative protection lock-in | Met target solution | Endemic status | IUCN category |
|---|---|---|---|---|---|---|---|
| *Hydraena perkinsi* | 22 | FALSE | 43 | 65 | TRUE | Endemic | NE |
| *Hydrocanthus oblongus* | 19 | FALSE | 91 | 93 | TRUE | Non-endemic | NE |
| *Hydrochus rugosus* | 0 | FALSE | 49 | 49 | TRUE | Non-endemic | NE |
| *Hydrometra australis* | 0 | FALSE | 37 | 37 | TRUE | Non-endemic | NE |
| *Hydrometra caraiba* | 0 | FALSE | 55 | 55 | TRUE | Non-endemic | NE |
| *Hydrometra gibara* | 0 | FALSE | 100 | 100 | TRUE | Endemic | NE |
| *Hydropsyche cubana* | 30 | TRUE | 49 | 58 | TRUE | Endemic | NE |
| *Hydropsyche dearmasi* | 0 | FALSE | 95 | 95 | TRUE | Endemic | NE |
| *Hydrovatus caraibus* | 29 | FALSE | 36 | 48 | TRUE | Non-endemic | NE |
| *Hydrovatus hornii* | 0 | FALSE | 100 | 100 | TRUE | Non-endemic | NE |
| *Hymenachne amplexicaulis* | 0 | FALSE | 100 | 100 | TRUE | Non-endemic | NE |
| *Hypolestes trinitatis* | 25 | FALSE | 43 | 48 | TRUE | Endemic | VU |
| *Idiataphe cubensis* | 2 | FALSE | 31 | 68 | TRUE | Non-endemic | LC |
| *Ischnura capreolus* | 38 | TRUE | 30 | 46 | TRUE | Non-endemic | LC |
| *Ischnura hastata* | 30 | TRUE | 42 | 67 | TRUE | Non-endemic | LC |
| *Ischnura ramburii* | 37 | TRUE | 33 | 47 | TRUE | Non-endemic | LC |
| *Jonga serrei* | 0 | FALSE | 95 | 95 | TRUE | Non-endemic | LC |
| *Joturus pichardi* | 25 | FALSE | 38 | 43 | TRUE | Non-endemic | LC |
| *Kryptolebias marmoratus* | 32 | TRUE | 61 | 94 | TRUE | Non-endemic | LC |
| *Labrundinia neopilosella* | 100 | TRUE | 89 | 100 | TRUE | Non-endemic | NE |
| *Laccobius antillensis* | 0 | FALSE | 100 | 100 | TRUE | Non-endemic | NE |
| *Laccomimus pumilio* | 0 | FALSE | 58 | 58 | TRUE | Non-endemic | NE |
| *Laccophilus bifasciatus* | 32 | TRUE | 38 | 51 | TRUE | Non-endemic | NE |
| *Laccophilus gentilis* | 25 | FALSE | 39 | 48 | TRUE | Non-endemic | NE |
| *Laccophilus proximus* | 30 | FALSE | 36 | 48 | TRUE | Non-endemic | NE |
| *Laccophilus vacaensis* | 2 | FALSE | 95 | 95 | TRUE | Non-endemic | NE |
| *Laccophilus venustus* | 27 | FALSE | 37 | 46 | TRUE | Non-endemic | NE |
| *Leptobasis vacillans* | 20 | FALSE | 71 | 71 | TRUE | Non-endemic | LC |
| *Leptonema poeyi* | 0 | FALSE | 37 | 37 | TRUE | Non-endemic | NE |
| *Lestes forficula* | 12 | FALSE | 57 | 69 | TRUE | Non-endemic | LC |
| *Lestes spumarius* | 0 | FALSE | 100 | 100 | TRUE | Non-endemic | LC |

| species | Relative protection SNAP | Met target SNAP | Relative protection free-choice | Relative protection lock-in | Met target solution | Endemic status | IUCN category |
|---|---|---|---|---|---|---|---|
| *Lestes tenuatus* | 24 | FALSE | 73 | 74 | TRUE | Non-endemic | LC |
| *Lethocerus colossicus* | 0 | FALSE | 90 | 90 | TRUE | Non-endemic | NE |
| *Libellula needhami* | 69 | TRUE | 98 | 98 | TRUE | Non-endemic | LC |
| *Limia vittata* | 31 | TRUE | 34 | 46 | TRUE | Endemic | LC |
| *Limnogonus franciscanus* | 26 | FALSE | 36 | 44 | TRUE | Non-endemic | NE |
| *Liodessus noviaffinis* | 0 | FALSE | 90 | 90 | TRUE | Non-endemic | NE |
| *Littoridinops monroensis* | 100 | TRUE | 100 | 100 | TRUE | Non-endemic | NE |
| *Lucifuga dentata* | 100 | TRUE | 53 | 100 | TRUE | Endemic | EN |
| *Ludwigia erecta* | 28 | FALSE | 49 | 85 | TRUE | Non-endemic | NE |
| *Ludwigia grandiflora* | 22 | FALSE | 78 | 100 | TRUE | Non-endemic | NE |
| *Ludwigia leptocarpa* | 32 | TRUE | 68 | 100 | TRUE | Non-endemic | LC |
| *Ludwigia peduncularis* | 0 | FALSE | 100 | 100 | TRUE | Non-endemic | NE |
| *Ludwigia peploides* | 5 | FALSE | 30 | 30 | TRUE | Non-endemic | NE |
| *Ludwigia peruviana* | 0 | FALSE | 69 | 69 | TRUE | Non-endemic | NE |
| *Ludwigia stricta* | 0 | FALSE | 100 | 100 | TRUE | Non-endemic | NE |
| *Lutrochus geniculatus* | 0 | FALSE | 91 | 91 | TRUE | Non-endemic | NE |
| *Macrobrachium acanthurus* | 30 | FALSE | 35 | 47 | TRUE | Non-endemic | LC |
| *Macrobrachium carcinus* | 0 | FALSE | 32 | 96 | TRUE | Non-endemic | LC |
| *Macrobrachium faustinum* | 30 | TRUE | 37 | 47 | TRUE | Non-endemic | LC |
| *Macrobrachium heterochirus* | 0 | FALSE | 100 | 100 | TRUE | Non-endemic | LC |
| *Macrodiplax balteata* | 30 | TRUE | 50 | 51 | TRUE | Non-endemic | LC |
| *Macronema tremenda* | 0 | FALSE | 93 | 93 | TRUE | Endemic | NE |
| *Macrothemis celeno* | 29 | FALSE | 30 | 40 | TRUE | Non-endemic | LC |
| *Marilia scudderi* | 12 | FALSE | 84 | 84 | TRUE | Endemic | NE |
| *Marilia wrighti* | 100 | TRUE | 100 | 100 | TRUE | Endemic | NE |
| *Marisa cornuarietis* | 0 | FALSE | 72 | 72 | TRUE | Non-endemic | LC |
| *Megadytes fraternus* | 100 | TRUE | 91 | 100 | TRUE | Non-endemic | NE |
| *Melanoides tuberculata* | 0 | FALSE | 45 | 40 | TRUE | Non-endemic | LC |
| *Meridiorhantus calidus* | 38 | TRUE | 36 | 51 | TRUE | Non-endemic | NE |
| *Merragata hebroides* | 0 | FALSE | 68 | 68 | TRUE | Non-endemic | NE |
| *Mesonoterus addendus* | 0 | FALSE | 100 | 100 | TRUE | Non-endemic | NE |

| species | Relative protection SNAP | Met target SNAP | Relative protection free-choice | Relative protection lock-in | Met target solution | Endemic status | IUCN category |
|---|---|---|---|---|---|---|---|
| *Mesoplocia inaccessibile* | 0 | FALSE | 75 | 75 | TRUE | Non-endemic | NE |
| *Mesovelia mulsanti* | 33 | TRUE | 33 | 45 | TRUE | Non-endemic | NE |
| *Metrichia munieca* | 0 | FALSE | 85 | 85 | TRUE | Endemic | NE |
| *Metrobates tumidus* | 11 | FALSE | 89 | 89 | TRUE | Non-endemic | NE |
| *Miathyria marcella* | 46 | TRUE | 37 | 56 | TRUE | Non-endemic | LC |
| *Miathyria simplex* | 0 | FALSE | 63 | 63 | TRUE | Non-endemic | LC |
| *Micrathyria aequalis* | 39 | TRUE | 31 | 48 | TRUE | Non-endemic | LC |
| *Micrathyria didyma* | 34 | TRUE | 31 | 34 | TRUE | Non-endemic | LC |
| *Micrathyria dissocians* | 0 | FALSE | 33 | 39 | TRUE | Non-endemic | LC |
| *Micrathyria hagenii* | 41 | TRUE | 42 | 46 | TRUE | Non-endemic | LC |
| *Micratya poeyi* | 5 | FALSE | 85 | 85 | TRUE | Non-endemic | LC |
| *Microvelia albonotata* | 0 | FALSE | 96 | 96 | TRUE | Non-endemic | NE |
| *Microvelia cubana* | 42 | TRUE | 36 | 52 | TRUE | Non-endemic | NE |
| *Microvelia paludicola* | 0 | FALSE | 100 | 100 | TRUE | Non-endemic | NE |
| *Mugil liza* | 0 | FALSE | 88 | 88 | TRUE | Non-endemic | DD |
| *Nandopsis ramsdeni* | 24 | FALSE | 33 | 35 | TRUE | Endemic | EN |
| *Nandopsis tetracanthus* | 24 | FALSE | 35 | 41 | TRUE | Endemic | LC |
| *Nectopsyche cubana* | 31 | TRUE | 44 | 55 | TRUE | Non-endemic | NE |
| *Nehalennia minuta* | 36 | TRUE | 50 | 86 | TRUE | Non-endemic | LC |
| *Nelumbo lutea* | 4 | FALSE | 95 | 99 | TRUE | Non-endemic | LC |
| *Neoepilobocera gertraudae* | 0 | FALSE | 100 | 100 | TRUE | Endemic | NE |
| *Neoerythromma cultellatum* | 100 | TRUE | 100 | 100 | TRUE | Non-endemic | LC |
| *Neoneura carnatica* | 0 | FALSE | 53 | 37 | TRUE | Endemic | DD |
| *Neoneura maria* | 12 | FALSE | 39 | 42 | TRUE | Endemic | DD |
| *Nereina punctulata* | 0 | FALSE | 70 | 70 | TRUE | Non-endemic | NE |
| *Nilothauma babiyi* | 0 | FALSE | 80 | 80 | TRUE | Non-endemic | NE |
| *Noelmis minima* | 0 | FALSE | 100 | 100 | TRUE | Endemic | NE |
| *Notomicrus sharpi* | 1 | FALSE | 98 | 98 | TRUE | Non-endemic | NE |
| *Notonecta indica* | 13 | FALSE | 49 | 49 | TRUE | Non-endemic | NE |
| *Nymphaea ampla* | 58 | TRUE | 35 | 81 | TRUE | Non-endemic | NE |
| *Nymphaea odorata* | 93 | TRUE | 55 | 93 | TRUE | Non-endemic | LC |
| *Ochrotrichia caramba* | 0 | FALSE | 96 | 96 | TRUE | Endemic | NE |

| species | Relative protection SNAP | Met target SNAP | Relative protection free-choice | Relative protection lock-in | Met target solution | Endemic status | IUCN category |
|---|---|---|---|---|---|---|---|
| *Ochthebius attritus* | 61 | TRUE | 35 | 61 | TRUE | Non-endemic | NE |
| *Ophisternon aenigmaticus* | 0 | FALSE | 53 | 45 | TRUE | Non-endemic | NE |
| *Orthemis discolor* | 21 | FALSE | 75 | 89 | TRUE | Non-endemic | LC |
| *Orthemis ferruginea* | 37 | TRUE | 35 | 49 | TRUE | Non-endemic | LC |
| *Pachychilus nigratus* | 0 | FALSE | 36 | 36 | TRUE | Endemic | NE |
| *Pachychilus violaceus* | 100 | TRUE | 48 | 100 | TRUE | Endemic | NE |
| *Pachydiplax longipennis* | 34 | TRUE | 34 | 68 | TRUE | Non-endemic | LC |
| *Pachydrus obniger* | 27 | FALSE | 36 | 47 | TRUE | Non-endemic | NE |
| *Palaemon pandaliformis* | 0 | FALSE | 74 | 74 | TRUE | Non-endemic | NE |
| *Paltostoma palominoi* | 0 | FALSE | 100 | 100 | TRUE | Non-endemic | NE |
| *Pantala flavescens* | 36 | TRUE | 34 | 48 | TRUE | Non-endemic | LC |
| *Pantala hymenaea* | 0 | FALSE | 71 | 71 | TRUE | Non-endemic | LC |
| *Paracymus londingi* | 0 | FALSE | 90 | 90 | TRUE | Non-endemic | NE |
| *Parakiefferiella coronata* | 0 | FALSE | 80 | 80 | TRUE | Non-endemic | NE |
| *Paraplea puella* | 0 | FALSE | 36 | 57 | TRUE | Non-endemic | NE |
| *Pelocoris poeyi* | 13 | FALSE | 34 | 38 | TRUE | Non-endemic | NE |
| *Pelonomus obscurus* | 35 | TRUE | 39 | 74 | TRUE | Non-endemic | NE |
| *Perithemis domitia* | 24 | FALSE | 35 | 42 | TRUE | Non-endemic | LC |
| *Phaenonotum exstriatum* | 0 | FALSE | 100 | 100 | TRUE | Non-endemic | NE |
| *Phylloicus chalybeus* | 31 | TRUE | 35 | 46 | TRUE | Endemic | NE |
| *Phylloicus cubanus* | 10 | FALSE | 99 | 99 | TRUE | Endemic | NE |
| *Physella acuta* | 33 | TRUE | 32 | 43 | TRUE | Non-endemic | LC |
| *Pisidium consanguineum* | 0 | FALSE | 93 | 93 | TRUE | Endemic | NE |
| *Planorbella duryi* | 47 | TRUE | 35 | 54 | TRUE | Non-endemic | NE |
| *Planorbella trivolvis* | 0 | FALSE | 100 | 100 | TRUE | Non-endemic | LC |
| *Platyvelia brachialis* | 100 | TRUE | 100 | 100 | TRUE | Non-endemic | NE |
| *Poecilophlebia pacoi* | 19 | FALSE | 72 | 74 | TRUE | Endemic | NE |
| *Polycentropus mathisi* | 14 | FALSE | 100 | 100 | TRUE | Endemic | NE |
| *Polycentropus turquino* | 100 | TRUE | 100 | 100 | TRUE | Endemic | NE |
| *Pomacea paludosa* | 28 | FALSE | 37 | 45 | TRUE | Non-endemic | LC |
| *Pomacea poeyana* | 26 | FALSE | 46 | 45 | TRUE | Endemic | NE |
| *Potimirin americana* | 0 | FALSE | 100 | 100 | TRUE | Non-endemic | NE |
| *Procambarus cubensis* | 23 | FALSE | 100 | 100 | TRUE | Endemic | DD |

| species | Relative protection SNAP | Met target SNAP | Relative protection free-choice | Relative protection lock-in | Met target solution | Endemic status | IUCN category |
|---|---|---|---|---|---|---|---|
| *Procambarus niveus* | 73 | TRUE | 73 | 73 | TRUE | Endemic | DD |
| *Progomphus integer* | 7 | FALSE | 38 | 38 | TRUE | Non-endemic | LC |
| *Protoneura caligata* | 21 | FALSE | 30 | 35 | TRUE | Endemic | EN |
| *Protoneura capillaris* | 23 | FALSE | 36 | 44 | TRUE | Endemic | DD |
| *Protoneura viridis* | 100 | TRUE | 68 | 100 | TRUE | Non-endemic | LC |
| *Pseudosuccinea columella* | 48 | TRUE | 38 | 57 | TRUE | Non-endemic | LC |
| *Psilopelmia hematopotum* | 32 | TRUE | 41 | 54 | TRUE | Non-endemic | NE |
| *Psilopelmia ochraceum* | 0 | FALSE | 85 | 85 | TRUE | Non-endemic | NE |
| *Pyrgophorus parvulus* | 0 | FALSE | 52 | 47 | TRUE | Non-endemic | LC |
| *Quintana atrizona* | 100 | TRUE | 100 | 100 | TRUE | Endemic | CR |
| *Ramphocorixa rotundocephala* | 0 | FALSE | 98 | 98 | TRUE | Non-endemic | NE |
| *Ranatra fabricii* | 0 | FALSE | 96 | 96 | TRUE | Endemic | NE |
| *Ranatra sagrai* | 0 | FALSE | 95 | 95 | TRUE | Endemic | NE |
| *Remartinia secreta* | 0 | FALSE | 93 | 93 | TRUE | Non-endemic | LC |
| *Rhagovelia collaris* | 21 | FALSE | 38 | 42 | TRUE | Non-endemic | NE |
| *Rhagovelia mira* | 0 | FALSE | 100 | 100 | TRUE | Endemic | NE |
| *Rheumatobates clanis* | 0 | FALSE | 100 | 65 | TRUE | Non-endemic | NE |
| *Rheumatobates meinerti* | 30 | TRUE | 30 | 30 | TRUE | Non-endemic | NE |
| *Rhionaeschna psilus* | 27 | FALSE | 37 | 47 | TRUE | Non-endemic | LC |
| *Rivulus cylindraceus* | 91 | TRUE | 91 | 91 | TRUE | Endemic | LC |
| *Sacciolepis striata* | 0 | FALSE | 100 | 100 | TRUE | Non-endemic | NE |
| *Sagittaria intermedia* | 0 | FALSE | 100 | 100 | TRUE | Non-endemic | NE |
| *Sagittaria lancifolia* | 87 | TRUE | 83 | 92 | TRUE | Non-endemic | NE |
| *Salvinia auriculata* | 44 | TRUE | 100 | 100 | TRUE | Non-endemic | LC |
| *Scapanea frontalis* | 26 | FALSE | 38 | 45 | TRUE | Non-endemic | LC |
| *Setaria geminata* | 0 | FALSE | 94 | 94 | TRUE | Non-endemic | NE |
| *Sicydium plumieri* | 32 | TRUE | 40 | 49 | TRUE | Non-endemic | DD |
| *Smicridea comma* | 8 | FALSE | 39 | 39 | TRUE | Non-endemic | NE |
| *Smicridea obesa* | 0 | FALSE | 95 | 95 | TRUE | Endemic | NE |
| *Steinovelia stagnalis* | 0 | FALSE | 96 | 96 | TRUE | Non-endemic | NE |
| *Suphis inflatus* | 0 | FALSE | 100 | 100 | TRUE | Non-endemic | NE |
| *Suphisellus insularis* | 0 | FALSE | 87 | 87 | TRUE | Non-endemic | NE |

| species | Relative protection SNAP | Met target SNAP | Relative protection free-choice | Relative protection lock-in | Met target solution | Endemic status | IUCN category |
|---|---|---|---|---|---|---|---|
| *Suphisellus nigrinus* | 24 | FALSE | 87 | 89 | TRUE | Non-endemic | NE |
| *Suphisellus tenuicornis* | 0 | FALSE | 100 | 100 | TRUE | Endemic | NE |
| *Sympetrum illotum* | 73 | TRUE | 87 | 90 | TRUE | Non-endemic | LC |
| *Tanytarsus limneticus* | 100 | TRUE | 89 | 100 | TRUE | Non-endemic | NE |
| *Tarebia granifera* | 22 | FALSE | 30 | 36 | TRUE | Non-endemic | LC |
| *Tauriphila australis* | 58 | TRUE | 97 | 97 | TRUE | Non-endemic | LC |
| *Telebasis dominicana* | 20 | FALSE | 35 | 40 | TRUE | Non-endemic | LC |
| *Thermonectus basillaris* | 28 | FALSE | 34 | 47 | TRUE | Non-endemic | NE |
| *Thermonectus circumscriptus* | 29 | FALSE | 36 | 47 | TRUE | Non-endemic | NE |
| *Thermonectus margineguttatus* | 0 | FALSE | 36 | 36 | TRUE | Non-endemic | NE |
| *Thermonectus succinctus* | 13 | FALSE | 39 | 44 | TRUE | Non-endemic | NE |
| *Tholymis citrina* | 44 | TRUE | 76 | 86 | TRUE | Non-endemic | LC |
| *Tramea abdominalis* | 38 | TRUE | 32 | 48 | TRUE | Non-endemic | LC |
| *Tramea calverti* | 0 | FALSE | 99 | 66 | TRUE | Non-endemic | LC |
| *Tramea insularis* | 58 | TRUE | 33 | 91 | TRUE | Non-endemic | LC |
| *Tramea onusta* | 42 | TRUE | 33 | 52 | TRUE | Non-endemic | LC |
| *Traverina cubensis* | 0 | FALSE | 100 | 100 | TRUE | Endemic | NE |
| *Traverina oriente* | 18 | FALSE | 35 | 38 | TRUE | Endemic | NE |
| *Trepobates taylori* | 0 | FALSE | 90 | 90 | TRUE | Non-endemic | NE |
| *Triacanthagyna septima* | 68 | TRUE | 91 | 100 | TRUE | Non-endemic | LC |
| *Triacanthagyna trifida* | 74 | TRUE | 97 | 74 | TRUE | Non-endemic | LC |
| *Tricorythodes cubensis* | 18 | FALSE | 43 | 46 | TRUE | Endemic | NE |
| *Tricorythodes grallator* | 25 | FALSE | 35 | 42 | TRUE | Endemic | NE |
| *Tricorythodes montanus* | 32 | TRUE | 41 | 52 | TRUE | Endemic | NE |
| *Tricorythodes sacculobranchis* | 13 | FALSE | 42 | 44 | TRUE | Endemic | NE |
| *Tricorythodes sierramaestrae* | 0 | FALSE | 100 | 47 | TRUE | Endemic | NE |
| *Tropisternus chalybeus* | 0 | FALSE | 96 | 96 | TRUE | Non-endemic | NE |
| *Tropisternus collaris* | 0 | FALSE | 96 | 96 | TRUE | Endemic | NE |
| *Tropisternus lateralis* | 0 | FALSE | 96 | 96 | TRUE | Endemic | NE |
| *Tropisternus mergus* | 0 | FALSE | 96 | 96 | TRUE | Endemic | NE |
| *Turquinophlebia grandis* | 0 | FALSE | 37 | 88 | TRUE | Endemic | NE |
| *Vitta usnea* | 100 | TRUE | 89 | 100 | TRUE | Non-endemic | NE |
| *Vitta virginea* | 72 | TRUE | 74 | 97 | TRUE | Non-endemic | LC |

| species | Relative protection SNAP | Met target SNAP | Relative protection free-choice | Relative protection lock-in | Met target solution | Endemic status | IUCN category |
|---|---|---|---|---|---|---|---|
| *Xiphocaris elongata* | 21 | FALSE | 36 | 43 | TRUE | Non-endemic | LC |
| *Xiphocaris gomezi* | 26 | FALSE | 36 | 82 | TRUE | Endemic | DD |
| *Xiphocentron cubanum* | 38 | TRUE | 89 | 96 | TRUE | Endemic | NE |
| *Xyris ambigua* | 77 | TRUE | 35 | 77 | TRUE | Non-endemic | NE |
| *Xyris grandiceps* | 0 | FALSE | 82 | 54 | TRUE | Non-endemic | NE |
| *Xyris jupicai* | 100 | TRUE | 100 | 100 | TRUE | Non-endemic | NE |
| *Xyris navicularis* | 49 | TRUE | 49 | 49 | TRUE | Non-endemic | NE |

Table S1.5. Pairwise Spearman correlations of planning-unit importance among taxonomic groups. Correlations were calculated separately for the free-choice and lock-in prioritization scenarios using planning units with positive irreplaceability for at least one of the two compared groups. Positive values indicate spatial congruence in taxon-specific planning-unit importance, whereas negative values indicate spatial mismatch.

| Scenario | Taxonomic group 1 | Taxonomic group 2 | Spearman's rho | Correlation strength | P-value | Number of planning units |
|---|---|---|---|---|---|---|
| free-choice | Ephemeroptera | Trichoptera | 0.873 | Strong positive | <0.001 | 6108 |
| free-choice | Coleoptera | Hemiptera | 0.842 | Strong positive | <0.001 | 6987 |
| free-choice | Mollusca | Odonata | 0.835 | Strong positive | <0.001 | 6989 |
| free-choice | Decapoda | Hemiptera | 0.753 | Strong positive | <0.001 | 6897 |
| free-choice | Coleoptera | Decapoda | 0.738 | Strong positive | <0.001 | 6986 |
| free-choice | Odonata | Vertebrates | 0.737 | Strong positive | <0.001 | 6989 |
| free-choice | Ephemeroptera | Hemiptera | 0.718 | Strong positive | <0.001 | 6912 |
| free-choice | Coleoptera | Odonata | 0.678 | Moderate positive | <0.001 | 6989 |
| free-choice | Mollusca | Vertebrates | 0.676 | Moderate positive | <0.001 | 6943 |
| free-choice | Coleoptera | Ephemeroptera | 0.663 | Moderate positive | <0.001 | 6986 |
| free-choice | Diptera | Trichoptera | 0.66 | Moderate positive | <0.001 | 5169 |
| free-choice | Diptera | Ephemeroptera | 0.658 | Moderate positive | <0.001 | 6023 |
| free-choice | Decapoda | Vertebrates | 0.637 | Moderate positive | <0.001 | 6872 |
| free-choice | Coleoptera | Vertebrates | 0.604 | Moderate positive | <0.001 | 6989 |
| free-choice | Hemiptera | Odonata | 0.599 | Moderate positive | <0.001 | 6989 |
| free-choice | Hemiptera | Vertebrates | 0.596 | Moderate positive | <0.001 | 6951 |
| free-choice | Hemiptera | Trichoptera | 0.538 | Moderate positive | <0.001 | 6891 |
| free-choice | Decapoda | Odonata | 0.53 | Moderate positive | <0.001 | 6989 |
| free-choice | Coleoptera | Trichoptera | 0.518 | Moderate positive | <0.001 | 6986 |
| free-choice | Decapoda | Ephemeroptera | 0.484 | Moderate positive | <0.001 | 6663 |
| free-choice | Coleoptera | Diptera | 0.422 | Moderate positive | <0.001 | 6986 |
| free-choice | Diptera | Macrophytes | -0.422 | Moderate negative | <0.001 | 5604 |
| free-choice | Decapoda | Mollusca | 0.372 | Weak positive | <0.001 | 6941 |
| free-choice | Diptera | Hemiptera | 0.358 | Weak positive | <0.001 | 6853 |
| free-choice | Ephemeroptera | Odonata | 0.355 | Weak positive | <0.001 | 6989 |
| free-choice | Coleoptera | Mollusca | 0.342 | Weak positive | <0.001 | 6989 |
| free-choice | Macrophytes | Mollusca | 0.306 | Weak positive | <0.001 | 6908 |
| free-choice | Hemiptera | Mollusca | 0.29 | Weak positive | <0.001 | 6980 |
| free-choice | Hemiptera | Macrophytes | 0.248 | Weak positive | <0.001 | 6901 |
| free-choice | Decapoda | Macrophytes | 0.246 | Weak positive | <0.001 | 6652 |
| free-choice | Ephemeroptera | Vertebrates | 0.235 | Weak positive | <0.001 | 6981 |
| free-choice | Ephemeroptera | Macrophytes | 0.231 | Weak positive | <0.001 | 6429 |
| free-choice | Odonata | Trichoptera | 0.229 | Weak positive | <0.001 | 6989 |
| free-choice | Decapoda | Trichoptera | 0.226 | Weak positive | <0.001 | 6523 |
| free-choice | Macrophytes | Odonata | 0.22 | Weak positive | <0.001 | 6989 |
| free-choice | Diptera | Mollusca | -0.206 | Weak negative | <0.001 | 6946 |

| Scenario | Taxonomic group 1 | Taxonomic group 2 | Spearman's rho | Correlation strength | P-value | Number of planning units |
|---|---|---|---|---|---|---|
| free-choice | Diptera | Odonata | 0.168 | Weak positive | <0.001 | 6989 |
| free-choice | Macrophytes | Trichoptera | 0.111 | Weak positive | <0.001 | 5887 |
| free-choice | Mollusca | Trichoptera | -0.103 | Weak negative | <0.001 | 6948 |
| free-choice | Coleoptera | Macrophytes | 0.082 | Little or no correlation | <0.001 | 6987 |
| free-choice | Diptera | Vertebrates | -0.045 | Little or no correlation | <0.001 | 6919 |
| free-choice | Trichoptera | Vertebrates | 0.043 | Little or no correlation | <0.001 | 6974 |
| free-choice | Macrophytes | Vertebrates | 0.032 | Little or no correlation | <0.01 | 6857 |
| free-choice | Decapoda | Diptera | 0.015 | Little or no correlation | n.s. | 6463 |
| free-choice | Ephemeroptera | Mollusca | 0 | Little or no correlation | n.s. | 6966 |
| lock-in | Coleoptera | Hemiptera | 0.831 | Strong positive | <0.001 | 10683 |
| lock-in | Mollusca | Odonata | 0.829 | Strong positive | <0.001 | 10685 |
| lock-in | Ephemeroptera | Trichoptera | 0.821 | Strong positive | <0.001 | 9408 |
| lock-in | Decapoda | Hemiptera | 0.729 | Strong positive | <0.001 | 10572 |
| lock-in | Coleoptera | Decapoda | 0.699 | Moderate positive | <0.001 | 10682 |
| lock-in | Diptera | Ephemeroptera | 0.69 | Moderate positive | <0.001 | 9285 |
| lock-in | Odonata | Vertebrates | 0.689 | Moderate positive | <0.001 | 10685 |
| lock-in | Coleoptera | Odonata | 0.669 | Moderate positive | <0.001 | 10685 |
| lock-in | Ephemeroptera | Hemiptera | 0.642 | Moderate positive | <0.001 | 10583 |
| lock-in | Decapoda | Vertebrates | 0.638 | Moderate positive | <0.001 | 10570 |
| lock-in | Diptera | Trichoptera | 0.634 | Moderate positive | <0.001 | 7557 |
| lock-in | Mollusca | Vertebrates | 0.614 | Moderate positive | <0.001 | 10642 |
| lock-in | Coleoptera | Vertebrates | 0.605 | Moderate positive | <0.001 | 10685 |
| lock-in | Hemiptera | Vertebrates | 0.586 | Moderate positive | <0.001 | 10645 |
| lock-in | Coleoptera | Ephemeroptera | 0.577 | Moderate positive | <0.001 | 10682 |
| lock-in | Hemiptera | Odonata | 0.563 | Moderate positive | <0.001 | 10685 |
| lock-in | Diptera | Macrophytes | -0.506 | Moderate negative | <0.001 | 7944 |
| lock-in | Decapoda | Odonata | 0.488 | Moderate positive | <0.001 | 10685 |
| lock-in | Hemiptera | Trichoptera | 0.383 | Weak positive | <0.001 | 10550 |
| lock-in | Coleoptera | Trichoptera | 0.376 | Weak positive | <0.001 | 10682 |
| lock-in | Coleoptera | Diptera | 0.372 | Weak positive | <0.001 | 10682 |
| lock-in | Decapoda | Mollusca | 0.35 | Weak positive | <0.001 | 10632 |
| lock-in | Decapoda | Ephemeroptera | 0.33 | Weak positive | <0.001 | 10203 |
| lock-in | Diptera | Hemiptera | 0.33 | Weak positive | <0.001 | 10501 |
| lock-in | Coleoptera | Mollusca | 0.324 | Weak positive | <0.001 | 10685 |
| lock-in | Diptera | Mollusca | -0.299 | Weak negative | <0.001 | 10642 |
| lock-in | Macrophytes | Mollusca | 0.277 | Weak positive | <0.001 | 10598 |
| lock-in | Hemiptera | Mollusca | 0.247 | Weak positive | <0.001 | 10676 |
| lock-in | Mollusca | Trichoptera | -0.21 | Weak negative | <0.001 | 10644 |
| lock-in | Ephemeroptera | Odonata | 0.209 | Weak positive | <0.001 | 10685 |
| lock-in | Macrophytes | Odonata | 0.158 | Weak positive | <0.001 | 10685 |
| lock-in | Ephemeroptera | Vertebrates | 0.154 | Weak positive | <0.001 | 10676 |
| lock-in | Ephemeroptera | Macrophytes | 0.136 | Weak positive | <0.001 | 9878 |

| Scenario | Taxonomic group 1 | Taxonomic group 2 | Spearman's rho | Correlation strength | P-value | Number of planning units |
|---|---|---|---|---|---|---|
| lock-in | Ephemeroptera | Mollusca | -0.134 | Weak negative | <0.001 | 10662 |
| lock-in | Macrophytes | Vertebrates | -0.12 | Weak negative | <0.001 | 10560 |
| lock-in | Trichoptera | Vertebrates | -0.088 | Little or no correlation | <0.001 | 10668 |
| lock-in | Hemiptera | Macrophytes | 0.087 | Little or no correlation | <0.001 | 10560 |
| lock-in | Odonata | Trichoptera | 0.082 | Little or no correlation | <0.001 | 10685 |
| lock-in | Decapoda | Diptera | -0.082 | Little or no correlation | <0.001 | 9900 |
| lock-in | Diptera | Odonata | 0.064 | Little or no correlation | <0.001 | 10685 |
| lock-in | Diptera | Vertebrates | -0.06 | Little or no correlation | <0.001 | 10612 |
| lock-in | Decapoda | Macrophytes | 0.054 | Little or no correlation | <0.001 | 10069 |
| lock-in | Coleoptera | Macrophytes | -0.054 | Little or no correlation | <0.001 | 10683 |
| lock-in | Macrophytes | Trichoptera | 0.01 | Little or no correlation | n.s. | 8430 |
| lock-in | Decapoda | Trichoptera | 0.007 | Little or no correlation | n.s. | 9987 |

Table S1.6. Pairwise Jaccard similarity among taxon-specific priority sets. Jaccard similarity was calculated between pairs of taxonomic groups for each prioritization scenario using planning units in the upper 5% of positive group-specific irreplaceability values. Higher values indicate greater overlap between the highest-importance planning units of two taxonomic groups, whereas lower values indicate spatial mismatch. The Jaccard class provides a categorical interpretation of overlap strength.

| scenario | group_1 | group_2 | Jaccard similarity | Jaccard class |
|---|---|---|---|---|
| free-choice | Decapoda | Hemiptera | 0.576 | High overlap |
| free-choice | Decapoda | Vertebrates | 0.564 | High overlap |
| free-choice | Hemiptera | Vertebrates | 0.483 | Moderate overlap |
| free-choice | Coleoptera | Hemiptera | 0.439 | Moderate overlap |
| free-choice | Coleoptera | Decapoda | 0.409 | Moderate overlap |
| free-choice | Ephemeroptera | Trichoptera | 0.404 | Moderate overlap |
| free-choice | Coleoptera | Odonata | 0.394 | Moderate overlap |
| free-choice | Diptera | Trichoptera | 0.388 | Moderate overlap |
| free-choice | Coleoptera | Vertebrates | 0.342 | Moderate overlap |
| free-choice | Mollusca | Odonata | 0.287 | Low overlap |
| free-choice | Odonata | Vertebrates | 0.287 | Low overlap |
| free-choice | Hemiptera | Odonata | 0.284 | Low overlap |
| free-choice | Decapoda | Odonata | 0.268 | Low overlap |
| free-choice | Diptera | Ephemeroptera | 0.223 | Low overlap |
| free-choice | Macrophytes | Mollusca | 0.196 | Low overlap |
| free-choice | Coleoptera | Ephemeroptera | 0.154 | Low overlap |
| free-choice | Ephemeroptera | Odonata | 0.13 | Low overlap |
| free-choice | Coleoptera | Mollusca | 0.121 | Low overlap |
| free-choice | Ephemeroptera | Hemiptera | 0.12 | Low overlap |
| free-choice | Macrophytes | Odonata | 0.109 | Low overlap |
| free-choice | Mollusca | Vertebrates | 0.109 | Low overlap |
| free-choice | Decapoda | Mollusca | 0.099 | Very low overlap |
| free-choice | Decapoda | Ephemeroptera | 0.096 | Very low overlap |
| free-choice | Ephemeroptera | Vertebrates | 0.085 | Very low overlap |
| free-choice | Coleoptera | Trichoptera | 0.085 | Very low overlap |
| free-choice | Hemiptera | Mollusca | 0.084 | Very low overlap |
| free-choice | Ephemeroptera | Macrophytes | 0.082 | Very low overlap |
| free-choice | Coleoptera | Macrophytes | 0.078 | Very low overlap |
| free-choice | Coleoptera | Diptera | 0.078 | Very low overlap |
| free-choice | Decapoda | Macrophytes | 0.071 | Very low overlap |
| free-choice | Hemiptera | Macrophytes | 0.069 | Very low overlap |
| free-choice | Macrophytes | Vertebrates | 0.06 | Very low overlap |
| free-choice | Ephemeroptera | Mollusca | 0.047 | Very low overlap |
| free-choice | Odonata | Trichoptera | 0.045 | Very low overlap |
| free-choice | Hemiptera | Trichoptera | 0.037 | Very low overlap |
| free-choice | Decapoda | Trichoptera | 0.033 | Very low overlap |

| scenario | group_1 | group_2 | Jaccard similarity | Jaccard class |
|---|---|---|---|---|
| free-choice | Macrophytes | Trichoptera | 0.028 | Very low overlap |
| free-choice | Trichoptera | Vertebrates | 0.024 | Very low overlap |
| free-choice | Diptera | Odonata | 0.023 | Very low overlap |
| free-choice | Mollusca | Trichoptera | 0.019 | Very low overlap |
| free-choice | Diptera | Vertebrates | 0.009 | Very low overlap |
| free-choice | Diptera | Hemiptera | 0.009 | Very low overlap |
| free-choice | Diptera | Macrophytes | 0.006 | Very low overlap |
| free-choice | Decapoda | Diptera | 0.005 | Very low overlap |
| free-choice | Diptera | Mollusca | 0.004 | Very low overlap |
| lock-in | Decapoda | Vertebrates | 0.569 | High overlap |
| lock-in | Decapoda | Hemiptera | 0.548 | High overlap |
| lock-in | Hemiptera | Vertebrates | 0.455 | Moderate overlap |
| lock-in | Ephemeroptera | Trichoptera | 0.439 | Moderate overlap |
| lock-in | Coleoptera | Hemiptera | 0.412 | Moderate overlap |
| lock-in | Diptera | Trichoptera | 0.407 | Moderate overlap |
| lock-in | Coleoptera | Odonata | 0.388 | Moderate overlap |
| lock-in | Coleoptera | Decapoda | 0.385 | Moderate overlap |
| lock-in | Mollusca | Odonata | 0.323 | Moderate overlap |
| lock-in | Coleoptera | Vertebrates | 0.311 | Moderate overlap |
| lock-in | Odonata | Vertebrates | 0.289 | Low overlap |
| lock-in | Hemiptera | Odonata | 0.274 | Low overlap |
| lock-in | Decapoda | Odonata | 0.263 | Low overlap |
| lock-in | Diptera | Ephemeroptera | 0.262 | Low overlap |
| lock-in | Coleoptera | Ephemeroptera | 0.172 | Low overlap |
| lock-in | Macrophytes | Mollusca | 0.166 | Low overlap |
| lock-in | Ephemeroptera | Hemiptera | 0.139 | Low overlap |
| lock-in | Ephemeroptera | Odonata | 0.136 | Low overlap |
| lock-in | Mollusca | Vertebrates | 0.12 | Low overlap |
| lock-in | Coleoptera | Mollusca | 0.119 | Low overlap |
| lock-in | Macrophytes | Odonata | 0.116 | Low overlap |
| lock-in | Ephemeroptera | Macrophytes | 0.113 | Low overlap |
| lock-in | Decapoda | Ephemeroptera | 0.106 | Low overlap |
| lock-in | Coleoptera | Macrophytes | 0.1 | Very low overlap |
| lock-in | Hemiptera | Macrophytes | 0.098 | Very low overlap |
| lock-in | Ephemeroptera | Vertebrates | 0.097 | Very low overlap |
| lock-in | Coleoptera | Diptera | 0.094 | Very low overlap |
| lock-in | Decapoda | Macrophytes | 0.093 | Very low overlap |
| lock-in | Decapoda | Mollusca | 0.093 | Very low overlap |
| lock-in | Coleoptera | Trichoptera | 0.089 | Very low overlap |
| lock-in | Hemiptera | Mollusca | 0.081 | Very low overlap |
| lock-in | Macrophytes | Vertebrates | 0.072 | Very low overlap |
| lock-in | Ephemeroptera | Mollusca | 0.057 | Very low overlap |

| scenario | group_1 | group_2 | Jaccard similarity | Jaccard class |
|---|---|---|---|---|
| lock-in | Odonata | Trichoptera | 0.046 | Very low overlap |
| lock-in | Hemiptera | Trichoptera | 0.044 | Very low overlap |
| lock-in | Decapoda | Trichoptera | 0.03 | Very low overlap |
| lock-in | Macrophytes | Trichoptera | 0.027 | Very low overlap |
| lock-in | Trichoptera | Vertebrates | 0.023 | Very low overlap |
| lock-in | Diptera | Odonata | 0.021 | Very low overlap |
| lock-in | Mollusca | Trichoptera | 0.02 | Very low overlap |
| lock-in | Diptera | Hemiptera | 0.016 | Very low overlap |
| lock-in | Diptera | Vertebrates | 0.007 | Very low overlap |
| lock-in | Decapoda | Diptera | 0.005 | Very low overlap |
| lock-in | Diptera | Macrophytes | 0.004 | Very low overlap |
| lock-in | Diptera | Mollusca | 0.003 | Very low overlap |
